# AtomWeaver: Multi-Component Flow Matching with a Structured Geometric Prior Facilitates Non-Canonical Peptide Design

**DOI:** 10.64898/2026.09.23.753608

**Authors:** Alexander Kitaygorodsky, David Earl Hostallero, Aron Broom, Elliot Layne, Sungwon Hwang, Ashlin K. Kanawaty, Tomáš Babej, Glenn L. Butterfoss, Mark Fingerhuth

## Abstract

Fixed-backbone sequence discovery, or *inverse folding*, is a critical recurring task in the development of new polypeptide therapeutics. Once promising backbones are established for a target pocket, computational inverse folding methods greatly help accelerate generation of candidate sequences. Such methods are mature for the traditional case of limiting to the fixed twenty-letter *canonical* vocabulary; however, they cannot access the broader space of non-canonical amino acids (NCAAs). This design constraint is exacerbated for peptide binders, a fast-growing modality that readily incorporates NCAAs, though in practice non-canonical design frequently depends on laborious medicinal-chemistry campaigns. An extension of inverse folding to NCAAs is thus critical to accelerating design of novel therapeutic peptides.

*AtomWeaver* uses a joint all-site, atom-level generative scheme that does not restrict side-chain categorical assignment by either predetermined or co-resolving residue identity. Conditioned only on a fixed peptide backbone and its target protein, its multi-component flow guides side-chain atoms as unlabeled points in R^3^ from a *nested shell* prior to a variable-count final atom cloud. Identity is then read by matching each predicted cloud against a reference library of canonical and non-canonical templates. Since identity is decided only at decode time, the addressable vocabulary is a property of the library rather than of the trained weights: a new NCAA costs one reference structure and no retraining, and the model can select residues it was never prompted for and never saw in training.

On a deep mutational scan of two peptide–target systems, AtomWeaver’s canonical readout shows high observed mean agreement with experimental values among the compared inverse-folding methods. In the mixed canonical–noncanonical setting that canonical-only baselines cannot support at all, it likewise retains ranking signal across both systems. AtomWeaver also displayed self-consistent designs on *de novo* binder backbones, with the highest interface confidence among compared methods. Notably, it reached these metrics *while* achieving broad empirical coverage of our 300-residue vocabulary, including four non-canonical types never visible in training. AtomWeaver thus serves canonical and non-canonical peptide design alike, while transforming residue vocabulary to an expandable inference-time choice.

## 1. Introduction

Peptide therapeutics are a fast-growing modality, combining the specificity of biologics with the manufacturability of small molecules [1]. Much of that advantage comes from the freedom to place a non-canonical amino acid (NCAA) at any position. NCAAs are residues beyond the canonical twenty, differing from canonicals in side-chain, backbone pattern, or stereochemistry. They greatly expand the available chemical space and can confer protease resistance, improve membrane permeability and tune pharmacokinetics [2]. Over a thousand variants are commercially available [3], so a method restricted to the canonical twenty reaches a fraction of the chemistry peptide synthesis allows. Non-canonical substitutions are conventionally proposed by structure-guided inspection and physics-based modeling with expanded rotamer libraries [4–6]. While valuable, such representations scale poorly and struggle to keep pace with ML-driven advances. Extending learned inverse folding to non-canonical chemistry would close that gap, broaden where these methods apply, and accelerate the development of new peptide therapeutics.

Machine learning-based inverse folding, proposing sequences that fold into an intended structure given a fixed backbone, has matured rapidly for canonical design. Traditionally, identity is predicted over the fixed twenty types, from which atom coordinates can then be modeled under tight geometric constraint. ProteinMPNN [7] and its descendants LigandMPNN [8], PeptideMPNN [9] and FAMPNN [10] predict that identity as a categorical distribution; of these, FAMPNN additionally produces side-chain coordinates, while the others output sequence alone. Throughout this work we focus on the target-conditioned setting, where the binder is designed against a specified receptor.

Trained on the PDB, these models inherit the statistics of its primarily wild-type (WT) protein corpus. Thus, the sequences they favor are the ones selection has already explored at similar environments. Although these methods achieve high WT recovery rates and have produced many experimentally validated designs, this strong prior can be a constraint on more exotic sequence design. The harder limit is the restricted vocabulary: a residue absent from it cannot be proposed no matter its complementarity to the backbone and target pocket. Furthermore, vocabulary expansion requires investment in costly model retraining, and is not possible *at all* without significant per-residue structural data. For example, RareFold [11] treats each of 20 canonical and 29 non-canonical types as a separate trained token.

Latent-space co-generative models loosen the ordering of identity into conditioned atom generation, introducing *co*-mediation of identity through a learned per-residue embedding of identity and geometry together. This can expand the discoverable vocabulary via a variational approach to interpolate space in the latent manifold; nevertheless, even such methods can struggle to reach unconventional chemistries beyond what their embeddings were robustly trained to represent. La-Proteina [12] flows C*α* coordinates alongside a per-residue variational autoencoder (VAE) embedding decoded to Atom37, with its binder extension Proteina-Complexa [13] adding target conditioning; PepGLAD [14] applies multi-modal diffusion in the latent-space; AnewOmni [15] runs diffusion over a block-level latent embedding, reaching non-canonicals when the user supplies the NCAA’s chemical graph as a prompt. NCFlow [16] scores user-enumerated candidate residues at a fixed peptide site by generating their conformations in the peptide–receptor environment and applying a separate affinity-scoring workflow [17]; it therefore does not define a coupled, site-wise distribution over a peptide sequence. UNAAGI [18] instead generates an atom-level substitution for a fixed local backbone environment but samples only one local side chain at a time. Both are important atom-level precedents, but neither performs whole-peptide inverse folding.

A model that generates atoms rather than labels meets a further representational obstacle: the per-residue atom count is variable, and must be inferred jointly with atom coordinates and identity. Existing approaches use latent compression [12], padding to a fixed budget with per-atom occupancy [19–21], or sampling the atom count from a learned categorical distribution before generating coordinates [22]. *AtomWeaver* ‘s unique solution to this problem is described below in §3.

Throughout our work, we focus primarily on side-chain-modified NCAAs and D-stereochemistry, as the classes most congruent with a fixed backbone. We nonetheless include coverage of *β*-types and other backbone-modifying NCAAs, and see notable recovery even on such exotic backbones.

## 2. Background

### Flow matching with a non-Gaussian source

AtomWeaver’s coordinate track is trained as a continuous normalizing flow under the conditional flow-matching (CFM) framework [23]. Given a base distribution *p*_*T*_ on ℝ ^*d*^ and a target distribution *p*_0_, CFM learns a velocity field ***v***_*θ*_(***x***, *t*) whose probability-flow ODE *d****x****/dt* = ***v***_*θ*_(***x***, *t*) transports *p*_*T*_ to *p*_0_. The marginal flow is intractable, so training uses the *per-sample* conditional vector fields admitted in closed form by the interpolant path ***x***_*t*_ = (1 *− s*(*t*)) ***x***_0_ + *s*(*t*) ***x***_*T*_:

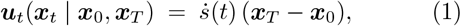

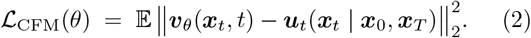

By the marginal–conditional equivalence of Lipman et al. [23], this objective is well-defined and yields the correct marginal velocity field for any base distribution *p*_*T*_. In place of the default Gaussian *p*_*T*_ = *N*(0, *I*_*d*_), we can thus substitute our own custom nested shell prior without disruption of theorem correctness.

#### Masked discrete diffusion

The element track is a masked-diffusion process in the style of D3PM [25] and SEDD [26]. For a finite vocabulary *V* containing a designated MASK token, the forward process noises a clean atom array ***e***_0_ toward an all-MASK end-state via class-weighted categorical transitions; the reverse process predicts *p*_*θ*_(***e***_0_ | ***e***_*t*_, ctx) at each step and samples ***e***_*t−*1_ from the corresponding posterior. We use the *absorbing* variant: a resolved slot cannot return to MASK, implemented by zeroing the MASK-class posterior probability for resolved slots and renormalizing.

## 3. AtomWeaver: framework

### Two coupled tracks

For a binder of length *L* and per-residue atom budget *K*, AtomWeaver jointly models side-chain positions ***x*** ∈ R^*L×K×*3^ and per-slot element types ***e* ∈** {GHOST, C, N, O, X}^*L×K*^ , where X is any element other than carbon, nitrogen or oxygen (sulfur, phosphorus, halogen, etc.). The element vocabulary includes a distinguished GHOST symbol for an absent atom, so a slot’s existence value is read directly from the element track and no separate existence track is modeled. The coordinate track is trained by conditional flow matching [23], with its velocity predicted by an SE(3)-equivariant denoiser [27]; the element track is a discrete absorbing masked-diffusion process [25, 26] coupled to the coordinate track on a shared timescale *t* ∈ [0, *T* = 250].

Side-chain generation is a two-component mixture: GHOST slots flow to C*α* and REAL^1^ slots to the ground-truth coordinates at the *t* = 0 endpoint, decomposed by FiLM-modulating [28] the shared per-slot representation with each slot’s GHOST/REAL state at every reverse step. The count of active atoms is thereby resolved jointly with conformation and type, following mixture modeling for diffusion and flow-matching generators [29– 31]. The receptor enters the context through a separate FiLM pathway, cross-attention, and direct participation of receptor atoms in the *equivariant graph* — the sparse radius graph over binder and target atoms on which the SE(3) denoiser attends, so that a receptor atom is a node the flow can see rather than a summary vector it is handed.

### The *nested shell* source distribution

Rather than initializing every atom from the same isotropic Gaussian cloud, AtomWeaver starts each atom close to where the training data reflects that atom should belong. Each is drawn from a shell at its characteristic radius along a shared backbone-derived pseudo-C*β* [24] ray (Fig. 1A,B):

**Figure 1:**
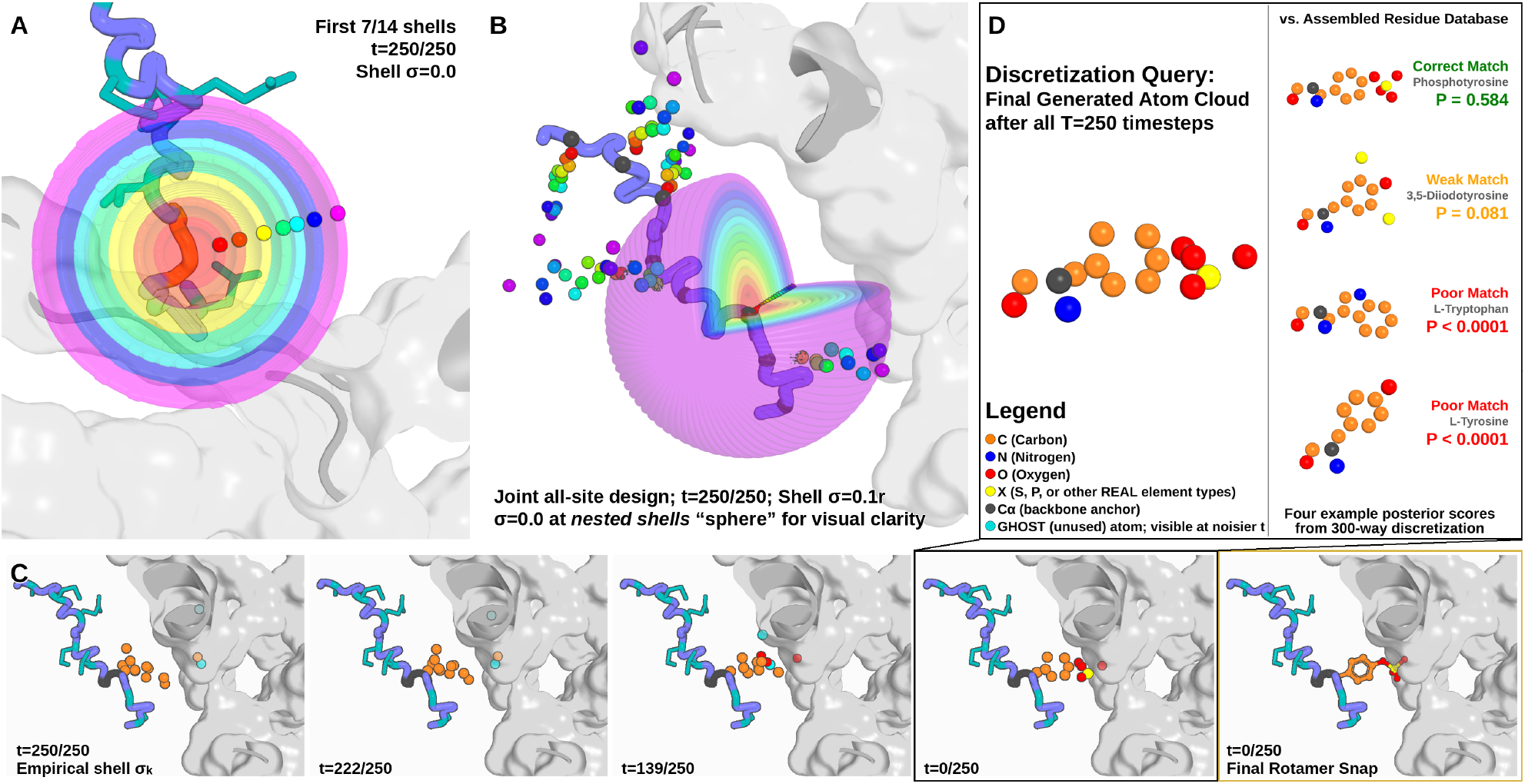
AtomWeaver packs side-chain atoms coordinates-first, then reads off identity. **(A)** Source distribution for one side-chain: atoms are drawn ***x***_*T*_ = C*α* + *r*_*k*_ ***d*** + *σ*_*k*_ ***η, η*** ∼ ***N***(0, *I*_3_), from *nested shells*, each at its characteristic radius *r*_*k*_ along a shared backbone-derived pseudo-C*β* [24] ray ***d***, rather than from an isotropic Gaussian; cross-section of the first *K* = 7 shells at *σ* = 0, giving no shell overlap or jitter. **(B)** The prior initialized jointly across all sites of a peptide backbone at the target interface at *σ* = 0.1 *r*, with jitter visible, and panel (A)’s nested sphere repeated at *K* = 14 in another view. **(C)** Reverse-sampling trajectory under the coupled flow: from empirical-*σ*_*k*_ starting points, a two-component mixture sends GHOST atoms (cyan) to C*α* and REAL atoms to refined side-chain coordinates, the per-slot GHOST/REAL state modulating the coordinate velocity at each step; the final frame is the packed side-chain after discretization. **(D)** At the penultimate step of (C), the predicted cloud is ranked against the reference library by the decode-time discretizer (§S5.3), which blends two scores per candidate class: a distance-matrix correlation, penalized for atom-count and element mismatches, and a learned classifier over the 300 residue types. The top match is phosphotyrosine, the residue realized at the end of (C), at posterior probability 0.58; the next highest falls below 0.09. Panels (A)–(B) set the jitter to a fixed fraction of each shell’s radius so that the shells read as a nested family at a glance; the trajectory in (C), and the model throughout, uses the empirical per-slot *σ*_*k*_ of Eq. 3, tabulated in Supplementary Table S1.

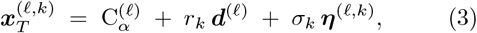

with isotropic noise ***η***^(*l,k*)^ ∼ *N*(0, *I*_3_), per-slot shell radius *r*_*k*_ along the residue-shared direction ***d***^(*l*)^, jittered by the per-slot shell thickness *σ*_*k*_. Both are *empirical*: *r*_*k*_ is the RMS distance of slot *k* from C*α* over the training data, and 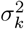 the corresponding per-slot radial variance Var(∥***x***_0_ ™ C*α*∥). At *σ*_*k*_ = 0 atoms are pinned to the ray; larger *σ*_*k*_ spreads them into overlapping shells. At *inference* we scale the jitter variance by 0.25, halving the shell thickness *σ*_*k*_ relative to training: sampling from these slightly tighter shells was empirically a little more stable. The direction ***d***^(*l*)^ is the analytic pseudo-C*β* unit vector: the position a C*β* would occupy given the residue’s N, C*α* and C atoms, placed at ideal tetrahedral geometry off the backbone frame and normalized to a unit vector from C*α* [24]. It is computed from backbone coordinates alone, so all angular spread in the source comes from the jitter term. The construction is applied at every residue position, including glycines, so that glycine state does not leak through the source distribution.

Because every slot begins in a distinct source neighborhood, its GHOST-versus-REAL state is informed by where it starts. Under an isotropic Gaussian source all slots instead share one initial distribution, and the GHOST/REAL decision must be resolved jointly with direction and radius only after coordinates have begun to organize. Across our training runs the nested shell prior appeared to make this optimization problem substantially more tractable, and we offer two hypotheses for why: (i) the nested radii already carry a coarse side-chain tree, in that slot *k* starts near the distance from C*α* at which the *k*-th heavy atom of a real side-chain tends to sit, so the source is closer to the data manifold than a single spherical Gaussian cloud; and (ii) because the shells are spread along a backbone-derived ray pointing the way the side chain projects — toward the receptor at buried and interface positions, into solvent at exposed ones — each subsequent slot is progressively exposed to whatever environment its own site actually presents, rather than all being drawn equally from one distribution centered on C*α*. Crucially, this differentiation is present from the very first reverse step: a slot does not have to wait for the cloud to organize to obtain unique context conditioning.

We present the nested shell prior as a workable and intriguing alternative to the ubiquitous isotropic Gaussian source. Ablations that quantitatively separate the prior’s contribution from other factors are ongoing. Note that the prior still leaks no per-position existence information at sampling: slot *k* of every residue is drawn from the same shell distribution regardless of that residue’s true labels, so the GHOST/REAL commitment remains a learned inference.

The same prior is used to pretrain on protein-bound small molecules. A ligand has no backbone frame, so the shared direction is taken toward the molecular centroid instead of along a pseudo-C*β* ray, anchored on a constructed pseudo-C*α* (Fig. S2); the two modalities’ measured shell statistics are given in Supplementary Table S1. Because the prior is built from geometry rather than residue identity in either case, the same formulation carries across protein-bound small molecules, single-chain monomers and peptide–protein complexes. This is what lets a single model be carried across all training phases, with one set of weights updated through small-molecule pretraining, peptide transfer, a combined stage adding non-canonical chemistry and antibody– protein interfaces, and a final peptide consolidation (§S10, §S14).

### Discretization at decode time

After sampling (Fig. 1C), each residue’s predicted atom cloud is assigned an identity by a *discretized* readout, applied at decode time only. To optimize the flow towards distinguishable atom configurations, a soft, purely geometric version of the same match is carried as an auxiliary loss in training. This helps push inter-atomic distance ratios toward configurations that map well to their GT identities, but is intentionally *not* a conditioning signal at sampling-time; leaving the atom flow maximum degrees of freedom. (§S7).

Our decode-time discretizer combines two components (Fig. 1D). The first is a purely geometric statistic: a normalized (atom-pairwise) distance matrix correlation between the predicted cloud and each reference rotamer, in the style of long-established techniques for comparison of structures via intramolecular distance matrices [32]. Additionally, this component carries static atom-count, element and chirality mismatch penalties — all critical, since a count- or handedness-blind match cannot disambiguate between very different atom-packing proposals.

The second is a *learned* component built on a similar featurization, with only per-slot element types and backbone *ϕ/ψ* angle geometry added. What distinguishes it is that it is a basic logistic regression fit on the model’s *own* sampled clouds, augmented with lightly corrupted reference rotamers as pseudo-data. Thus, it observes and can learn to adapt to the model’s actual (occasionally noisy; Fig. 2, Table 1) atoms output rather than idealized geometry; while still remaining restricted to the training dataset.

**Table 1:** Coordinates-first reconstruction quality on the 1k held-out test set (full-peptide de-novo design, 10 designs per peptide). Side-chain geometry, element identity and atom count are read directly off the generated clouds *before* any discretization. *Two-component mixture:* GHOST slots denote “no atom” and are designed to flow to the residue’s C*α*, while REAL slots settle at side-chain radii; the raw C*α* distances (no alignment) confirm the two components separate cleanly. Atom matching, the per-metric denominators and the *χ*1 restriction are given in §S15.

| Metric | Value |
| --- | --- |
| Side-chain RMSD (Kabsch, matched cloud) | 0.40 Å |
| Matched element accuracy | 0.75 |
| $\chi_1$ deviation (circular median $\vert \Delta\chi_1\vert $) | 23.9° |
| $\chi_1$ within 20° | 0.448 |
| Composition overlap (element multiset) | 0.67 |
| Per-residue atom-count MAE | 1.64 |
| Per-residue atom-count correlation ($r$) | 0.41 |
| GHOST slot $\rightarrow C\alpha$ distance | $0.27 \pm 0.14$ Å |
| REAL slot $\rightarrow C\alpha$ distance | $2.85 \pm 1.29$ Å |

**Figure 2:**
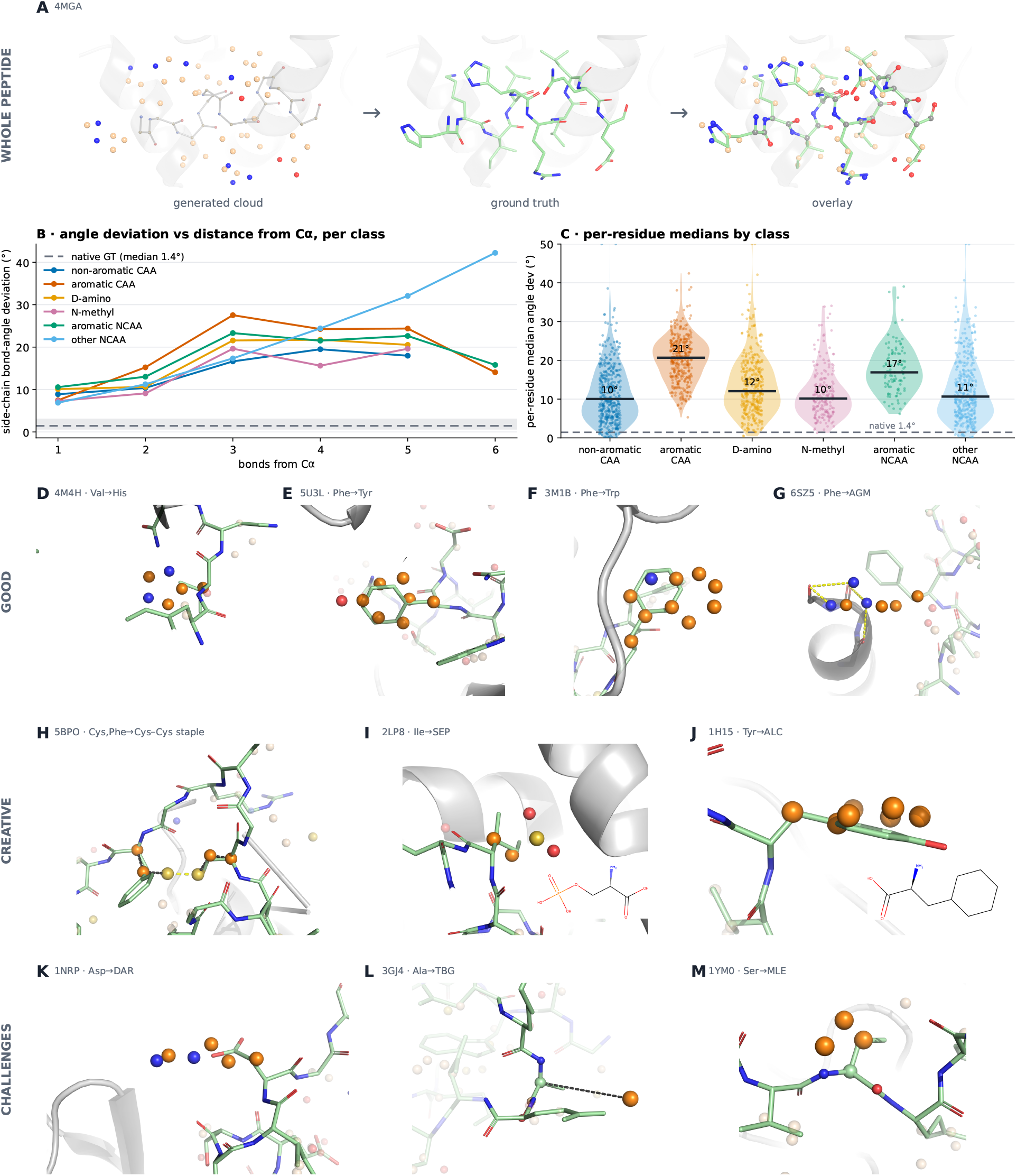
Generated atom clouds: accuracy, geometry, and chemical scope. **(A)** A whole-peptide generated cloud (4MGA), shown left to right as the generated atom cloud (its fixed backbone muted), the ground-truth peptide, and both overlaid. **(B, C)** Atom-cloud geometry evaluated against the ideal bond angles of the *assigned* residue identity: median side-chain bond-angle deviation versus bonds from C*α*, per residue class (B), and per-residue medians by class (C), from 10° (non-aromatic canonicals) to 21° (aromatic canonicals) against a 1.4° native-structure floor. **(D–F)** Examples of accurate atom placement for canonical side-chain ring systems. **(G)** A cloud finding polar contacts to the target backbone carbonyls via an arginine-variant arrangement (yellow dashes). **(H–J)** More exploratory cloud arrangements: a cysteine staple; an installed post-translational modification, phosphoserine (note the sulfur in the cloud, which the discretizer resolved to phosphorus); and a cloud assigned to a saturated cyclohexyl owing to its puckered placement (2D structures inset). **(K–M)** Challenging cases of poor or ambiguous atom placement: poor spatial organization (a collapsed guanidinium in an assigned D-arginine), a bulky *tert*-butyl reduced to a single misplaced atom, and an N-methyl-leucine assignment where the cloud may be converging on proline. Colors: generated side-chain cloud as element-colored spheres (highlighted-residue carbons orange (D–M), other carbons wheat; N blue, O red, S yellow), ground-truth peptide as green sticks, target as grey cartoon; labels read GT → generated (readout identity).

The two are mixed per candidate class: the learned component is the stronger ranker on canonical residues, where training clouds are plentiful, while the distance-matrix score does not fit a model and so treats a rare non-canonical the same as a common one; unbiased by data exposure. The blend therefore leans on the learned component for canonical classes and weights the two evenly for non-canonical ones, keeping each component functional in the regime it is good at. Full details of both components, and the blend itself, are in Supplementary Methods §S5 and §S5.3. Because the flow model itself does not depend on seeing residue-identity labels during training, a new NCAA is added by supplying one reference structure and simply refitting the decode-time logistic regression discretizer, a matter of minutes. No retraining of the generative model is required. Our current decode-time discretizer spans a 300-residue vocabulary — the 20 canonical amino acids plus 280 non-canonical types, detailed in full in Supplementary Table S9.

### Distinguishing design decisions

Three choices separate AtomWeaver from prior atom-level generative models [10, 12, 14, 19]: **(i)** no learned representation of residue identity gates coordinate formation, explicitly or implicitly; **(ii)** the per-slot ghost/real state is carried in the element vocabulary and modulates the coordinate flow at every reverse step, rather than being resolved by *post-hoc* packing downstream of a fully resolved cloud; **(iii)** the source distribution is a structured directed shell prior rather than an isotropic Gaussian. Decisions (i) and (iii) keep the residue-identity axis out of the learned continuous state, which is what enables non-canonical generalization at inference. AtomWeaver additionally carries auxiliary streams that encode a coarser zoom level, injecting both loose residue-level geometric features (approximations of side-chain centroid and first torsion *χ*_1_) and volumetric context into the atom-level flow at every equivariant layer. Further details are provided in Supplementary Methods, §S4.

### Model architecture

AtomWeaver utilizes a two-track denoiser on one shared timescale: an atom track carrying side-chain coordinates through ten equivariant blocks, and an element track running absorbing discrete diffusion. The two are coupled at a single FiLM junction, driven by the GHOST vs. REAL decision state, so that atom count and geometry are resolved together. Conditioning enters along two routes: residue-level encoders over the binder backbone and the target, whose outputs are injected into every equivariant block, and the target and backbone atoms themselves, which enter the equivariant graph directly as nodes. The full architecture, including the conditioning streams, the deep-injection bus, and the two feedback paths, is given in Supplementary Methods, §S1.

## 4. Results

### 4.1 Geometrical and structural analysis

AtomWeaver places and labels atoms in a step separate from whole-residue assignment, so the atom clouds can be evaluated in their own right; the final residue assignment, however, can influence how the quality scores are read. Figure 2(panel A) shows a full-peptide atom cloud for a test case with high ground-truth WT recovery (at the residue-assignment stage), displayed left to right as the generated cloud, the ground-truth peptide, and their overlay. The accuracy of cloud atom placement, measured as the three-atom angle deviation from the idealized geometry of the assigned residue, is smallest at the first bond from C*α* (≈7–11°, already several times the 1.4°native floor) and roughly doubles by the third bond before leveling off across most residue types (B). This accumulation of geometric error with distance from the fixed backbone is expected: positional noise compounds along the chain, and the training data are biased toward proximal atoms (every longer residue also carries the shorter ones). The lack of a plateau for the other NCAA class is, in part, a data-composition effect: at the sixth bond there are few examples and ∼90% of the measured angles come from a single residue, N^6^-acetyl-lysine, whose inherent flexibility makes terminal atoms especially hard to place. Note that to keep the cloud positions well-defined relative to the angular reference, analysis is restricted to clouds whose heavy-atom topology matches the assigned residue, which drops ∼24% of positions overall: 14% of canonical but 52% of non-canonical sites. Aromatics, both canonical and non-canonical, face the largest assembly challenge, having the highest per-residue angle deviations (panel C), again unsurprising, as forming an aromatic ring requires a coordinated flow of atoms into a single plane. Even so, clouds frequently take on identifiable ring geometry (panels D–F).

They also form novel contacts: for example a guanidinium reaching target backbone carbonyls (panel G), and more exploratory chemistries: a disulfide staple (panel H), a post-translational modification (phosphoserine, panel I), and a saturated-ring non-canonical resolved from a puckered cloud (panel J). Ambiguous or chemically unreasonable clouds also appear, marking areas for improvement: ill-formed functional groups (a collapsed D-arginine guanidinium, panel K), a fragmented residue (panel L; the “side-chain” is a single disconnected atom assigned as *tert*-butyl), and clouds ambiguous between an N-methyl residue and proline (panel M). Training the model to draw a sharper distinction between proline and N-methyl options is a clear avenue for gains.

Finally, we report several quantitative metrics in Table 1 to verify that the flow itself is behaving well before any residue identity is read. Side-chain atoms are placed quite accurately, within 0.40 Å RMSD of ground truth on average. Recovery of atom count, elemental makeup and precise rotameric state is more challenging, although assessment of this is difficult to disentangle from the generative model (rightly) considering alternate residue possibilities at each site. In any case, the atoms geometry is broadly consistent with the WT residue identity being scored highly by the model. From a mechanistic standpoint, the two-component mixture decomposes cleanly: measured as raw distances with no alignment, GHOST slots correctly collapse onto the residue’s C*α* while REAL slots settle out significantly further. This indicates healthy operational (FiLM) coupling of the element (existence) and coordinate flow tracks.

### 4.2 Joint design recovery

Identity is read off each resolved cloud by the 300-residue discretizer readout as described in §S5.3. The same discretizer is used for every result in this paper.

The one variation arises in the task-specific readouts used for the DMS benchmark (§4.5) and the Hirulog-3 analysis (§4.6): in each case, the readout is refit over a restricted residue set, using the same procedure with different output alphabets. Designing every site of a peptide jointly, AtomWeaver recovers the crystallographic canonical residue at 0.236 on the 1k held-out test set, using the same PDB-cluster-level split as PeptideMPNN [9] and restricting evaluation to the twenty canonical types (Table 2). The flow-side metrics behind these numbers, matched-cloud RMSD, atom-count MAE, and per-residue count correlation, are reported in Table 1.

**Table 2:** WT recovery (canonical top-1) on the 1k held-out test set, by structural environment. Every entry is a mean over 10 designs per site; *n* is the number of scored sites, a property of the ground-truth structures and therefore shared across methods. Buried, partial and exposed are relative-solvent-accessibility bins that partition the canonical positions: buried relSASA *<* 0.05, partial 0.05 ≤ relSASA *<* 0.25, and exposed relSASA ≥ 0.25. Interface is a *non-exclusive* cross-cut: a position whose side-chain loses at least 5 Å^2^ of solvent-accessible surface on binding is counted as interface *and* retains its free-chain burial class. Every row of this table, including the baselines, is binned by the same procedure, defined in full in §S19. The *T* column reports the baseline sampling temperature. Baselines emit residue labels directly and so pass through no readout of ours. AtomWeaver has no equivalent temperature parameter and therefore appears as a single row, sampled under its standard configuration. All methods design every site of the peptide, given only the fixed backbone and target with no native residue context retained, and all are scored over the same canonical twenty-residue candidate set. They differ in how those sites are resolved: the baselines decode autoregressively, committing one residue at a time over as many steps as the peptide is long, whereas AtomWeaver resolves every site in a single generative pass. The final block is not a design method but an empirical bound: how often the wild-type residue is among the best-measured substitutions at a position in deep mutational scanning, binned by the same environment criteria. It rests on 21,323 scanned positions across 382 domains for folding stability and 57 pooled protein–protein interface positions for binding. The protein–peptide scans in the same collection suggest that docked peptides may be somewhat less constrained still (0.125 over 24 interface positions), but that subset is small, so we conservatively report the higher protein–protein figure here. Note that this block uses the scans only to bound what wild-type recovery can be expected to reach; a direct comparison between each method’s residue rankings and measured ΔΔ*G* is reported separately in §4.5. Datasets, filters and per-dataset breakdowns for the bound are in §S19; §4.2 discusses what the comparison implies.

| Method | $T$ | Overall | Buried | Partial | Exposed | Interface |
| --- | --- | --- | --- | --- | --- | --- |
| *total sites* |  | 19,671 | 3,814 | 6,529 | 9,328 | 8,918 |
| FAMPNN [10] | 0.2 | 0.813 | 0.916 | 0.858 | 0.740 | 0.856 |
|  | 1.0 | 0.723 | 0.822 | 0.768 | 0.652 | 0.766 |
| LM-Design [33] | 0.2 | 0.586 | 0.766 | 0.730 | 0.411 | 0.709 |
|  | 1.0 | 0.495 | 0.691 | 0.638 | 0.315 | 0.611 |
| PeptideMPNN [9] | 0.2 | 0.583 | 0.775 | 0.719 | 0.409 | 0.721 |
|  | 1.0 | 0.491 | 0.678 | 0.619 | 0.326 | 0.619 |
| ProteinMPNN [7] | 0.2 | 0.479 | 0.729 | 0.681 | 0.235 | 0.661 |
|  | 1.0 | 0.348 | 0.581 | 0.500 | 0.147 | 0.491 |
| ESM-IF1 [34] | 0.2 | 0.449 | 0.744 | 0.660 | 0.181 | 0.629 |
|  | 1.0 | 0.373 | 0.664 | 0.546 | 0.133 | 0.528 |
| **AtomWeaver** | — | 0.236 | 0.379 | 0.308 | 0.128 | 0.305 |
| *Measured ceiling from deep mutational scanning (§S19)* |  |  |  |  |  |  |
| Folding stability | — | 0.273 | 0.524 | 0.351 | 0.158 | — |
| Interface binding | — | — | — | — | — | 0.421 |

Recovery splits sharply by structural environment (Table 2). Buried positions score highest (≈0.38); packing constrains them most tightly. Interface and partially-buried positions fall in between (≈0.30), and solvent-exposed positions score lowest (≈0.13). Both readouts produce the same ordering.

Exposed side-chains are genuinely under-determined: several rotamers and several residue identities fit the same local geometry. A model that recovered them as reliably as buried positions would therefore be reproducing one crystallographic choice rather than reading a constraint. The ordering across environments tells us more about the model’s behavior than any single pooled number does.

The discriminative baselines in Table 2 recover the deposited residue considerably more often than AtomWeaver does; they are not, however, performing the same task. Those methods emit a categorical distribution over the twenty canonical labels at each position, having been trained to place mass on the residue the PDB records in similar environments. At inference, they return that label directly: so recovering the deposited residue is both what they are optimized for and what they output. AtomWeaver never represents residue identity in its generative state, either as a label or as a latent variable that accumulates identity information over sampling. Thus, at no point during generation is there a quantity that could be steered toward a particular residue. What the model produces is a set of atom positions; an identity is attached afterwards, by matching that packed geometry against a reference library.

Table 2 reports the baselines at both their near-greedy deployment temperature (*T* = 0.2) and their nominal one (*T* = 1.0). The distinction matters for what is being compared: sharpening the logits concentrates mass on the single highest-probability label, which maximizes agreement with the deposited residue but narrows the set of residues the method will ever propose. At *T* = 1.0, with no such sharpening, the baselines sample across the breadth of their distribution. This is a little more similar in outcome to AtomWeaver, whose decode-time readout considers the whole 300-residue vocabulary. The closer correspondence shows up in the recovery pattern itself as well: relaxing the temperature moves every baseline toward AtomWeaver, and most so at exposed positions, where the environment constrains identity least: ProteinMPNN falls from 0.235 to 0.147 and ESM-IF1 from 0.181 to 0.133, against AtomWeaver’s 0.128. We therefore use *T* = 1.0 as the primary baseline setting throughout the paper.

Deep mutational scans (DMS) give an experimental handle on how much of that under-determination is real. By measuring every single-residue substitution at a position, they indicate directly how accommodating a backbone is of mutation. Empirically, these results show high tolerance of many alternate residue types; especially at less buried, less sterically constrained sites. The bottom block of Table 2 summarizes that as a rate: how often the wild-type residue is in fact among the best-measured substitutions at a position. Measured folding stability places it at 0.524 of buried positions but only 0.158 of exposed ones, and measured binding affinity at 0.421 of interface positions. Environments are binned by the same criteria used for our own rows, so the columns are comparable in definition; datasets, filters and per-dataset breakdowns are in §S19.

Deep mutational scan data suggests the distribution of proposed designs we would like a generative method to reach. Focusing on canonical residue breadth for now before we consider non-canonicals in the next section, we consider a site where many substitutions measure as well as the wild type. Here, a method that can only express the deposited residue is leaving most of the viable options unreachable. On the other hand, one that assigns them a meaningful share of its probability mass recovers greater valid design space. Whether AtomWeaver’s rankings track measured fitness, rather than merely spreading mass more widely, is tested directly in §4.5.

### 4.3 Non-canonical and stereochemical recovery

It is worth restating that in AtomWeaver, expanding the residue vocabulary requires no retraining of the generative model. A new type is added by supplying one reference structure and only refitting the decode-time discretizer, which takes minutes and does not touch the generative model’s weights. We report against a curated *300-residue vocabulary* (Supplementary Table S9): the 20 canonical residues plus 280 non-canonical types spanning post-translational modifications, halogenated and ring-substituted aromatic analogs, aliphatic and basic homologs, and modified cysteine, histidine and acidic residues. Types were retained from a larger design library when a reference rotamer set could be built for them reliably with RDKit [35]. The discretizer classifies on a 300-residue vocabulary with no catch-all class. Of the 280 non-canonicals, 274 are seen during training and 6 *are withheld* (MK8, 2MR, KCR, OMY, CGU, MP8): a genuine zero-shot set, carried in the discretizer by reference geometry alone (no sampled training clouds). All results below are from full *joint* peptide de novo design, with positions grouped by the ground-truth residue’s class, and all are scored under the single decode-time discretizer (§S5).

Two analyses follow, both reported in full in Supplementary §S15. First, the model resolves *stereochemistry* without being told: every site is initialized in the L-chiral direction — ie: the pseudo-C*β* ray of the source distribution is built L-handed — with no ground-truth chirality information supplied. From that purely L-initialized start the model recovers D at 69.6% of D-configured positions while mislabeling only 2.9% of L-chiral residues, separating the two at ROC-AUC 0.869 (Supplementary Fig. S3). Second, at positions whose native residue is non-canonical, the model surfaces seen non-canonical types near the top of the 300-residue vocabulary (median percentile 95.7, 47.0% in the top ten). Opening the vocabulary from 20 to 300 types costs essentially nothing on the canonicals. The six withheld types are the hardest category we report (median percentile ≈63), and should be read as an estimated floor on our per-type results rather than as an estimate of what zero-shot recovery can necessarily reach in principle. Per-type numbers for all 46 seen types and the six holdouts are in Supplementary Table S2.

### 4.4 Benchmarking self-consistency with de novo binder backbones

#### *In silico* refolding benchmark

To decouple WT recovery from design success metrics, we developed PeptideArena [36]^2^, a set of *de novo* designed peptide binders against 12 protein targets. The targets were selected to represent a diverse set of protein families (see Supplementary Table S3 for details). For each target, we generated 1000 peptide backbones of lengths *l* ∈ 8, 16, 24 using BoltzGen [21], for which hotspots were mined from existing structures and literature. The set of peptides backbones were then clustered and filtered, resulting in a total of 353 backbones (10 for each *l*-target pair, except for 2 targets). Using AtomWeaver and all baseline inverse folding models, we sampled 10 sequence designs per method for each of these backbones. Baseline models were ran with temperature *t* = 1.0. The designs were refolded with OpenDDE [37], an open-source all-atom biomolecular structure-prediction model, chosen here over comparable folding models for its improved handling of non-canonical residues. Each model was then evaluated by target-aligned peptide *self-consistency* RMSD (scRMSD). After refolding from the designed peptide sequence and the original target, the predicted and intended complexes are rigidly superposed using the target only; peptide backbone RMSD is then measured without a second alignment on the peptide. Thus scRMSD reports consistency of both the peptide backbone and its pose relative to the target, rather than peptide shape in isolation. This is the standard design– refold self-consistency principle [38, 39], adapted here for peptide–protein complexes. We intend to expand the target set in future work.

#### Refolding within margin

We use two scRMSD thresholds for considering a design success, <2 Å and <5 Å, where the former generally indicates that the sequence reproduces the exact binding pose while the latter roughly indicates a correct binding epitope with some conformational inconsistencies. Results of the bench-marking are in Fig 3A, and over the full target set in Supplementary Fig. S6. A random control, where sequences were unconditionally sampled based on the marginal distribution of the natural amino acids in our training set, is included to denote an empirical floor. Interestingly, the various sequence design methods show high concordance of success rates across the targets and peptide lengths. By these metrics, AtomWeaver is competitive with the field when designing peptides with canonical AAs. Additionally, scRMSD of the designs remain in the same regime even when the chemical design space is expanded to the much larger NCAA vocabulary. Notably, AtomWeaver-canon reported the highest <5 Å count on the 8-mers of 4Y7R and 6YOO. AtomWeaver-canon and AtomWeaver-open tied on the highest <5 Å count on 9CDT’s 16-mer designs. Note that we do not expect strictly better *in silico* scRMSD for NCAA-containing sequence designs against their canonical versions, as folding models trained on a canonical-dominated corpus typically have less confidence in NCAAs by nature.

**Figure 3:**
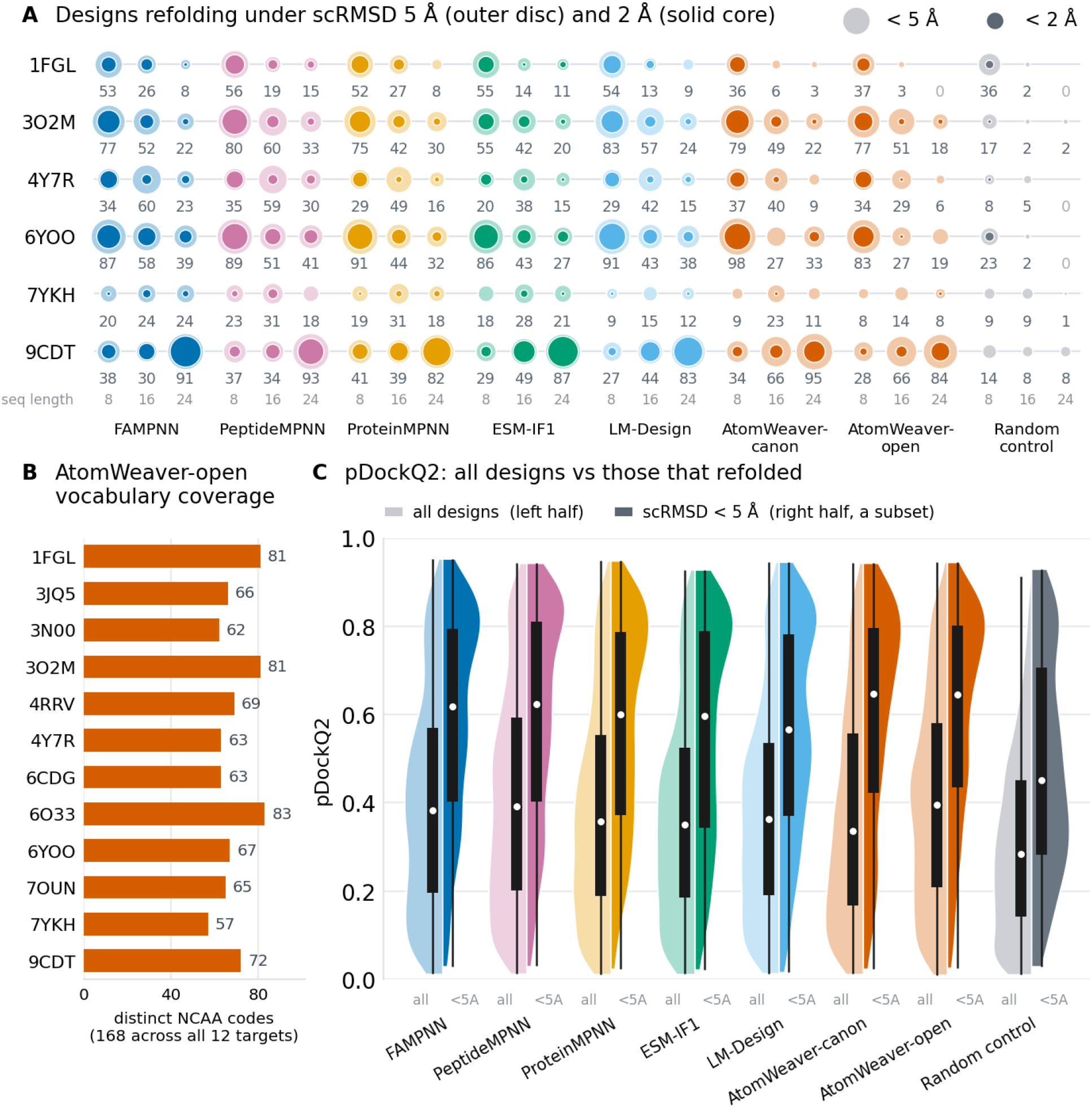
PeptideArena performance. (A) PeptideArena designs with target-aligned peptide scRMSD below 5 Å (outer disc) and 2 Å (solid core), by target and peptide length. The target protein is aligned before RMSD is computed over the peptide backbone, so the metric assesses the designed binding pose as well as peptide conformation. Each cell holds one disc per peptide length, 8 / 16 / 24 from left to right. The annotation corresponds to the number of designs passing the 5 Å threshold (outer-disc size). Random control is a randomly sampled sequence based on natural amino acid frequency in the training set. Only the six targets with highest average success rates across models are shown for visibility; the full plot is in Supplementary Figure S6. (B) Number of unique NCAAs emitted by AtomWeaver-open across all targets. AtomWeaver sampled 168 unique NCAAs across 3.5k sampling draws. (C) Distributions of pDockQ2. For each arm, we show the distribution over all designs (left half) beside the subset with target-aligned peptide scRMSD below 5 Å (right half).

#### Non-canonical usage

Across the 3.5k designs, AtomWeaver-open emitted 168 unique NCAAs, covering 60% of the NCAAs in our allowed set (Figure 3B). Four of the designed NCAAs (2MR, CGU, MP8, and KCR) were never seen by the model during training. This showcases the ability of our model to generate a large diverse variety of side-chain modifications. Although the amount of NCAAs per sequence was not capped (i.e., the model could design a full-NCAA sequence), we observed that it emitted a median of two NCAAs per sequence and a 90th percentile of four NCAAs in a single sequence; implying that AtomWeaver-open still has general preference for canonical residues.

#### Interface confidence

We also report on the pDockQ2 [40] of the designs based on OpenDDE’s refolding. pDockQ2 estimates the model’s confidence that a predicted interface is real. Figure 3C shows the pDockQ2 distribution over all designs beside the subset whose peptide refolded to within 5 Å scRMSD. Among designs that refolded within <5 A, both AtomWeaver variants lead (0.647 canon, 0.646 open) against 0.566–0.623 for the baselines. Unfiltered, AtomWeaver-open has the highest median of the eight methods (0.396, against 0.350–0.391 for the baselines), despite having the fewest total refolded designs, indicating that strong interface confidence can be maintained even with heavy inclusion of non-canonical content.

### 4.5 Deep mutational scan benchmark of two peptide binders

#### Data

WT recovery tests whether a model reproduces the deposited residue, but not whether it ranks experimentally measured alternatives by function. We therefore benchmark AtomWeaver against binding free-energy changes from the nonproteinogenic deep mutational scan of Rogers et al. [41]. The scan measures

ΔΔ*G* for canonical and 21 non-canonical substitutions at each assayed position in two peptide–target systems, introduced by flexizyme-charged tRNA in an mRNA-display selection. Both have solved complexes: PUMA BH3 bound to MCL1 (PDB 2ROC, chains B and A), and the macrocyclic peptide CP2 bound to KDM4A (PDB 5LY1, chains E and C). PUMA was scanned as a 34-mer, but positions 4–30 are the 27 positions resolved in the complex and used for inverse folding. CP2 contributes 12 positions (residues 2–13). Overall, the benchmark contains 39 positions and 1,548 measured mutants: 729 canonical and 819 non-canonical substitutions.

#### Benchmarking setup

We report two candidate sets at each position. *CAA-only* ranks the canonical substitutions measured in the scan (19 for PUMA and 18 for CP2; the published CP2 scan omits methionine, and both sets exclude the wild-type residue). *CAA+NCAA*, unsupported by canonical inverse-folding baselines, ranks canonical and non-canonical substitutions together. This mixed candidate set tests whether an open-vocabulary method can select from the complete assayed chemical space, rather than forcing a non-canonical choice by excluding canonical alternatives.

Orthogonal to the candidate alphabet is the design regime. In *single-site* design, each query position is inpainted independently while every other position is held at its deposited identity, mirroring the scan’s one-substitution, otherwise wild-type background. In *joint* design, all peptide positions are sampled together in each atom-cloud draw. The site-specific ranking is read from the marginal distribution across these coupled draws, integrating over identities selected at the other positions. This setting also amortizes sampling across the peptide, avoiding a separate inpainting trajectory for each of the 39 positions.

We compare AtomWeaver with ProteinMPNN [7], PeptideMPNN [9], ESM-IF1 (causal) [34], LM-Design [33], FAMPNN [10], and NCFlow. The canonical-label baselines are evaluated only in CAA-only, whereas AtomWeaver and NCFlow also support the mixed candidate set. For joint design, AtomWeaver samples all peptide positions in one coupled atom-cloud trajectory; the baselines use their model-native multi-site procedures, which differ from AtomWeaver’s simultaneous joint conditioning (Supplementary Section S17); complete baseline protocols are given in the Supplementary Methods. UNAAGI [18] is the closest generative inverse-folding model with non-canonical support, but was not publicly available for this benchmark. NCFlow is retained as a reference-only comparator: it is not an inverse-folding model, but scores a user-defined residue set at a fixed backbone position through AEV-PLIG [17].

#### Metric

We report mean within-site agreement: the sign-reversed Spearman correlation, *™ρ*, between each candidate’s model score and measured DMS ΔΔ*G*, such that larger positive values indicate stronger agreement. Values are averaged separately within CP2 and PUMA, and their equal-system average is reported as the mean. AtomWeaver rows use 100 stored draws. Joint-design baseline rows use 100 draws at sampling temperature *T* =1.0, and the explicitly labelled sensitivity rows use *T* =0.2. Temperature controls only these stochastic joint-design samplers. Single-site baselines instead read their native conditional distributions at each queried position, with all other peptide positions fixed; they do not draw a sequence and therefore have no sampling-temperature parameter. A separate logit-temperature scaling could be applied as probability calibration, but it is not part of this model-native single-site protocol; the rank-based statistic is invariant to positive scaling. Candidate curation and score aggregation are specified in Supplementary Section S17.

#### Generator-fixed vocabulary extension

For this benchmark, we evaluate three learned readouts:

AtomWeaver-canon over the 20 canonical amino acids, AtomWeaver-DMS over the 41 identities assayed in the scan, and AtomWeaver-open over the full 300-residue library. The AtomWeaver generative model and its sampled atom clouds are held fixed; only the post-hoc identity readout changes. We therefore call this a *generator-fixed vocabulary extension*; it does not imply that the discriminative readout itself is zero-shot.

For benchmark construction and evaluation details, see Supplementary Section S17.

#### Findings

Table 3 reports single-site and joint-design agreement separately. AtomWeaver labels identify the decode-time reference library: *open* uses the full 300-residue library, *DMS* the mixed 41-residue DMS library, and *canon* the 20 canonical amino acids. In CAA-only single-site design, AtomWeaver-open has the numerically highest observed inverse-folding equal-system mean (0.274). The system-level results remain complementary: FAMPNN has the numerically highest observed CP2 value (0.354), whereas AtomWeaver-DMS has the numerically highest observed PUMA value (0.269).

**Table 3:** DMS Spearman rank agreement for single-site and joint design. Entries are mean within-site, sign-reversed Spearman rank correlation (*™ρ*), reported separately for CP2 and PUMA. Mean is the equal-system average of the CP2 and PUMA values. The Library column gives the canonical and non-canonical identities available to the readout, rather than identities selected in a design. All AtomWeaver and joint-design baseline rows use 100 stored draws. The double dagger identifies the *T* =0.2 joint-design baseline sensitivity setting; all other baseline rows use *T* =1.0 for joint design. Single-site baseline agreement instead uses temperature-invariant model-native conditional scores. Bold indicates the numerically highest inverse-folding result in a column, not a statistically resolved pairwise difference; stratified site-bootstrap confidence intervals are reported in Supplementary Table S5. NCFlow is a reference-only comparator and is not eligible for bolding.

| Method | Library | Single-site design |  |  |  |  |  | Joint design |  |  |  |  |  |
| --- | --- | --- | --- | --- | --- | --- | --- | --- | --- | --- | --- | --- | --- |
|  |  | CAA-only |  |  | CAA+NCAA |  |  | CAA-only |  |  | CAA+NCAA |  |  |
|  |  | CP2 | PUMA | Mean | CP2 | PUMA | Mean | CP2 | PUMA | Mean | CP2 | PUMA | Mean |
| AtomWeaver-open | 20 + 280 | 0.315 | 0.233 | **0.274** | 0.208 | 0.126 | 0.167 | 0.283 | 0.244 | **0.264** | 0.195 | 0.136 | 0.165 |
| AtomWeaver-DMS | 20 + 21 | 0.220 | **0.269** | 0.244 | **0.222** | **0.151** | **0.186** | 0.210 | **0.293** | 0.251 | **0.202** | **0.154** | **0.178** |
| AtomWeaver-canon | 20 + 0 | 0.199 | 0.259 | 0.229 | – | – | – | 0.202 | 0.279 | 0.241 | – | – | – |
| ProteinMPNN ($T=0.2$)<sup>‡</sup> | 20 + 0 | – | – | – | – | – | – | 0.280 | 0.187 | 0.233 | – | – | – |
| ProteinMPNN ($T=1.0$) | 20 + 0 | 0.294 | 0.207 | 0.250 | – | – | – | 0.296 | 0.172 | 0.234 | – | – | – |
| PeptideMPNN ($T=0.2$)<sup>‡</sup> | 20 + 0 | – | – | – | – | – | – | 0.372 | 0.117 | 0.244 | – | – | – |
| PeptideMPNN ($T=1.0$) | 20 + 0 | **0.357** | 0.139 | 0.248 | – | – | – | **0.372** | 0.125 | 0.249 | – | – | – |
| ESM-IF1 ($T=0.2$)<sup>‡</sup> | 20 + 0 | – | – | – | – | – | – | 0.232 | 0.191 | 0.212 | – | – | – |
| ESM-IF1 ($T=1.0$) | 20 + 0 | 0.236 | 0.251 | 0.244 | – | – | – | 0.223 | 0.217 | 0.220 | – | – | – |
| FAMPNN ($T=0.2$)<sup>‡</sup> | 20 + 0 | – | – | – | – | – | – | 0.251 | 0.185 | 0.218 | – | – | – |
| FAMPNN ($T=1.0$) | 20 + 0 | 0.354 | 0.132 | 0.243 | – | – | – | 0.251 | 0.185 | 0.218 | – | – | – |
| LM-Design ($T=0.2$)<sup>‡</sup> | 20 + 0 | – | – | – | – | – | – | 0.161 | 0.249 | 0.205 | – | – | – |
| LM-Design ($T=1.0$) | 20 + 0 | 0.114 | 0.245 | 0.179 | – | – | – | 0.155 | 0.262 | 0.208 | – | – | – |
| NCFlow side-chain [16]<sup>†</sup> | – | -0.008 | 0.151 | 0.071 | 0.145 | 0.363 | 0.254 | -0.008 | 0.151 | 0.071 | 0.145 | 0.363 | 0.254 |
| NCFlow peptide<sup>†</sup> | – | 0.289 | 0.161 | 0.225 | 0.392 | 0.322 | 0.357 | 0.289 | 0.161 | 0.225 | 0.392 | 0.322 | 0.357 |

Library: candidate identities available to the readout.
<sup>‡</sup> $T=0.2$ : joint design only.
<sup>†</sup>Reference-only affinity scorer.

In CAA-only joint design, AtomWeaver-open has the numerically highest observed inverse-folding equal-system mean (0.264), ahead of AtomWeaver-DMS (0.251), PeptideMPNN at *T* =1.0 (0.249), and PeptideMPNN at *T* =0.2 (0.244). PeptideMPNN at *T* =1.0 has the numerically highest observed CP2 value (0.372), whereas AtomWeaver-DMS has the numerically highest observed PUMA value (0.293), illustrating system-specific behavior under the equal-system aggregate. The temperature panel changes the stochastic joint-sequence ensemble. Single-site rows instead read conditional distributions without sampling a sequence; no separately tuned logit-temperature calibration is reported. AtomWeaver jointly samples all peptide positions in one coupled atom-cloud trajectory; its result therefore shows that the coordinate-to-identity readout retains functional ranking signal when the rest of the peptide is redesigned concurrently.

In the CAA+NCAA benchmark, canonical and non-canonical candidates compete in the same set. AtomWeaver-DMS has the numerically highest observed inverse-folding value in every reported column, with equal-system agreement of 0.186 in single-site design and 0.178 in joint design. It exceeds AtomWeaver-open in this mixed setting, showing that a readout tailored to the relevant candidate library can extract more functional ranking signal while the atom-cloud generator remains fixed.

The stratified site-bootstrap intervals in Supplementary Table S5 place these point estimates in context, quantifying variation across benchmark positions conditional on the fixed DMS measurements and stored model draws (not independent experimental-replication interval). They provide uncertainty for each benchmark statistic, not a pairwise significance test, so small differences between the reported means should not be read as statistically resolved effects.

NCFlow is displayed only as a reference-only affinity-scoring comparator, because it evaluates specified ligand–receptor arrangements rather than inverse-folding residue rankings; its values are not part of the inverse-folding comparison.

### 4.6 Case study: Hirulog-3 bound to thrombin

This case study asks how AtomWeaver can support medicinal chemistry on an existing peptide lead, where objectives extend beyond affinity optimization to exploring broader chemical space for modifications compatible with target engagement and developability. We therefore focus on the recovery and exploration of non-canonical and stereochemical choices compatible with the bound complex.

#### The Hirulog-3 peptide

Hirulog-3 is a 20-residue, bivalent thrombin inhibitor derived from the much larger natural hirudin scaffold (complexed with human *α*-thrombin in PDB 1ABI [42]). Its non-canonical chemistry is a medicinal-chemistry design choice, not simply an attempt to maximize affinity: the sequence contains a D-phenylalanine (DPN, residue 1) and a *β*-homoarginine (HMR, residue 3). Although 1ABI labels residue 12 as DPN, we treat it as L-Phe: the reported Hirulog-1/-3 C-terminal sequence is DFEEIPEEYL, placing its canonical Phe at residue 12 [43]. HMR’s additional methylene group displaces the scissile bond from thrombin’s catalytic Ser195 while preserving the key guanidinium interaction, making the derivative non-cleavable. We therefore use Hirulog-3 to ask whether AtomWeaver can recover, from a fixed experimental complex, the stereo-chemical and backbone-adjacent chemistry of a compact thrombin-binding scaffold designed for proteolytic resistance. Hirulog-3 was excluded from AtomWeaver training, whereas thrombin was present in the training set. This is therefore a peptide- and chemistry-held-out test on a fixed experimental complex, rather than a receptor-held-out generalization test.

#### Recovery with a D/*β*-aware vocabulary

We use the experimentally determined Hirulog-3–*α*-thrombin complex with both AtomWeaver-open, which uses the full 300-residue readout, and AtomWeaver-DBeta, which applies a refit 43-residue discretizer to the same generated atom clouds. The DBeta readout contains the 20 canonical amino acids, the 19 D-amino-acid counterparts of the chiral canonical amino acids, and four *β*-amino acids drawn from the 300-residue library (B3L, B3Q, HMR, and HT7). No separate generative model is trained for this setting. Residue probabilities are averaged over 100 generated atom clouds before ranking.

HMR is a *β*-homoarginine and, like other *β*-amino acids, contains an additional backbone methylene relative to its *α*-amino-acid counterpart. Including all four *β*-amino acids, rather than only HMR, asks whether the full AtomWeaver pipeline, from atom-cloud generation through identity readout, can recognize this altered chemistry and distinguish it from closely related alternatives. This is deliberately a demanding boundary case: AtomWeaver performs fixed-backbone side-chain design and was not developed to generate or remodel backbone-modified peptides. The experiment therefore tests recognition of D-residue and *β*-amino-acid choices compatible with a provided, chemically modified backbone, rather than backbone modification itself.

The D/*β*-aware AtomWeaver-DBeta readout improves recovery of the deposited Hirulog-3 chemistry when stereochemical and backbone-modified alternatives are available in the same library. The same atom-cloud generator also retains recovery under AtomWeaver-open, where each deposited identity is ranked against the full 300-residue library. Recall-at-*K* curves for both readouts are reported in Supplementary Fig. S5 (Supplementary Section S18), illustrating useful chemical discrimination across both a focused medicinal-chemistry library and a more open design space.

#### Prospective design prioritization

The same fixed complex can be used to generate model-prioritized, structurally plausible testable Hirulog-3 analogues. We refold AtomWeaver’s top single-site suggestions with OpenDDE and use target-aligned peptide scRMSD together with pDockQ2 to prioritize candidates for follow-up. Per-site sequence-logo diagnostics for the single-site and joint settings are reported in Supplementary Figs. S8 and S9 (Supplementary Section S18).

Supplementary Fig. S7 provides a conservative prospective-design screen rather than an affinity claim. Both reference libraries prioritize substitutions at Gly5; the AtomWeaver-open Gly5 → Asp analogue has higher predicted complex confidence in all ten paired OpenDDE refolds, increasing mean pDockQ2 from 0.856 for the corrected native to 0.888 while maintaining a low mean target-aligned peptide-backbone scRMSD of 1.53 Å. The structural subpanel provides an interaction-level hypothesis for this candidate. pDockQ2 is a predicted complex-confidence score, and neither it nor scRMSD measures binding potency or proteolytic stability.

#### Explorative joint design

Designing every site at once, rather than one substitution at a time, is where AtomWeaver-open’s broad vocabulary becomes visibly exploratory. Figure 4 shows a joint atom-cloud design decoded with AtomWeaver-open’s full 300-residue discretizer, refolded with OpenDDE in the thrombin complex, and superposed on the deposited structure. Thus, the coordinated non-canonical proposal is read directly from the open-library setting, rather than from a Hirulog-specific candidate set. Despite multiple non-canonical substitutions, the refolded peptide retains a close backbone and interface pose, making the proposed interactions interpretable in the experimental context.

**Figure 4:**
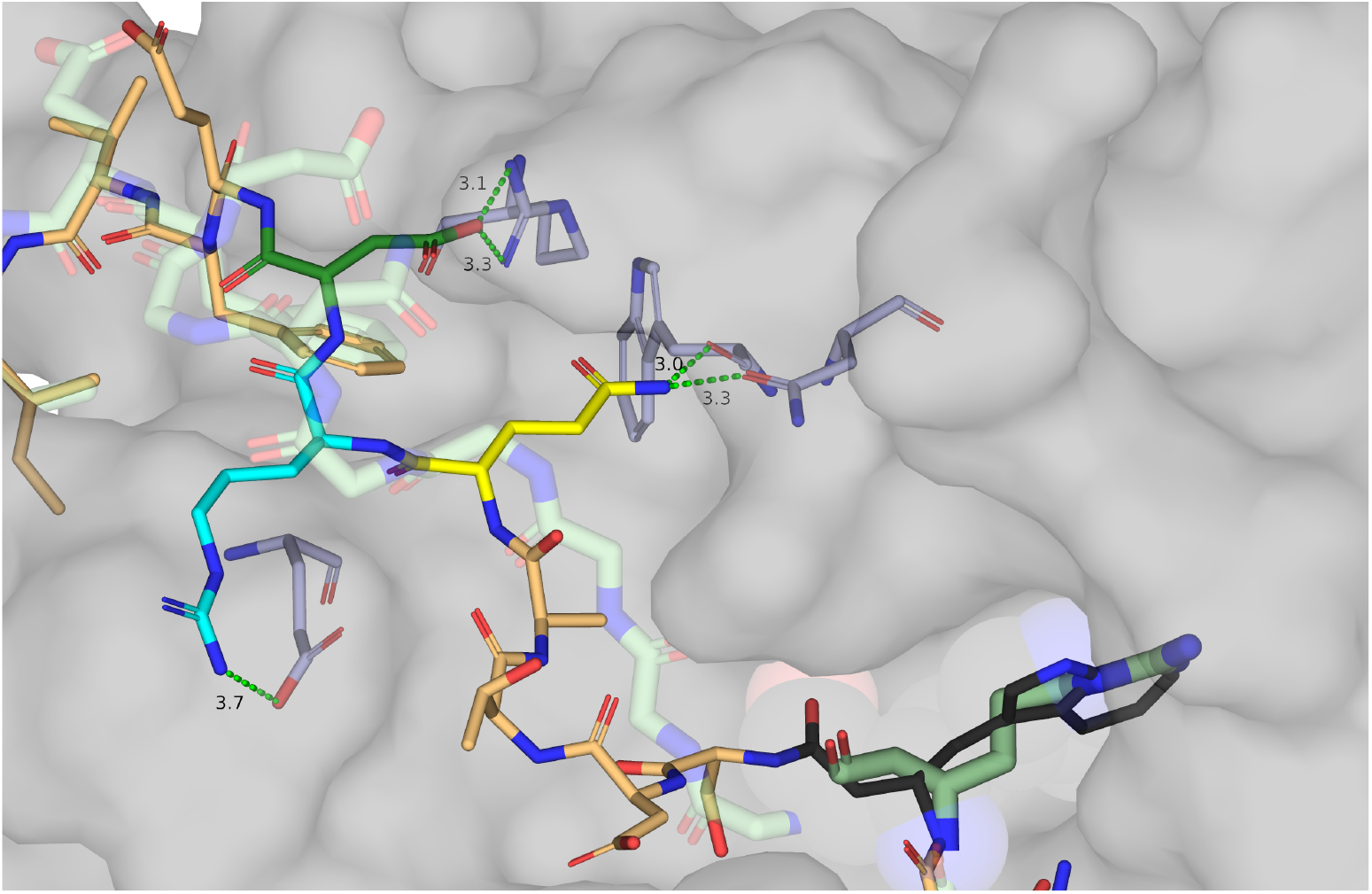
OpenDDE-refolded AtomWeaver-open Hirulog-3 joint design. The Hirulog-3 *β*-homoarginine is substituted for a *β*-homotryptophan (black), exchanging the salt bridge to Asp189 for pocket-filling contacts. An Asn at position 8 (yellow) is poised for hydrogen bonds to Asn143 and the Trp141 backbone. An Arg9 (cyan) reaches for a new salt bridge to Glu25, and an Asp10 (dark green) forms a salt bridge with Arg68, slightly register-shifted from the native contact. The Asn–Arg–Asp motif appears alongside a D-alanine at position 7. The AtomWeaver-open design was refolded with OpenDDE in complex with thrombin before visualization; despite these non-canonical substitutions, the predicted peptide retains a close backbone and interface pose relative to deposited Hirulog-3. Ground-truth Hirulog-3 is shown in green, the refolded design in light orange (unless highlighted), and the thrombin surface in grey; dashed green lines mark polar contacts with distances in Å.

The *β*-homoarginine of Hirulog-3 is exchanged for a *β*-homotryptophan, trading the salt bridge to Asp189 for pocket-filling contacts. Three further positions reorganize together: an Asn at position 8 poised for hydrogen bonds to Asn143 and the Trp141 backbone, an Arg9 reaching a new salt bridge to Glu25, and an Asp10 forming a salt bridge with Arg68 that is register-shifted from the native contact. That the Asn–Arg–Asp motif appears alongside a D-alanine introduced at position 7 is the kind of coupled, stereochemistry-aware proposal a site-by-site search cannot readily reach. We report these as structural hypotheses, not experimentally validated analogues.

## 5. Discussion

AtomWeaver is a multi-component flow-matching framework for peptide inverse folding in which side-chain atoms are placed in coordinate space *first*, conditioned on the binder backbone and target protein, and only afterwards read off to residue identity by a decode-time discretizer. The open reference vocabulary can be freely adjusted by the user at decode time: removing existing residues, adding new residues, and/or modifying each residue’s rotamer structures. Note that the discretizer has been kept intentionally simple: a basic logistic regression for the training-cloud-informed learned component, combined on equal terms with a geometric distance-matrix statistic in the noncanonical domain to avoid overfit.

Because residue identity never enters AtomWeaver’s learned continuous state, the addressable design space at inference is prescribed by the contents of that vocabulary rather than by a fixed twenty-letter alphabet or a learned representation of residue identity. In other words, the model proposes non-canonical residues whenever its predicted atom cloud matches one better than any canonical residue, *without* needing to have seen that residue in training and *without* any user prompt for the intended chemistry. The technical ingredient we consider most responsible is the newly introduced *nested shell* prior: per-slot overlapping radial shells along a shared, backbone-derived direction, which give each atom slot- and target-specific geometric context from the first reverse-sampling step and, we hypothesize, let the model commit GHOST/REAL atom existence earlier than it could from an unstructured cloud. The new prior worked well for us and is worth wider attention; a robust comparison that would establish how much of the gain it contributes, rather than other design choices made alongside, is in progress (§S3).

### Functional ranking under canonical design

The DMS benchmark complements WT recovery and structural self-consistency with a functional test: within a fixed position, does a model rank substitutions in agreement with the measured binding-energy preferences? AtomWeaver-open has the numerically highest observed CAA-only equal-system agreement in both single-site (0.274) and joint design (0.264). The primary *T* =1.0 joint panel is complemented by a matched *T* =0.2 sensitivity panel. At *T* =1.0, PeptideMPNN reaches 0.249 overall (0.244 at *T* =0.2) and has the highest observed CP2 value (0.372), whereas AtomWeaver-DMS has the numerically highest observed PUMA value (0.293).

For joint design, each comparator is evaluated with its own standard sequence-generation procedure rather than being forced into a common sampler: AtomWeaver resolves all peptide positions simultaneously, while the other methods construct a multi-position sequence through ordered autoregressive, progressive-unmasking, or masked-segment-inpainting steps (§4.5). The resulting conditional distributions are therefore not identical across methods. Even under this deliberately heterogeneous comparison, AtomWeaver’s joint result shows that its coordinate-to-identity readout retains site-specific functional ranking signal while the rest of the peptide is redesigned.

### An experimentally testable open chemical space

The CAA+NCAA comparison exposes a distinction that is hidden by canonical-only benchmarks: most established inverse-folding methods cannot assign scores to the non-canonical candidates, whereas AtomWeaver can rank them alongside canonical residues from the same generated atom clouds. AtomWeaver-DMS has the numerically highest observed inverse-folding value in every CAA+NCAA column. Together with the site-bootstrap intervals in Supplementary Table S5, these point estimates support a library-dependent functional-ranking signal but not a statistically resolved pairwise winner claim. This result concerns a fitted DMS-41 readout, not zero-shot classification by an arbitrary newly added template. This is a practical advantage for peptide design, where useful chemistry is often target- and campaign-specific: new residue templates can be added to the reference library and evaluated in the same structural design workflow, rather than being restricted to a fixed twenty-label output layer.

### A broader, physically viable design space

The results demonstrate that the set of sequences a fixed backbone:target complex can physically accept is larger than the recovery-to-wildtype criterion suggests. AtomWeaver’s atoms-first generator is able to populate this set. Joint all-site design attains the canonical recovery reported in §4.2, but the sharper evidence is independent of any single reference. The deep mutational scan (§4.5) shows that these backbones tolerate substantially more non-wildtype substitution than recovery alone would suggest, so the breadth we exploit is a property of the biology rather than an artifact of a permissive decoder. The discriminating test is structural: designs are assessed by whether they refold into the intended backbone and binding pose rather than by how closely they reproduce a deposited sequence (§4.4).

It is worth stating the consequence for recovery directly, since the headline numbers invite the opposite reading. Measured against the deep mutational scans rather than against the crystal structure, our atom-first approach, with no residue identity fixed in advance or accumulating in a learned representation, recovers considerably more of the DMS-accessible residue space than its wild-type recovery suggests and tracks the measured ceiling more closely than the discriminative baselines do (Table 2). The baselines’ advantage on wild-type recovery is largest exactly where the scans say the constraint is weakest: at exposed, sterically unconstrained positions, where they return the deposited residue several times more often than physical measurement can account for. That surplus is evolutionary precedent asserting itself, and it is genuinely valuable when a well-precedented interface is the goal. It does not, however, indicate that those positions are more constrained than the experimental measurements show. For therapeutic peptide design the distinction is consequential: novelty relative to the deposited sequence is frequently the objective rather than a failure mode, whether to improve stability or selectivity, to reach chemistry that natural sequences do not contain, or to escape existing intellectual property. At weakly constrained positions a departure from the crystal residue is therefore better read as the design space the scans report than as an error.

### Open vocabulary and non-canonical design

Because identity is read at decode time, the same mechanism that packs canonical side-chains also proposes non-canonical ones, including residues withheld from training entirely (§4.3 and §4.4) and, in the Hirulog-3 case study, the non-canonical residues of a clinically relevant therapeutic peptide (§4.6). Identity-first inverse-folding methods cannot do this: their output space is a fixed set of categorical labels, whereas here extending the addressable chemistry costs one reference structure and no retraining of the generative model.

### A geometric likelihood that any prior can be read against

Because the readout produces a distribution over the whole reference vocabulary from geometry alone, rather than a single label, its output is usable as *evidence* rather than as a verdict. Any prior over residue identity can be multiplied against it and the two weighed explicitly. Figure S4 shows the simplest case: composing the readout with a natural-frequency prior over the canonical classes shifts the designed composition toward the common residues and pulls the non-canonical share below its native rate. Nothing about that composition is special to natural frequencies. The same slot can accept a discriminative inverse-folding prior, such as an MPNN-family posterior over the canonical twenty, so that a site is decided by geometric fit and sequence precedent together, with the relative weight a knob rather than an architectural commitment. This is the practical form of the ordering argument of §3: an identity prior can be layered onto a coordinates-first model, while an open vocabulary cannot be recovered from a model that only ever emits twenty labels. We do not pursue these compositions here, but they are available without retraining, and they are the natural way to use this model where evolutionary precedent is desired alongside physical target pocket fit.

### Stereochemistry resolved from geometry

Because the pseudo-C*β* ray places each residue on one side of the backbone plane at *t* = *T* , handedness could in principle be inherited from a source distribution hint rather than inferred. *It is not*: from a uniformly L-chiral start the model still crosses the mirror plane at D sites, at a specificity that rules out indiscriminate D-calling (§4.3). Bulkier side-chains resolve their handedness earlier in sampling than smaller ones (Fig S3c), consistent with steric determinacy: a larger side-chain admits fewer viable tetrahedral placements, so its chirality is more constrained by the backbone and pocket, and the learned prior reflects this as higher, earlier confidence. This matters beyond the D-amino acids themselves. Identity is assigned by comparing inter-atomic distances, and a distance matrix is invariant under reflection — a side-chain and its mirror image have identical distance matrices — so handedness cannot come from that comparison at all. It is supplied instead by the reflection-odd backbone *ϕ/ψ* features of the learned component and by an explicit chirality gate (§S5). Without that machinery an open-vocabulary readout could not separate any enantiomeric pair.

### Limitations and extensions

Peptide length is capped at 32 residues in the current training corpus. The X element in the element vocabulary was used as a catch-all due to sparse data coverage preventing confident separation of sulfur, phosphorus, and halogen options. In future versions of the model we intend to add discrimination between such element types, as they can result in quite different noncanonical residue suggestions that currently cannot be reliably disambiguated.

The atom clouds themselves carry a geometric cost that the identity readout partly absorbs: bond-angle deviation is already several times the native floor at the first bond from C*α* and roughly doubles by the third (Fig. 2, panels B and C), so error accumulates with distance from the fixed backbone. Controlling that accumulation — through a loss that penalizes angular error directly rather than only through the distance-matrix term, and through training data less weighted toward proximal atoms — is the change we expect to matter most in the next version of the model. Aromatic rings are the specific case to beat, canonical and non-canonical alike, since assembling a planar ring demands a coordinated flow of atoms into a single plane. The geometric flow metrics (Table 1) point the same way: further improvements to side-chain atom placement, rotameric precision, and atom count precision are expected in future iterations of the model and should translate to even stronger inverse folding suggestions.

### Positioning

Identity-first, latent-mediated and atom-first generation answer different questions about the same site: what selection installed and a fixed vocabulary can name; what a learned embedding reaches past that vocabulary, given a prompt or matched training; and what the site will physically accommodate, from which a residue identity is read at decode time. The mutational scan indicates that the space this opens is populated by substitutions the biology tolerates. A prompting head could be added *on top* of a coordinates-first framework, whereas recovering an open-ended vocabulary from a model whose output space is a fixed set of categorical labels is not available by definition. In practice, for AtomWeaver extending the design space costs one reference structure and no generative model retraining. Where a high-likelihood canonical sequence in a well-precedented interface is wanted, the evolutionary prior is an asset the identity-first family can exploit directly.

AtomWeaver’s access to previously unreachable design space most suits contexts that can tolerate the higher risk of NCAA integration, in synthesis cost and less-characterized building blocks, in exchange for properties such as cell permeability and proteolytic stability a twenty-letter vocabulary cannot reach. Our working hypothesis for the mechanism is that an atoms-first treatment never collapses the site to a narrow twenty-residue choice, and so retains degrees of freedom that a model committing to an identity first has already spent. This is compounded by joint all-site prediction and by a source distribution that hands every atom interface-contextual information before the first reverse step. That the model also *chooses* well within the open vocabulary is empirical, established here on multiple systems; separating the hypothesized mechanism from the alternatives (the training corpus, the discretizer’s tolerance, the reference library’s composition) would require targeted ablations we have not run.

## 6. Conclusion

AtomWeaver reframes docked peptide–protein inverse folding as a coordinate-first problem. It generates side-chain atom clouds containing coordinates, elements, and occupancy, then assigns identity by geometric agreement with a chosen reference library. This separation of generation from chemical decoding makes the design vocabulary an inference-time decision: the same generator can be read out over canonical amino acids, D-amino acids, and exotic non-canonical chemistries without retraining the generative model for every library. Complementing the architecture’s explorative tendency, its staged training combines small-molecule pretraining with synthetic NCAA-augmented protein monomers and dimers, antibody–protein interfaces, and peptide–protein complexes, further broadening the range of side-chain atom conformations it will consider for a target pocket.

On the challenging new PeptideArena benchmark that we introduce alongside this work, we show that AtomWeaver’s non-canonical integration into its designs is not merely diversity-expansion: topical structural models refold these designs back to the backbone with preservation of docking site, pose, and backbone shape - and *boosts* estimated peptide–target interface scores while doing so. This is suggestive of real benefits of utilizing non-canonicals for binder design projects, and coincides with our own experience of working with NCAAs for therapeutic peptides.

Across recovery, structural self-consistency, and deep mutational scanning, we also show that an extensible chemical vocabulary need not come at the expense of canonical design performance. AtomWeaver-open has the numerically highest observed equal-system canonical DMS agreement among the compared inverse-folding methods in both single-site and joint design, while AtomWeaver-DMS has the numerically highest observed agreement when canonical and non-canonical candidates compete head-to-head. The corresponding site-bootstrap intervals are overlapping, so these numerical orderings should not be read as statistically resolved pairwise differences. The D/*β*-aware Hirulog-3 case study further illustrates the central opportunity: atom-first generation can interrogate medicinal-chemistry choices such as stereochemistry and backbone-expanded residues from the same structural complex, rather than treating them as exceptions outside a fixed twenty-letter alphabet. AtomWeaver therefore provides a general design framework in which geometry is generated once and chemical space can be searched deliberately, at individual atom resolution.

## 7. Methods

Full details of the model architecture, joint generative formulation, nested shell source distribution, SE(3) denoiser and target conditioning, discretization and readout, training objective and schedule, sampling, evaluation, cross-modality pretraining, and data processing are given in Supplementary Methods §S1 onward. Everything required to reproduce the reported model is stated there.

## 8. Model availability

AtomWeaver is released at https://github.com/ProteinQure/atomweaver, which provides trained model weights, inference code, and the 300-residue reference library of Table S9: the residue structures and rotamers consumed by the decode-time readout. PeptideArena, the self-consistency benchmark of §4.4, is available at https://github.com/ProteinQure/peptidearena.

## 9. Acknowledgements

We thank Julie Owen for valuable feedback during the development of AtomWeaver, her careful review of this manuscript, and her insights into non-canonical amino acids and their application in peptide design. We also thank members of the ProteinQure technical team, Nicholas Coish, Parth Vora, and Gabe Szombathelyi, for supporting distributed training runs and extensive benchmarking of inverse-folding models. Finally, we thank ProteinQure’s leadership team, Lucas Siow, David White, Dave Garman, Christopher Ing, Tomáš Babej, and Mark Fingerhuth, for their support in making AtomWeaver and PeptideArena publicly available.

## Footnotes

1 A REAL slot is any slot whose element is *not* GHOST; that is, one assigned C, N, O or X, where X denotes any element other than carbon, nitrogen or oxygen.

2 https://github.com/ProteinQure/peptidearena

