## Supplementary Information for "AtomWeaver: Multi-Component Flow Matching with a Structured Geometric Prior Facilitates Non-Canonical Peptide Design"

### Supplementary Methods

#### S1 Model architecture

Figure S1 lays out the reported model. Fixed inputs (the target and the binder backbone) reach the flow along two distinct routes. A *featurization* stage (the grey band of Fig. S1) encodes them at residue level: a backbone encoder over the four backbone atoms, a target encoder that additionally carries a learned residue-type embedding, a residue-frame stream predicting $\chi_1$ and the side-chain centroid in each residue's local SE(3) frame, a volumetric stream describing the free pocket volume, and a polarity head whose element-composition prior reweights the element logits. Separately, target and backbone atoms enter the equivariant graph *directly as nodes*, carrying per-atom features that the residue-level encoders never see.

Four of those streams (target, residue frames, pocket volume and the stereochemistry head) merge into a single additive signal injected into *every* equivariant block rather than only at the input. Drawing them as one bus is not a simplification: the trunk literally adds one conditioning tensor per layer, and each source contributes through its own per-layer projection.

The trunk itself runs two tracks on one shared timestep. The atom track carries side-chain coordinates through ten equivariant blocks of 432 channels each; the element track runs an absorbing discrete diffusion from all-MASK over $\{\mathrm{GHOST}, \mathrm{C}, \mathrm{N}, \mathrm{O}, \mathrm{X}\}$. Their coupling is the FiLM junction at the end of the trunk, where the per-slot ghost/real state modulates the shared representation before *all* heads, namely velocity, element, occupancy, and mixture, so that atom count, element identity, and geometry are resolved together rather than independently.

Two feedback paths close the loop and are easy to miss in prose. The stereochemistry head takes a prior from the pocket volume and has it modulated by the evolving cloud, with a weight that is exactly zero until the midpoint of sampling and rises to one at $t = 0$; its resolved face preference then both rides the deep-inject bus and biases the velocity directly. The head is trained to *resolve* L/D handedness rather than inherit it, under the mirror and in-plane ($\hat{e}_3$-plane) corruption of §S7. The second path is an *occluded* recycle into the volumetric stream. A residue's free-pocket estimate is poorly defined without knowing where its neighbors' side-chains sit, so each residue is fed the Euler projection $\hat{\boldsymbol{x}}_0$ of its *neighbors'* predicted clean coordinates, but never its own. In training these neighbor estimates come from an explicit recycle pass (a first forward whose outputs, corrupted as in §S7, condition the packing pass); at inference the recycle is run only at the first reverse step ($t{=}T$), and every later step reuses the previous step's $\hat{\boldsymbol{x}}_0$, so recycle-like context costs a single amortized pass over the whole trajectory rather than doubling the per-step cost. Occluding a residue from its own estimate is deliberate: in training we observed that self-conditioning a residue on its own predicted coordinates tended toward belief entrenchment — the residue over-committing to an earlier guess — whereas conditioning on neighbors only, together with the input corruption of §S7, was more effective in our runs. We report this as an empirical observation, not a settled mechanism; it merits further study.

#### S2 Joint generative formulation

Let $L$ be the peptide length and $K$ the maximum side-chain atom budget per residue. Ground-truth side-chain atom positions are $\boldsymbol{x}_0 \in \mathbb{R}^{L\times K\times 3}$ and per-slot element types are $\boldsymbol{e}_0 \in \mathcal{V}_{\text{elem}}^{L\times K}$. Given binder backbone coordinates $\boldsymbol{b} \in \mathbb{R}^{L\times 4\times 3}$ (N, C$\alpha$, C, O) and a target protein structure $\mathcal{T}$, AtomWeaver models $p_\theta(\boldsymbol{x}_0, \boldsymbol{e}_0 \mid \boldsymbol{b}, \mathcal{T})$ through two coupled tracks sharing one denoiser $f_\theta$ and uses $K = 14$ at every stage, including for small-molecule pretraining.

**Element vocabulary.** AtomWeaver uses a five-symbol element vocabulary plus a MASK token: $\{\mathrm{PAD/GHOST}, \mathrm{C}, \mathrm{N}, \mathrm{O}, \mathrm{X}\}$, where PAD/GHOST denotes an absent atom and X denotes *any* non-CNO heavy atom (sulfur, phosphorus, halogens, selenium and boron all collapse to it). We resolved only those element groups with the strongest training-data support, leaving the remainder in the catch-all X class; splitting further symbols out of X is intended for future versions of the model. The consequence is a deliberate degeneracy: types distinguished only by a non-CNO element, such as sulfo-tyrosine and phospho-tyrosine, are indistinguishable on the element track and are separated, if at all, by geometry alone.

**Slot layout.** The reported model uses $K = 14$ side-chain slots per residue, ordered canonically rather than by arbitrary atom name. Slot index 0 is reserved for the atom connecting to the backbone nitrogen: this gives smoother support for $N$-methyl (and other $N$-substituted) residues such as sarcosine — their $N$-substituent occupies the dedicated slot 0 rather than displacing the rest of the side-chain — and, by absorbing that variability, keeps slot index 1 reliably C$\beta$ in meaning across residue types. Because both the geometric and learned readouts compare clouds slot-by-slot against a reference library stored in the same layout, this alignment is load-bearing: a one-position shift silently degrades the match.

**Mixture structure.** At any reverse-sampling time each slot is one of two components: *real* (an atom resolving into interface geometry) or *ghost* (an atom collapsing back to C$\alpha$). The two components follow different generative trajectories, making the process a mixture rather than a single homogeneous flow, and the shell prior gives them distinct starting contexts. Related mixture-prior formulations include mixDDPM [29], stochastic interpolants with data-dependent couplings [30], and informative prior bridges [31].

The two tracks are:

1. **Coordinates.** A continuous flow-matching process [23] in which the denoiser predicts the conditional path velocity. The conditional path is the interpolant $\boldsymbol{x}_t = (1 - s(t))\,\boldsymbol{x}_0 + s(t)\,\boldsymbol{x}_T$ with

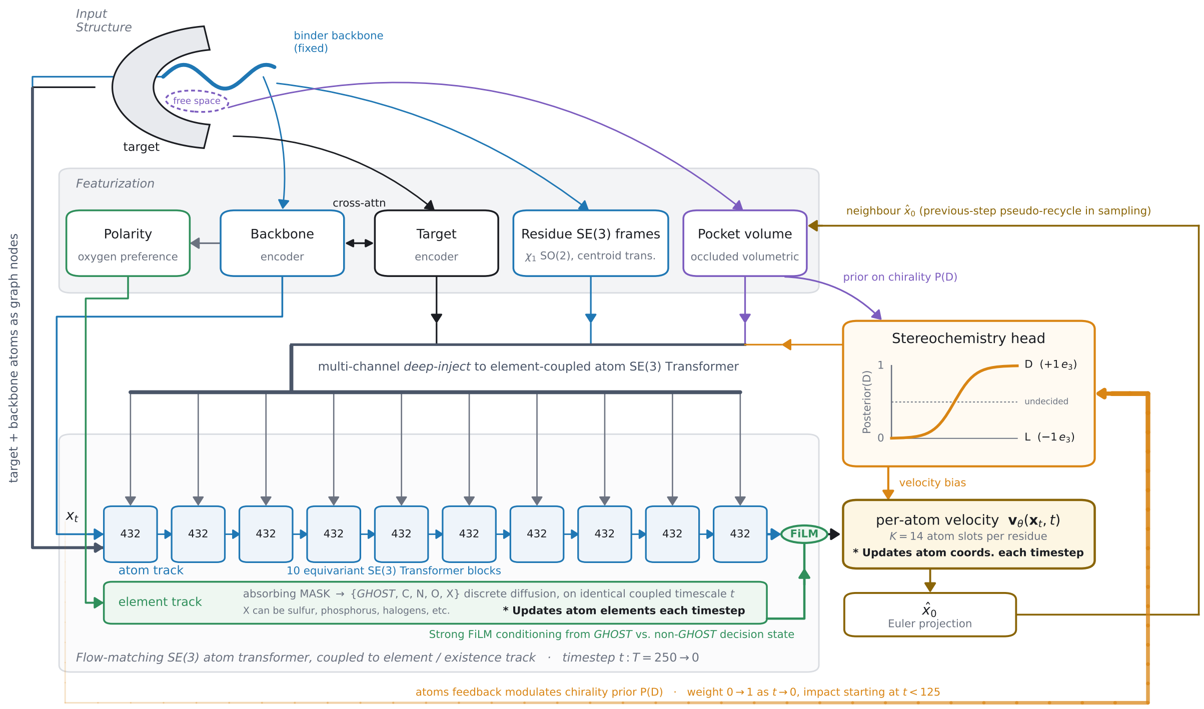


Figure S1: **AtomWeaver architecture.** Fixed inputs (top left): a target protein and the binder backbone held inside its interface, with the unfilled volume between them supplying the volumetric stream. **Featurization** (grey band) encodes these at residue level into five streams. Target and backbone atoms additionally enter the equivariant graph directly as nodes (left), bypassing the residue-level encoders. **Deep-inject**: the target, residue-frame, pocket-volume and stereochemistry streams merge into one multi-channel signal added to every one of the ten equivariant blocks, each through its own per-layer projection. **Trunk**: the atom track (432 channels, ten SE(3)-equivariant blocks) and the element track (absorbing discrete diffusion from all-MASK) advance on one shared timestep $t : T{=}250 \to 0$; at the FiLM junction the per-slot ghost/real state modulates the shared representation before all heads, which is how atom count, element identity and geometry are resolved jointly. **Stereochemistry head**: a chirality prior from the pocket (purple) is modulated by the evolving atom cloud with a weight rising from zero at the midpoint of sampling to one at $t = 0$ (amber, thickening); the resolved face preference feeds both the deep-inject bus and an out-of-plane velocity bias. Note the absence of any arrow back to the shell prior: the source distribution commits every residue to L, and handedness is discovered during generation rather than chosen up front. **Outputs**: the per-atom velocity $\boldsymbol{v}_\theta(\boldsymbol{x}_t, t)$ updates coordinates each step, while the element head resolves masked slots over $\{\mathrm{GHOST}, \mathrm{C}, \mathrm{N}, \mathrm{O}, \mathrm{X}\}$, updating atom elements each step and, because GHOST denotes an absent atom, fixing each slot's existence and hence the residue's atom count without a separate occupancy track. The velocity's Euler projection $\hat{\boldsymbol{x}}_0$ returns to the volumetric stream as the next step's neighbor context.

a per-slot power schedule $s(t) = t^{p_k}$, and the velocity target is $\dot{s}(\boldsymbol{x}_T - \boldsymbol{x}_0)$. Powers interpolate linearly across slot index from $p_{\text{prox}} = 2.5$ at the proximal end, giving more supervision near $t = 0$, to $p_{\text{dist}} = 1.0$ at the distal end. Under the shell source no ghost-specific schedule is applied, which avoids leaking ground-truth occupancy through the interpolant.

2. **Element types.** An absorbing discrete diffusion [25, 26] over the element vocabulary. The source distribution at $t = T$ is all-MASK; the reverse process predicts $p_\theta(\boldsymbol{e}_0 \mid \boldsymbol{e}_t, \boldsymbol{x}_t, \boldsymbol{b}, \mathcal{T})$ and, for slots already resolved, zeroes the MASK-class posterior and renormalizes.

**Element–velocity coupling.** The per-slot ghost/real state read from the element track modulates the shared side-chain representation through a FiLM layer [28] applied before all output heads such that a slot's coordinate velocity, element prediction and occupancy prediction are conditioned on whether that slot is currently believed to carry an atom. During training the modulation is driven by the ground-truth side-chain mask; at sampling it is a hard binary read off the element track, with unresolved MASK slots assigned a fixed intermediate value.

### S3 Nested shell source distribution

**Shell radii and thickness.** For each slot $k$ we compute the empirical per-slot C$\alpha$-to-atom radial mean $\mu_k$ and variance $\sigma_k^2$ from the training data, conditional on the slot being real, and set the shell radius to $r_k = \sqrt{\mu_k^2 + \sigma_k^2}$ and the shell thickness to the empirical per-slot radial standard deviation $\sigma_k$ itself, rather than tying it to the radius. Across the interior slots $\sigma_k$ runs 0.19–0.25 $r_k$, so neighboring shells overlap rather than partition the radial axis. At *inference* the jitter variance is scaled by 0.25, halving $\sigma_k$ relative to training. This tightens the source less than the factor suggests: per-axis source variance is $r_k^2/3 + \sigma_k^2$, and only the second term is scaled, so the realized source variance falls to ≈0.86 of its training value (per-axis width ≈0.93). Training over a wider source exposes the model to starting points further from the data, while sampling from a narrower one keeps each trajectory closer to the region it was trained to resolve.

Radii are measured per data modality: small molecule, antibody, peptides, buried protein sites; with small molecule representation differing especially from the others and being interpolated to side-chain radii gradually during that transfer learning phase in training (Table S1).

**Direction.** A single unit vector $\boldsymbol{d}$ per residue, the analytic pseudo-C$\beta$ direction computed from the residue's backbone frame, is used in both forward noising and reverse sampling, which eliminates exposure bias. All angular spread in the source comes from the jitter term $\sigma_k \boldsymbol{\eta}$; there is no separate directional-noise parameter. The pseudo-C$\beta$ construction [24] is standard in structure validation for glycines, which lack a real C$\beta$; we apply it uniformly at every position so that glycine state does not leak through the source distribution.

**Existence.** Slots within a residue sit at different shell radii, but slot $k$ is sampled from the same per-slot shell distribution regardless of that residue's true ghost/real labels.

### S4 SE(3) denoiser and target conditioning

The denoiser is an SE(3)-equivariant transformer [27] on a radius-cutoff graph over binder backbone atoms, binder side-chain slots, and nearby target protein atoms. Edges are typed as intra-residue, inter-residue (between binder residues), or binder–target, with the edge-type embedding added to the attention logit so that each class can develop distinct attention patterns. Edge cutoffs are 8 Å for intra- and inter-residue binder edges and 15 Å for edges involving target atoms. The denoiser uses 432 channels, 10 equivariant blocks, 8 attention heads, and 3 cross-attention layers.

Target conditioning enters through three independent mechanisms. **(a) Per-residue cross-attention:** a separate target encoder maps the target backbone and residue identities to per-residue features, into which binder tokens cross-attend at each block. **(b) FiLM modulation:** the target encoder's pooled global summary is mapped to per-channel scale and shift parameters that gate binder features at every block [28]. **(c) Target atoms in the graph:** target atoms participate directly as nodes in the equivariant graph, giving the attention raw geometric access to interface walls and polar contact points. Mechanisms (a) and (b) condition on residue-level chemistry; (c) is per-atom geometric conditioning.

**Auxiliary streams and deep injection.** AtomWeaver additionally carries supervised streams whose representations are injected into the equivariant trunk at *every* layer rather than only at its input, through a per-layer "deep-inject" bus in which each source contributes through its own zero-initialized residual projection, so grafting a stream onto a trained trunk does not perturb the existing solution at initialization. In the reported model the deep-inject bus carries three streams: the target encoder, the residue-frame stream (predicting side-chain orientation, $\chi_1$ and the local centroid), and the volumetric stream (predicting local side-chain density and available pocket volume from target-aware context); the stereochemistry head rides the same bus. The volumetric module is pretrained separately and *frozen* throughout fine-tuning, acting as a fixed occluded-volume conditioning stream (§3). Backbone-encoder, bond-angle and the earlier residue-frame variants are *not* deep-injected in this configuration. A per-residue chemical-composition (polarity) head reweights the element logits but is a readout head rather than a deep-inject source. We describe the configuration rather than ablating each injection, as no per-injection ablation is planned for this preprint.

These streams are residue-level in *resolution*, not in identity. That is, they do not embed a residue or a residue class; rather, they carry loosely constraining geometric summaries, including a side-chain centroid estimate that helps keep a residue's atoms from collapsing in on each other, and a $\chi_1$ rotamer state read off the backbone frame. $\chi_1$ is the first side-chain torsion, N–C$\alpha$–C$\beta$–C$\gamma$, which sets the side-chain's rotational placement about the C$\alpha$–C$\beta$ bond and indicates roughly how much room a side chain needs and which way it points. Note that under all of these loose auxiliary signals, each site remains compatible with a wide range of identities; a detailed per-residue-type embedding would not leave that freedom. What they buy is a second, coarser zoom level at which the geometry can be supervised, and their measured effect is on geometric and structural quality rather than on identity assignment.

### S5 Discretization

After reverse sampling, each residue's predicted atom cloud is discretized against a reference library. Entries store atom coordinates, per-atom element types, and a backbone alignment triple for frame alignment; canonical residues are represented by multiple rotamers. Discretization is a decode time operation: it reads identity off the generated atoms and never returns a label to the generative state, which is what keeps the residue vocabulary open-ended (§4.3). We use two readouts over the same cloud, described below.

#### S5.1 Normalized (pairwise) distance matrix readout

For the predicted side-chain we form the pairwise distance matrix over non-ghost slots, normalize it, and correlate it against the analogous matrix of each reference rotamer. The score is rotation- and translation-invariant, so the match requires no alignment, and normalization removes scale effects, leaving the match dependent on relative rather than absolute atom–atom distances. The distance-matrix score alone is retained throughout the paper as a learning-free geometric baseline; it is one of the two components of the decode-time discretizer described below.

Three mismatch penalties are applied. The *atom-count* penalty (weight 0.5) and *element* penalty (weight 0.3) are load-bearing: a count- and element-blind match inflates recovery, because a cloud with the wrong number

| Slot $k$ | Peptide | | Small molecule (pretraining) | |
|---|---|---|---|---|
| | $r_k$ (Å) | $\mu_k \pm \sigma_k$ (Å) | $r_k$ (Å) | $\mu_k \pm \sigma_k$ (Å) |
| 0 | 3.23 | $2.52 \pm 2.01$ | 1.44 | $1.43 \pm 0.16$ |
| 1 | 1.63 | $1.54 \pm 0.55$ | 2.29 | $2.25 \pm 0.40$ |
| 2 | 2.59 | $2.52 \pm 0.58$ | 2.65 | $2.62 \pm 0.41$ |
| 3 | 3.26 | $3.16 \pm 0.79$ | 3.30 | $3.24 \pm 0.63$ |
| 4 | 4.01 | $3.92 \pm 0.86$ | 3.84 | $3.77 \pm 0.72$ |
| 5 | 4.88 | $4.80 \pm 0.91$ | 4.42 | $4.34 \pm 0.84$ |
| 6 | 5.21 | $5.11 \pm 1.02$ | 4.98 | $4.88 \pm 0.97$ |
| 7 | 5.79 | $5.67 \pm 1.16$ | 5.49 | $5.39 \pm 1.06$ |
| 8 | 6.35 | $6.18 \pm 1.48$ | 6.00 | $5.88 \pm 1.17$ |
| 9 | 6.41 | $6.25 \pm 1.41$ | 6.46 | $6.34 \pm 1.24$ |
| 10 | 6.70 | $6.53 \pm 1.49$ | 6.97 | $6.84 \pm 1.33$ |
| 11 | 7.77 | $7.53 \pm 1.92$ | 7.51 | $7.37 \pm 1.43$ |
| 12 | 8.80 | $8.58 \pm 1.96$ | 8.00 | $7.85 \pm 1.50$ |
| 13 | 9.95 | $9.21 \pm 3.75$ | 8.44 | $8.30 \pm 1.52$ |

Table S1: **Empirical per-slot shell statistics, by data modality.** The quantities that define the nested shell source distribution, as calibrated for the production model: $\mu_k \pm \sigma_k$ is the C$\alpha$-to-atom radial mean and standard deviation for slot $k$ in Å, conditional on the slot being real, and $r_k = \sqrt{\mu_k^2 + \sigma_k^2}$ is the resulting shell radius. Both contexts are measured over a 4,000-window subsample at $K = 14$ slots, using the same calibration pass consumed by the model rather than a separate full-corpus statistic. *Peptide* are the shells used at inference; *small molecule* (CrossDocked, $\leq 16$ heavy atoms) is the pretraining start, from which radii are interpolated gradually toward the peptide side-chain values during transfer learning. The two differ in the way that interpolation exists to absorb: small-molecule shells are tighter and smoothly graded, growing in near-even steps with variance rising monotonically from 0.03 to 2.31 Å$^2$, whereas peptide shells are wider and markedly noisier at the top, since most residues are short canonicals that rarely reach the outer slots while every ligand carries ten to sixteen heavy atoms. Peptide slot 0 is reserved and is not the first side-chain atom, so the physical progression runs over slots 1–13; the small-molecule column does not reserve it. No slot reached the variance floor, so every value is a measurement.

of atoms can still correlate well in shape. A *chirality* penalty (weight 10 at training, 50 at evaluation) is applied because the distance matrix is mirror-invariant and cannot otherwise distinguish an L from a D side-chain. Per-reference scores are aggregated to a residue type by max-pooling over that type's rotamers.

**Reference rotamer construction.** The reference library is built from chemistry rather than lifted from deposited structures, which is what lets it carry types the training corpus never contains. Most entries (277 of 300, including all 20 canonicals) begin from the Chemical Component Dictionary ideal coordinates for the type and are rebuilt with RDKit's ETKDGv3 [35, 44] under basic-knowledge terms with chirality enforcement; a further 12 types have no CCD code and are embedded directly from SMILES by the same procedure, alongside 8 $\beta$-amino acids and 3 legacy coverage-plus types built along parallel paths. Embedded geometries are relaxed with MMFF94, hydrogens are removed afterwards since the readout scores heavy atoms only, and each conformer is trimmed to its peptide-bonded form and stored in the slot convention of §S2.

Side-chain rotamers are then drawn from observed torsion statistics rather than enumerated as conformers. Canonical wells come from a backbone-dependent library culled with PISCES [45]; each non-canonical type inherits the wells of its nearest canonical parent, mirrored for D-configured types, with marginal $\chi$ distributions as a fallback and embedded conformers for the SMILES-built types. The number kept per type is chemistry-dependent and capped at twelve. Of the 300 types, 129 reach the cap, while six with no rotatable side chain (ALA, GLY, AIB, DAL, FLA, TBG) carry a single entry, yielding 2,626 rotamers in total and a median of 10 per type. Five canonicals (ARG, ASP, HIS, LYS, TRP) additionally carry supplementary $\chi$ wells beyond the culled set.

Reference conformers are planarity-validated on the shipped library: aromatic six-membered rings are planar to a median out-of-plane deviation of 0.003 Å (p95 $\leq$ 0.011 Å), with mean aromatic C–C bond length 1.40 Å, and carboxylate groups planar to within 0.004 Å. The three coverage-plus and eight $\beta$-amino acid reference types were not independently planarity-verified; all eleven are *seen* non-canonicals, none canonical and none a zero-shot holdout. Types were retained from a larger design library only when a reference rotamer set could be built for them reliably.

At training time a soft version of this match is used as an *auxiliary discretization loss*, so the generative model is optimized not only toward the ground-truth coordinates but toward inter-atomic distance ratios that discretize correctly against the reference library. The loss is contrastive, in the InfoNCE sense: rather than a full 300-way softmax, each site is scored over a candidate subset of its ground-truth reference plus 19 decoys, and the objective is the softmax cross-entropy of picking the ground truth out of that subset. Scoring against a small set of *plausible* alternatives, rather than the whole library, is what forces discrimination from near-neighbors instead of from arbitrary chemistry.

Decoys are drawn from a clustering of the reference library by ECFP4 fingerprint similarity [46], using extended-connectivity fingerprints at radius 2 to group residues by local bonding environment. Each type therefore has a set of chemically close neighbors. An *in-cluster* decoy is one drawn from the ground-truth residue's own cluster, and is therefore the hardest kind; *out-of-cluster* decoys are spread across the remaining clusters. Decoy composition also differs by class: at a canonical site the sampler either draws all decoys from the other canonical types (a pure canonical-versus-canonical contrast, taken at a fixed probability per site) or falls through to the cluster-stratified scheme, whereas at a non-canonical site the decoys are always cluster-stratified, there being no comparably small closed alphabet to draw from. The in-cluster count is *ramped up* over training — from 2 to 8 of the 19 decoys over roughly the first half, with per-batch jitter so a fixed decoy difficulty cannot be memorized — so the discrimination problem sharpens as the flow's geometry improves. The loss uses residue-*identity* labels nowhere in the generative state: it scores geometry against reference geometry, which is why an open vocabulary survives training (§S7).

#### S5.2 Learned component

The second component is a single discriminative classifier over the 300-residue vocabulary (20 canonical + 280 non-canonical), fit once and applied at decode time only, with no retraining of the generative model and no change to what it produces. Its feature vector is 294-dimensional: 190 pairwise distances over the cloud's slots with the residue's own backbone atoms (N, C$\alpha$, C, O) entered into the same matrix, 100 per-slot element one-hots (20 slots $\times$ the five-symbol vocabulary, which also encodes each slot's existence state), and 4 backbone $\phi/\psi$ sine/cosine terms. Backbone-relative distances encode $\chi_1$, so no separate frame alignment is needed.

The four $\phi/\psi$ terms are small in number but not redundant: they are the only part of the vector that is *odd* under reflection, since $\sin\phi$ and $\sin\psi$ change sign where distances do not. They are therefore the component's entire source of handedness information, and the fit mirrors them for D-configured reference codes. Ablating them roughly halves D-recovery (0.64 $\to$ 0.31), which is why the chirality results of §4.3 depend on them. The classifier is a logistic regression.

**Self-distillation.** The learned component is trained on the *union* of (a) the model's own sampled clouds on the *training* peptides (self-distillation), covering the $\approx$ 280 types for which the training data contain examples, and which the model therefore emits when sampling — and (b) lightly-to-medium corrupted reference rotamers (atom-drop 0.1–0.2, coordinate jitter 0.3–0.6 Å), weighted by $\phi/\psi$ rotamer preference so that reference-only classes are represented by conformers appropriate to their backbone context rather than uniformly, for all 300 types — every type has a buildable reference structure — which supply the 15 trained types with no sampled clouds and the 6 held-out types (21 reference-only classes total). Those 21 are why the two counts differ. The 6 are the zero-shot holdouts, withheld from the generative model deliberately; the other 15 were kept in the vocabulary for chemical diversity when the 300-residue set was assembled, but are rare enough that the model never emitted them during self-distillation. For all 21, corrupted reference rotamers are the only signal the learned component receives. This is also why the distance-matrix component is helpful specifically in the non-canonical vocabulary (§S5.3): being a statistical score rather than a ML model, it cannot overfit to classes that have few or no sampled clouds. The self-distilled clouds are what let the learned component read the model's actual output. A logistic regression fit on clean reference rotamers alone transfers poorly to the model's noisier, sometimes under-counted clouds: it mislabels $\approx 90\%$ of canonical positions as non-canonical (right-count canonical accuracy $\approx 0.03$). This collapse is a property of the *untrained* learned classifier, not of the reference geometry itself. When the same reference rotamers are scored by distance-matrix correlation, the learning-free readout recovers 0.23 canonical top-1 overall (0.30 at the interface and 0.38 buried), showing that reference geometry carries real signal. Self-distillation is what brings the learned component up to that bar: without it the learned component sits far below the geometric readout; with it, its full-set canonical top-1 (0.24, Table 2) matches the distance-matrix score's 0.23, with a non-canonical false rate below 0.3%. Because the sampled clouds come from training peptides while the gain is measured on held-out test peptides, this is domain adaptation to the model's output distribution, not memorization of it; the corrupted-reference floor is what still lets held-out chemistry be ranked. A deliberate light touch of corruption pulls the clean reference toward the model's output distribution. It marginally improves canonical accuracy and unlocks non-zero held-out top-1 (§4.3), whereas heavy corruption only raises the false-non-canonical rate.

**Chirality gate.** The distance-matrix features are reflection-invariant, so the discretizer carries an explicit *chirality gate* that penalizes any candidate whose catalogued handedness disagrees with the sign of the cloud's pseudo-C$\beta$ signed volume, the same quantity used by the geometric chirality penalty; without it a D-configured cloud collapses onto its L mirror. The features are slot-order-sensitive, so the discretizer is fit and applied under a single slot convention (C$\beta$ first, position 0 reserved); mixing conventions between fitting and scoring silently degrades it. The distance-matrix score ranks only candidates the distance-matrix match can score, whereas the learned component ranks the full vocabulary directly.

#### S5.3 The decode-time discretizer

Every number reported in this paper comes from a single *decode-time discretizer*. It takes two distributions over the same 300 candidate classes — $p_L$, the learned component's probabilities, and $q_N$, a distribution derived from the distance-matrix scores — and mixes them per class. The two components fail in opposite regimes. The learned component is the better ranker on the canonical residues, where the empirical training-set clouds are plentiful and a fitted model has something to fit. The distance-matrix score does not attempt to fit anything: it compares distance matrices, so it treats a rare non-canonical on exactly the same footing as a common canonical, and its ranking of a residue seen twice in training is no worse than its ranking of one seen ten thousand times. That indifference is a liability where data are abundant and an asset where they are not, which is precisely where the learned component is weakest. The discretizer keeps each component in the regime it is good at.

**From distance-matrix scores to a distribution.** The raw distance-matrix scores are a correlation with penalties attached, not a calibrated distribution, and their tail is long: many chemically unrelated references still correlate moderately with a given cloud. We therefore convert them by an explicit plateau transform rather than a softmax. Candidates are sorted by score; we walk down the sorted list while each consecutive gap is at most $\varepsilon$ and stop at the first gap larger than $\varepsilon$. Everything above that break forms a near-tied *winner plateau* and shares mass $m = 0.8$ equally; everything below shares the remaining 0.2 equally. We use $\varepsilon = 0.02$ in *absolute* score units rather than as a fraction of the score range: the chirality and count penalties push the range to $\approx 58$, so a range-relative tolerance would be far too coarse to separate the top competitors. The transform is deliberately plateau-shaped — it concentrates mass on genuine near-ties and flattens the structurally-similar tail — and one consequence is that candidates in the tail receive identical mass by construction, which is why per-type percentile statistics are reported under a midrank convention.

**Per-class convex blend.** The two distributions are combined per candidate class $c$ as

$$P(c) \;\propto\; w_c\, p_L(c) \;+\; (1 - w_c)\, q_N(c), \qquad w_c = \begin{cases} 1 & c \text{ canonical} \\ 0.5 & c \text{ non-canonical}, \end{cases} \tag{4}$$

renormalized over the vocabulary, with identity read off as the argmax. The weighting is an explicit class-dependent gate, not an emergent consequence of the two score scales: canonical identity is decided by the learned component alone, while each non-canonical class is an equal mixture of the learned component and the geometric plateau.

**Why weight the two classes differently.** The learned component is the stronger ranker wherever it has empirical support, but its quality tracks how much of the model's own output distribution it has seen for a given type. On non-canonical types with sparse or entirely absent sampled clouds — exactly the types an open vocabulary exists to serve — it is fit almost entirely on corrupted reference geometry and degrades accordingly. The geometric match has no such dependence: it compares distance matrices and is indifferent to how often a type was emitted during training. Equation 4 lets geometry co-rank the non-canonicals without disturbing the canonical decision, and that is the behavior we want: canonical mass is essentially unchanged relative to the learned component alone, while non-canonical recovery improves because the geometric term surfaces candidates the learned component ranks with little evidence.

The two components are more alike than their names suggest. Both are driven principally by *pairwise atom–atom distances*; they differ in how non-geometric information enters. The distance-matrix score layers it on after the fact as explicit penalties — atom-count mismatch, element mismatch, chirality — applied to a correlation score. The learned component folds it into the feature vector instead, encoding element and existence state for every slot as one-hot features alongside the distances and fitting a single, deliberately simple logistic model over the whole set at once.

**Hand-built candidate vetoes.** The chirality gate above is a special case of a more general idea. Rather than scoring every one of the 300 candidates and letting soft penalties express implausibility, a rule can declare a candidate *impossible* for a given cloud and remove it from the ranking outright. The chirality gate does exactly this for handedness: a candidate whose catalogued configuration disagrees with the sign of the cloud's measured pseudo-C$\beta$ volume is ruled out, not merely down-weighted, because no amount of shape agreement should make a D-cloud read as its L mirror.

Other structural properties admit the same treatment, and are cheap to test from the cloud directly. A predicted cloud whose heavy atoms do not form a closed ring cannot be an aromatic or a proline analog, whatever its distance matrix correlates with. A cloud carrying no non-CNO atom cannot be a phospho- or sulfo-residue. A cloud whose atoms are not mutually bonded at plausible distances — the single disconnected atom in Fig. 2L — cannot be a branched *tert*-butyl, and a rule reading connectivity would have removed that candidate rather than ranking it first.

Vetoes of this kind are attractive because they are interpretable, need no fitting, and act where the distance-matrix representation is structurally blind rather than merely uncertain. Every site still receives an assignment: the veto narrows which candidates compete, never whether one is returned, so recovery remains a single rate over the same set of positions. We report results without vetoes beyond chirality, so the numbers here reflect an unaided geometric match; adding them is a clear avenue for removing the specific failure modes in Fig. 2(K–M).

### S6 Side-chain bond-angle deviation metric (Fig. 2B,C)

To characterize the geometry of a generated side-chain cloud independently of the identity readout, we measure how far its bond angles depart from the ideal geometry of the residue the readout assigns to it. For an assigned type we retrieve the reference conformer from the RCSB Chemical Component Dictionary (CCD) and build a graph over its heavy atoms, with bonds as edges; backbone atoms are tagged by role (N, C$\alpha$, C, O) and side-chain atoms are left generic. The generated cloud's heavy atoms are matched to this template by *element-tolerant* subgraph isomorphism (VF2), anchored on the backbone roles, so that an atom placed with the wrong element, for example a sulfur where a phosphorus belongs, still matches its template position. A residue is scored only if at least 70% of the template's atoms are matched; the rest are discarded. This keeps the angular reference well defined to the cloud positions but drops the clouds whose atom set does not correspond to the assigned identity: $\sim 24\%$ of positions overall (14% of canonical, 52% of non-canonical).

For every bond angle centered on a side-chain atom of the matched subgraph we record the absolute deviation from the corresponding ideal angle in the template. Panel B of Fig. 2 reports the median deviation as a function of topological distance (number of bonds) from C$\alpha$; panel C reports the per-residue median deviation grouped by residue class. Classes are: non-aromatic canonical (median 10°), aromatic canonical (Phe/Tyr/Trp, 21°), D-amino (12°), N-methyl (10°), aromatic non-canonical (17°), and other non-canonical (11°); histidine is grouped with the non-aromatic canonicals, its small imidazole behaving geometrically like the non-aromatic residues rather than the larger aromatic rings. Non-canonical aromaticity is detected as a planar ring in the CCD template. Applying the identical pipeline to the deposited (ground-truth) side chains gives a median deviation of $\sim 1.4°$, the native floor drawn in both panels.

Deviations grow with distance from the fixed backbone (panel B), as positional noise compounds along the chain and the training data is weighted toward proximal atoms. The two aromatic classes—canonical and non-canonical alike—are the hardest, well above the $\sim$10–12° of every non-aromatic class, so the difficulty is specific to assembling a planar aromatic ring rather than a penalty for being non-canonical.

Two caveats apply to the outermost bins. Because a cloud is scored whenever its heavy-atom topology matches the assigned residue at $\geq 70\%$ coverage, loosely placed distal atoms of long side chains are retained in the angle calculation. Those bins are also sparse: at the sixth bond the measured angles are few, and $\sim$90% of them come from a single residue ($N^6$-acetyl-lysine). The largest-distance points should therefore be read as indicative rather than quantitative.

### S7 Training objective and schedule

The total loss is a weighted sum of terms in six families. We give the families and their roles rather than a table of coefficients: the terms operate on very different natural scales, so raw coefficients are a poor guide to their relative influence.

**(1) Coordinate.** The flow-matching velocity loss on side-chain atoms, with a fill-corrected variant that accounts for slots whose existence is still unresolved. **(2) Element.** Cross-entropy over $\{\mathrm{PAD/GHOST}, \mathrm{C}, \mathrm{N}, \mathrm{O}, \mathrm{X}\}$, with PAD excluded from the target and a false-negative reweight so that real atoms are not lost to the absorbing state. **(3) Atom count.** Not one term but several — a per-residue count regression together with correlation, Pearson-$r$ and pairwise-ranking terms — weighted heavily as a family, since count fidelity rather than coordinate error is the dominant limiter of end-to-end recovery. **(4) Existence.** A per-slot occupancy term plus a low-$t$ occupancy-match guard; there is no separate existence track (§3). **(5) Discretization.** The contrastive geometric readout loss of §S5, which pushes inter-atomic distance ratios toward configurations that read out correctly against the reference library. **(6) Geometric and interfacial.** A bond and bond-angle regulariser; the residue-frame stream supervising side-chain centroid, $\chi_1$ and the stereochemistry logit; an interaction-intent classifier; a polarity / element-composition head; pocket-proximity and pocket-contact terms; and a shape prior. The volumetric stream is *frozen* — no gradient reaches its head — and contributes only self-consistency and decoy auxiliaries. A scale regulariser and a family of bonded-geometry, valence, rotamer-RMSD and mixture terms exist in the codebase but are switched *off* in the reported model.

**The coordinate loss.** The first family is the one the SE(3) denoiser is actually trained against, so we give it explicitly. Let $\boldsymbol{x}_0$ be the clean endpoint and $\boldsymbol{x}_1$ a draw from the shell source of Eq. 3, and write $\tau = t/T \in [0, 1]$. Each slot $k$ follows a linear interpolant in a slot-specific time reparameterisation,

$$\boldsymbol{x}_t^{(k)} \;=\; \big(1 - s_k\big)\, \boldsymbol{x}_0^{(k)} \;+\; s_k\, \boldsymbol{x}_1^{(k)}, \qquad s_k = \tau^{\,p_k}, \quad (5)$$

whose time derivative is the regression target,

$$\boldsymbol{v}^{(k)} \;=\; \dot{s}_k \left(\boldsymbol{x}_1^{(k)} - \boldsymbol{x}_0^{(k)}\right), \qquad \dot{s}_k = p_k\, \tau^{\,p_k - 1}. \quad (6)$$

The denoiser predicts a velocity $\boldsymbol{v}_\theta(\boldsymbol{x}_t, t, \boldsymbol{c})$ from the noised cloud, the timestep and the conditioning $\boldsymbol{c}$ (backbone, target and the per-slot GHOST/REAL state), and the loss is a occupancy-weighted squared error against Eq. 6,

$$\mathcal{L}_{\mathrm{coord}} \;=\; \mathbb{E}_{t,\,\boldsymbol{x}_0,\,\boldsymbol{x}_1}\Big[\, \sum_k w_k \,\big\|\boldsymbol{v}_\theta(\boldsymbol{x}_t, t, \boldsymbol{c})^{(k)} - \boldsymbol{v}^{(k)}\big\|^2 \Big]. \quad (7)$$

Three details are specific to this setting. The exponent $p_k$ is *per slot*, interpolated linearly by slot depth from $p = 2.5$ at the slot nearest C$\alpha$ to $p = 1.0$ at the outermost. Because $\tau^p$ is smaller for larger $p$ at any intermediate $\tau$, proximal atoms are drawn toward their clean positions earlier in the trajectory than distal ones — the same ordering seen in the geometry of the resolved clouds (Fig. 2, panel B). The endpoint $\boldsymbol{x}_0^{(k)}$ is the ground-truth coordinate for a REAL slot and C$\alpha$ for a GHOST slot, which is what makes the two-component mixture a single regression rather than two separate objectives. And $w_k$ is the *fill correction*: the element track's own $1 - P(\mathrm{PAD})$ for that slot, so a slot whose existence is still unresolved contributes in proportion to how real the model currently believes it is, rather than being counted in full or dropped outright.

**Schedules.** Three guidelines matter for reproduction: **(i)** the coordinate loss is ramped up over the later part of training (roughly doubling by the end), letting the element track stabilize before geometry is emphasized; **(ii)** the discretization loss is introduced and strengthened as training proceeds — both its weight and the difficulty of its decoys (§S5) — so that the readout objective sharpens as the flow's geometry becomes good enough to discriminate; by the final fine-tuning stage it is held at a high, fixed weight rather than still climbing; and **(iii)**

the stereochemistry t-resolution feedback is ramped in over a short warm-up and then held, while the element cross-entropy is left unramped throughout. Hyperparameter and schedule optimization for producing the best possible model is ongoing work.

**Corruption of self-referential inputs.** The neighbor-context recycle (§3) is reliable under training on reference-path samples, but not during joint sampling, where the neighbors a residue conditions on are themselves being accumulating error in reverse sampling steps and can be noisy. We therefore corrupt these self-referential inputs during training. Every corruption edits the model *input* only: the clean endpoint is left untouched and the flow-matching velocity target is rebuilt from the implied corrupted source, so the learning target stays consistent, and a low-noise gate ($\tau \geq 0.05$) disables the geometric corruptions near $t$=0, where that rebuild is ill-conditioned. Two act on the noised side-chain relative to the N–C$\alpha$–C backbone plane, whose unit normal $\hat{e}_3$ separates the L and D hemispheres: *in-plane flattening* ($p$=0.08) zeroes each real atom's out-of-plane component, producing the ambiguous "which face?" mid-state that forces the stereochemistry head to *resolve* handedness rather than copy a neighbor; and *mirror reflection* ($p$=0.10) reflects the side-chain across that plane onto the wrong (D-for-L) face, so the head must recover correct chirality from a mirror-inverted context — the sampling-time drift this guards against. The neighbor cloud itself is perturbed by randomly dropping real atoms and adding phantom ones (each $p$=0.35) and jittering coordinates ($p$=0.9, $\sigma$=0.75 Å); and the discrete track is corrupted in kind, replacing the existence mask fed to the element–velocity coupling on half of samples with the model's own three-valued read of its noised element state (real / MASK / PAD $\rightarrow$ 1.0/0.5/0.0), so atom count is trained under the mask distribution the two-track sampler actually sees rather than a leaked ground-truth one. The reported model was trained with all of the above active (only an azimuthal-rotation corruption was left off), alongside a small global input jitter (0.1 Å) and an occasional ±1–2 atom-count perturbation ($p$=0.1).

### S8 Sampling

Unless stated otherwise, sampling integrates 250 reverse steps of the coordinate flow coupled to 250 steps of the discrete element process. The coordinate track is integrated as conditional flow matching; the discrete element track uses a stochastic (DDPM-style) posterior over the absorbing vocabulary, with its sampling temperature annealed from a high value early (so identity stays fluid while geometry forms) down toward greedy near $t = 0$. A slot's existence is read directly off the resolved element track (the PAD/GHOST symbol), so atom count follows from element resolution with no separate occupancy sampler. Single-site redesign uses the replacement inpainting sampler, which pins non-designed binder residues, their element states and their coupling inputs to ground truth at every step.

### S9 Evaluation

**Recovery and geometry.** WT recovery — the fraction of designed sites at which the readout returns the deposited residue, pooled over sites rather than averaged within sequences — is scored with the atom-count and element mismatch penalties enabled. Every recovery number is reported with atom-count MAE and per-residue count correlation, and broken down by burial class (interface, buried, exposed). Exposed side-chains are under-determined by rotamer degeneracy, so exposed recovery is not optimized toward the single crystallographic rotamer.

**Co-folding.** Redesigned complexes are re-folded with OpenDDE [37] using real target MSAs and scored for pDockQ2 [40], iPTM [47], and target-aligned peptide scRMSD. For scRMSD, the target is rigidly aligned to its designed complex before peptide-backbone RMSD is evaluated; no peptide-only realignment is performed. Every comparison includes composition-scramble and random-sequence controls.

### S10 Cross-modality pretraining

AtomWeaver couples cross-modality transfer with mixed-source fine-tuning, and the transfer is productive only under the conditions noted below. Small-molecule pretraining on protein-bound ligands (CrossDocked2020 [48]) establishes an all-atom base, from which the model is fine-tuned on peptides (the PeptideMPNN training corpus [9]). Non-canonical chemistry enters not as a separate sequential pretraining stage but as single-chain NCAA-monomer data folded into the combined fine-tuning mixture (§S14), where it is trained single-site (below).

In the small-molecule stage the binder is a ligand and the shell prior requires an anchor, so we select a *pseudo-C$\alpha$* (Fig. S2): a convex-hull ligand atom whose remaining atoms fall within a narrow outward cone, mimicking the geometry of a real C$\alpha$ on a protein backbone surface, with sampling biased retained as real residues so that target conditioning trains faithfully.

Fine-tuning from a fully trained small-molecule base lowers peptide validation coordinate loss relative to a from-scratch baseline at every checkpoint we evaluated. The effect depends on the strength of the base: a weak base gives negative transfer, with the fine-tune spending early epochs unlearning misaligned priors.

Monomer data must be restricted to *single-site* NCAA windows, due to only its deeply-buried site(s) having been mutated; to many different NCAA types (§S12). Training in joint mode on monomer data overfits to the constant part of these structures, and consequently degrades buried-site peptide performance. On the other hand, single-site windows retain NCAA exposure without that cost: thus, monomer data serves as an NCAA-chemistry supplement rather than a second peptide corpus.

### S11 Data and data processing

**Peptide corpus.** This consists of real peptide:target interfaces, a subset of the PeptideMPNN-curated

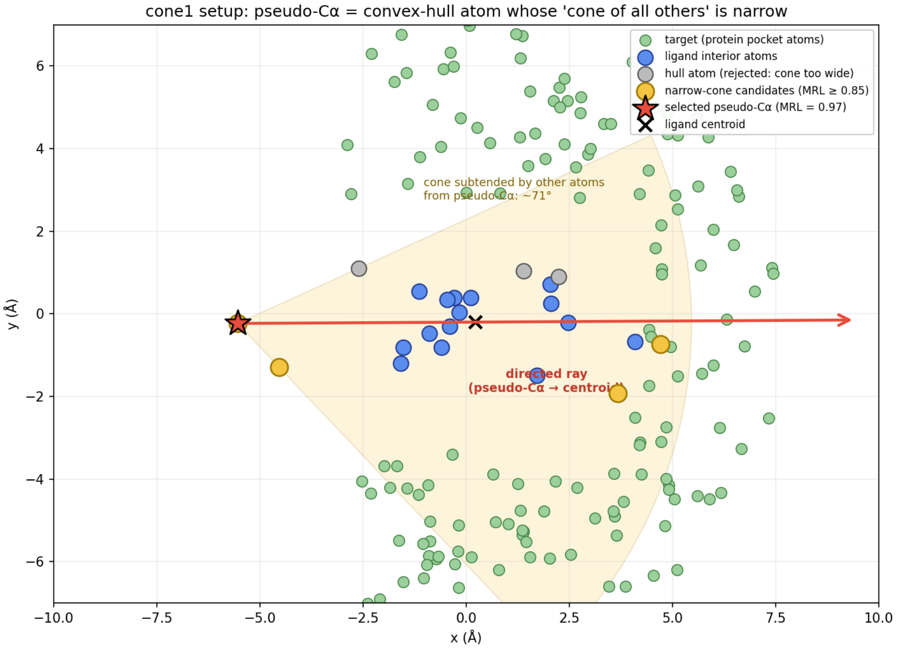


Figure S2: **Giving a small molecule a backbone-like anchor, so the same shell prior applies.** The nested shell prior needs two things a ligand does not have: an *origin* to measure radii from, and a *direction* to spread the shells along (Eq. 3). On a peptide these are the residue's C$\alpha$ and its pseudo-C$\beta$ ray; a small molecule has neither. We therefore construct both. The origin is a *pseudo-C$\alpha$*: an atom on the ligand's convex hull chosen so that all remaining ligand atoms subtend a *narrow* cone from it (here $\approx 71°$; red star, selected from the yellow candidates, with grey hull atoms rejected for too wide a cone). The direction $\boldsymbol{d}$ is then the ray from that anchor to the ligand centroid (red arrow). The narrow-cone criterion is what makes the construction work rather than arbitrary: a real C$\alpha$ sits on the protein surface with its side-chain projecting away from the backbone into one restricted solid angle, so an anchor whose molecule hangs off it in a single direction presents the flow with the same local geometry it will meet on a peptide — atoms at increasing radii along one outward ray. Chosen this way, the small-molecule stage and the peptide stage pose the same problem in the same coordinates, which is what makes the pretraining transfer (§S10); an anchor picked without the cone criterion would scatter the ligand's atoms around the origin and teach a prior the peptide model cannot use. Green atoms are the surrounding protein pocket, blue the ligand interior.

dataset [9] with peptide length 5–32 residues, to which we added 548 further experimentally determined peptide:protein complexes carrying non-canonical residues, taken from crystal coordinates without modeling. The two are treated as one corpus of real interfaces throughout. We used the exact same PDB-cluster level split as [9], taking 1000 complexes as a cluster-based holdout. Clustering was based on the peptide sequence only, therefore identical receptors may exist in both training and holdout. We ensured that none of the additional complexes overlapped with the holdout. Additionally, we filtered out structures that contain the holdout NCAAs (MK8, 2MR, KCR, OMY, CGU, MP8), resulting in a training set of 11,072 complexes. Out of these, 2,939 (26.5%) have at least one NCAA of the 520 unique NCAAs in the peptide chain.

**Small molecules.** Positive (active-site) ligand–receptor pairs from CrossDocked2020 [48], with receptors preserved as real residues for target conditioning and ligands filtered to the atom budget ($K = 14$).

**Reference library.** A library of residue templates and rotamers underlies both the geometric readout and the corrupted-reference half of the learned component. For reporting we fix a 300-*residue vocabulary* — the 20 canonical residues plus 280 non-canonical types with sufficient structural support — of which 6 (MK8, 2MR, KCR, OMY, CGU, MP8) are withheld: absent from the generative model's training corpus and training-time library and from the readout's self-distillation, so recovery on them is zero-shot. They remain scorable because the readout carries a reference entry for every vocabulary type.

The generative model sees more non-canonical chemistry than the readout vocabulary contains: training draws on the 455 NCAA types of §S12, whereas the reported vocabulary is the polished 300. The two differ for practical rather than principled reasons — some types could not be given a reliable rotamer set, and others were dropped because their structures were not distinguishable from another code's — and nothing about the method fixes the number. A user can add or remove any canonical or non-canonical type and refit the readout in minutes, without retraining the generative model (§S5.3).

### S12 Synthetic non-canonical amino acid data

The sparsity of experimental structures containing non-canonical residues presents a major hurdle for any data-based generative method. We therefore supplemented the corpus of §S10 with a heavily filtered synthetic data set. In brief, two complementary physicsbased methods were used to introduce a single NCAA into otherwise all-canonical experimental template structures. In the bulk of this set the NCAA was mutated into fully or partially buried positions within monomeric proteins; we considered this the lower-risk case for synthetic data, since better-packed positions inherently provide more quality-filtering information. A smaller fine-tuning set was generated at interfacial positions of peptides complexed to targets, and was also stringently filtered.

**Synthetic NCAA structure generation.** Template structures for NCAA mutation were curated from the RCSB PDB [49] (single-chain, resolution $\leq 2.0$ Å, protein-only; ligands and other non-solvent heteroatoms excluded, crystallographic waters retained). Secondary structure was annotated with DSSP [50] and relative solvent accessibility computed with a Shrake–Rupley algorithm [51], and disulfides were resolved from records plus S$\gamma$–S$\gamma$ distance checks. NCAA mutation positions were selected by burial (relative SASA and heavy-atom neighbor count) and backbone region, spanning buried, intermediate and exposed environments. During generation a single native side chain was replaced by a NCAA on the retained backbone, and the structure was minimized by one of two complementary back-ends: an internal build assigning Amber ff14SB [52] and GAFF2 [53] parameters through AmberTools [54] and relaxing the local shell (steepest descent then conjugate gradient) in `sander`, or a MOE build under AMBER10:EHT [5]. NCAA coverage is bounded by the chemistries the back-end can parameterize.

**Quality control.** A successfully parameterized and minimized candidate structure was retained as a training example only if it passed a logical-AND of checks against its minimized wild-type counterpart: below-threshold side-chain strain and backbone RMSD (local-shell and mutated-residue), and an energy gate (a per-NCAA energy-residual outlier cut on the internal track; a $\Delta$E and per-NCAA-median signed-delta cut on the MOE track). Charged side-chains in buried environments additionally faced a desolvation veto on the MOE and dimer tracks.

**Composition.** The pipeline delivered ≈146,000 quality-passed all-atom structures over 455 distinct NCAA types[3] (∼90% of a 503-type reference corpus): the MOE track sampled chemistry broadly (24,878 structures, 418 types) and the in-house method more deeply (117,413 structures, 223 types), with a dimer stream extending the build to peptide:protein interfaces (4,157 structures, 80 types; the NCAA installed on the peptide chain and gated additionally on interface integrity). As a further complement to the experimental PeptideMPNN data set (vide supra), which contains some, but limited, NCAA examples, we surveyed the PDB for additional high-quality NCAA-containing structures. This second batch of experimental, "native" NCAA poses was assembled directly from crystal coordinates without modeling (monomer 347 structures, 77 types; dimer interface 680 structures, 233 types).[4] Training and evaluation are split at the whole-structure level (fixed seed, ≈20% held out); the installed-NCAA vocabulary, the withheld zero-shot set, and the training mixtures (with the canonical peptide sources) are described in §S11.

### S13 Antibody CDR loop data

**SAbDab CDR pseudo-peptide dataset generation.** Antibody–antigen structures were curated from SAbDab [55] (downloaded 2 July 2026). Only entries with a peptide or protein antigen and with resolution better than 3.5 Å were kept, and complexes involving MHC were excluded. Antibody and antigen sequences were clustered separately at 95% sequence identity with 80% coverage, and one representative was selected for each antibody–antigen cluster pair, prioritizing resolution and *R*-free value.

Structures were downloaded from the RCSB PDB [49], and antibody chains were renumbered according to the IMGT scheme with ANARCI [56]. Protein atoms were retained and non-protein components — water, ions, glycans, small molecules and nucleotides — were removed. CDR residues were defined by IMGT positions 27–38, 56–65 and 105–117 for CDR1, CDR2 and CDR3 respectively. Structures were retained only when at least one CDR heavy atom fell within 4.5 Å of a protein antigen heavy atom.

To construct peptide-like training examples, each antigen-contacting CDR loop was treated as an independent pseudo-peptide binder, with the remainder of the antibody structure removed. Loops shorter than 3 residues or longer than 32 residues were excluded. Applying these criteria yielded 21,055 CDR pseudo-peptide complexes.

### S14 Training

The reported model is produced by a staged curriculum of weights-only continuations (each resumes the previous stage's weights and resets its epoch counter), which we summarize by data regime. Training passes through roughly four data stages: a small-molecule pretrain (≈500 epochs on protein-bound ligands, §S10); a first peptide stage (≈100 epochs) that transfers the atom flow into peptide side-chain context; a combined multi-source stage (≈280 epochs) that folds in non-canonical chemistry and structural diversity; and a final peptide consolidation (several hundred further epochs across the corruption and discretization fine-tunes above) that yields the reported model. The first two, as well as the final peptide stage, ran on two RTX 5090 GPUs, and the combined multi-source stage on a cluster of fourteen.

The stages differ mainly in how many binder residues are masked for design at once — the "K-mix" — and

[3] The 455 synthetic types exceed the polished 300-residue rotamer database used at decode time. Types outside that database carry no discretization loss during training — they have no reference entry to be scored against — but they are retained because they still supply the coordinate-side objectives: the additional chemistry broadens the structural diversity the atom flow is exposed to, which is what those losses are trained on.

[4] Native type counts are by PDB chemical-component code; synthetic type counts are by internal NCAA identifier, so the two are not on the same basis.

the guiding principle is that *real peptide interfaces are trained full-joint, synthetic NCAA data single-site.* (i) Small-molecule pretraining applies no K-masking: the whole ligand cloud is packed at once. (ii) The peptide stages use a *reversed-K* schedule that begins fully joint (every residue designed simultaneously, $K{=}L$) and only later relaxes to a mixture of full-joint ($K{=}L$), an intermediate site count ($K \sim \text{Uniform}\{2, \ldots, L{-}1\}$), and single-site ($K{=}1$) draws — weighted 0.55/0.25/0.20 in the reported model. Beginning joint and introducing single-site last was the fastest route to transfer from the small-molecule base to full-peptide side-chain packing. (iii) The synthetic non-canonical sources (an NCAA-monomer set windowed to 8–14 residues, and synthetic NCAA dimers) are instead trained heavily *single-site-biased* — a geometric $K$-draw concentrated on $K{=}1$ — because a synthetic non-canonical example supplies only one genuinely designed residue, and full-joint design over such scaffold-poor data overfits to the recurring non-designed positions. Meanwhile, the peptide and antibody sources already carry the joint-mode signal. The combined stage mixes these sources at roughly 47% peptide, 25% NCAA-monomer, 14% synthetic NCAA-dimer, 9% antibody and 4% native complexes; the final consolidation drops back to peptides plus native dimers under the reversed-K schedule.

The final fine-tune ran for 100 epochs under PyTorch Lightning distributed data-parallel, with a per-GPU batch of 1 and no gradient accumulation (effective batch 2); activation checkpointing (stride 2) holds a full-length peptide and its target in memory at single precision. The optimizer is AdamW at a fixed fine-tuning learning rate of $2 \times 10^{-5}$, warmed up linearly over 1000 steps and then cosine-annealed to $2 \times 10^{-6}$ on a per-step schedule; weights are restored from the base checkpoint with the optimizer state reset — a deliberate plateau-breaking restart. Fine-tuning data are peptide and dimer binders only (no non-canonical-monomer mixing at this stage). A family of non-canonical types is withheld from training at the binder level — any binder containing one is dropped, so the residue is never seen in a peptide context — and the six of these that are representable in the readout vocabulary constitute the reported zero-shot leg-D set (§4.3).

#### Supplementary Results

### S15 Extended non-canonical recovery results

**Reconstruction-quality metrics: matching and denominators.** Table 1 reports the flow-side quantities. Predicted and ground-truth side-chain atoms are matched by optimal (Hungarian) assignment on C$\alpha$-relative distance; RMSD is post-Kabsch on the matched pairs and matched element accuracy is the fraction of matched atoms sharing an element, whereas composition overlap is an order-free element-histogram overlap (assignment-independent). *Denominators differ by metric.* RMSD, matched element accuracy and composition overlap are over $n$=180,354 residues (scored positions with a non-empty matched cloud); per-residue count MAE and correlation over $n$=199,350 designed positions, which include glycine at a ground-truth count of zero; GHOST/REAL distances over the 14 slots of each of 198,830 residues.

$\chi_1$ is the first side-chain torsion (N–C$\alpha$–C$\beta$–C$\gamma$). Deviations are circular, so we report the circular median of $|\Delta\chi_1|$ and the fraction within 20°, over the scored positions further restricted to residues where $\chi_1$ is defined ($n$=146,243): excluded are GLY and ALA (22,970), a ground truth lacking C$\beta$/C$\gamma$ (2,810), and a non-canonical ground-truth identity (2,530). C$\beta$ and C$\gamma$ are the predicted atoms the Hungarian assignment pairs to them, not fixed slots, so $\chi_1$ uses the same correspondence as the RMSD and element rows. A further 24,797 positions are excluded because the assignment reached no predicted atom for the ground-truth C$\gamma$; this row is therefore conditioned on the model having placed a $\gamma$ atom, and should be read alongside the count metrics rather than independently of them. $\chi_1$ is the first and most constrained side-chain torsion, so it is the most favorable rotamer measurement; torsions further out are not reported.

**Dynamic stereochemical discovery.** Having established that canonical recovery is well-behaved (§4.2), we turn to the non-canonical residues, beginning with the broad machinery: does the model resolve *stereochemistry* at all? It does. Distance-matrix matching is mirror-invariant, so handedness is read from the predicted cloud, and every site was initialized in the L-chiral direction at $t = T$ with no ground-truth chirality information supplied (Fig S3); from that purely L-initialized start the model recovers D at 69.6% of the scorable D-configured positions (415 of 596; positions that collapse to an achiral atom count are unscored), while mislabeling only 2.9% of the L-chiral residues as D (97.1% specificity; Fig S3a, inset). Glycine is achiral — no C$\beta$, hence no handedness — and is excluded from the D-vs-L evaluation (it is signed at chance whenever a spurious side-chain atom is placed, so lumping it with L would overstate the L-chiral error rate). The supervised stereochemistry head steers the atoms toward the resolved face during sampling, and the handedness read from the predicted cloud's geometry separates D from L-chiral at ROC-AUC 0.869 (Fig S3a), so the call is not made by indiscriminately assigning D. The correctly recovered D sites fall in the mirror-image ($+\phi$) basin of the Ramachandran map — where D-residues belong and canonical L-residues almost never sit; the false-D predictions instead concentrate in the strongly-L $\alpha/\beta$ basins and are largely genuine chirality errors, with only ≈26% landing in the mirror-of-$\alpha_L$ pocket where D is geometrically permissible (Fig S3d,e). Tracking the per-step geometry resolves the mechanism (Fig S3b,c): at D sites the side-chain crosses the backbone mirror plane and locks onto the D face, earlier for bulkier residues; at canonical L sites it stays L, false-D calls being rare (13/282) and confined to residues whose side-chain collapses to a single atom — a read-out degeneracy, not a stereochemical error.

**Dynamic non-canonical recovery.** We then turn to detailed per-type recovery — again sampled jointly (full joint peptide de novo) and measured on the 1k held-out test set. At positions whose native residue is non-canonical, the model surfaces seen non-canonical types near the top of the 300-residue vocabulary without being directed to look for them (median percentile 95.7, 47.0% in the top ten). The six withheld types — unseen by both the generative model and the readout's self-distillation, and carried by reference geometry alone — are the hardest category we report, and their numbers (median percentile ≈ 63) should be read as a floor on *our* per-type results rather than as an estimate of what zero-shot recovery can necessarily reach in principle; KCR is the standout, reaching the top ten at 0.45. Recovery varies with residue size and distinctiveness (Supplementary Table S2). The readout recovers a substantial set at top-1 (HYP, PTR, DAL, M3L) and localizes the phospho/sulfo modifications and several D-types at top-3/top-10 (TYS, PTR, TPO, DAL, B3L); among the zero-shot holdouts, MK8, 2MR and KCR are all represented within the top ten. Opening the vocabulary from 20 to 300 types costs essentially nothing on the canonicals. Top-1 in an open 300-residue match is low for most non-canonical classes, so we report rank alongside argmax; since much of the identity signal routes through atom count, a count-matched non-canonical can rank well for size-driven reasons. D-amino acids are resolved by geometry (Fig. S3) and recovered rather than suppressed — D-Ala, for instance, reaches top-1 0.34 and top-10 0.71.

The discretizer readout spans the full canonical alphabet and broadly tracks the ground-truth composition (Fig. S4), over-assigning the aggregate non-canonical bin relative to its 1.3% native rate. That over-assignment is correctable at decode time without touching the model: multiplying in a natural-frequency prior over the canonical classes pulls the non-canonical share back below the native rate (Fig. S4). We report the uncorrected readout throughout, since a prior fitted to natural frequencies is the wrong instrument for a method whose purpose is proposing residues that natural frequencies do not cover; the comparison is shown here only to locate the readout against that baseline. No configuration reproduces the ground-truth composition exactly, so what this supports is coverage of the alphabet rather than agreement with it.

### S16 The full PeptideArena target set

Table S3 provides details on the targets selected for PeptideArena and the number of de novo backbones included in the evaluation set. Figure S6 reports every target, of which the main text shows the six with the highest average success rate across models. Designs clearing 5 Å but not 2 Å recover the docking site and the general binding pose. We prioritize the 5 Å threshold because exact backbone recovery could not be easily disentangled from the fact that peptides are generally floppier and less rigid.

### S17 Deep mutational scan: benchmark construction and evaluation

**Candidate curation.** CAA+NCAA evaluates the 20 canonical amino acids together with the 21 non-canonical amino acid identities assayed in the DMS scan. Each identity has a corresponding reference structure, including Bzt, tert-butyl-L-alanine (tBu), O-methylhomoserine (hSM), and *N*-methyl-2-aminobutyric acid (MeB), and is scoreable at all 39 benchmark sites in both single-site and joint-design evaluations. In the reference vocabulary these four are represented as 4OG, 0JY, Z36 and Z37 respectively (Supplementary Table S9). At each site, we exclude the wild-type identity of the assayed construct from the substitution set, rather than the residue observed in the deposited structure; this distinction matters at construct–structure mismatch sites.

**Baseline protocols.** For joint design, ProteinMPNN and PeptideMPNN construct multi-site sequences through ordered autoregressive conditionals using random orders, ESM-IF uses a left-to-right autoregressive order, and FAMPNN uses its native progressive-unmasking sampler. LM-Design uses the official ByProt inpainting mode, with the full peptide chain masked as one segment and the target chain held visible, followed by its default five denoising iterations. These model-native protocols differ from AtomWeaver's simultaneous joint conditioning. The primary joint panel uses 100 whole-peptide draws at sampling temperature $T$=1.0; a matched sensitivity panel repeats these protocols at $T$=0.2. Temperature controls these stochastic joint-design samplers. Single-site baselines instead read their direct conditional distributions at each queried position, with all other peptide positions fixed, so they need not sample a sequence. A separate logit-temperature scaling could be applied as probability calibration, but it is not part of this model-native single-site protocol; the rank-based statistic is invariant to positive scaling. Canonical-label baselines are evaluated only in CAA-only; NCFlow is a reference-only affinity-scoring comparator, not an inverse-folding model.

**Benchmark arms, sampling, and readout.** We evaluate single-site and joint design separately under two candidate-set views: CAA-only (canonical amino acids) and CAA+NCAA (canonical and non-canonical amino acids competing jointly). At each site, the assay/construct wild-type identity is excluded from its own substitution candidate set. CAA-only therefore contains 19 candidates per PUMA site and 18 per CP2 site; CAA+NCAA additionally includes all 21 non-canonical identities, yielding 40 and 39 candidates per site, respectively. We do not report an NCAA-only benchmark since it is a contrived benchmark given that several positions prefer canonical over non-canonical substitutions.

Each AtomWeaver condition retains 100 stored atom-cloud draws per benchmark position. The primary sampled joint-design baseline panel likewise uses 100 stored whole-peptide draws per system at sampling temperature $T$=1.0; a matched sensitivity panel repeats the same procedures with 100 draws at $T$=0.2. Single-site baseline evaluations instead read the native direct conditional-score distribution at each queried position, with no sampled sequence or generative draw axis. We do not apply a separate logit-temperature calibration; for the primary Spearman metric, any positive temperature scaling would leave candidate order unchanged. Within a design mode, AtomWeaver-open, AtomWeaver-

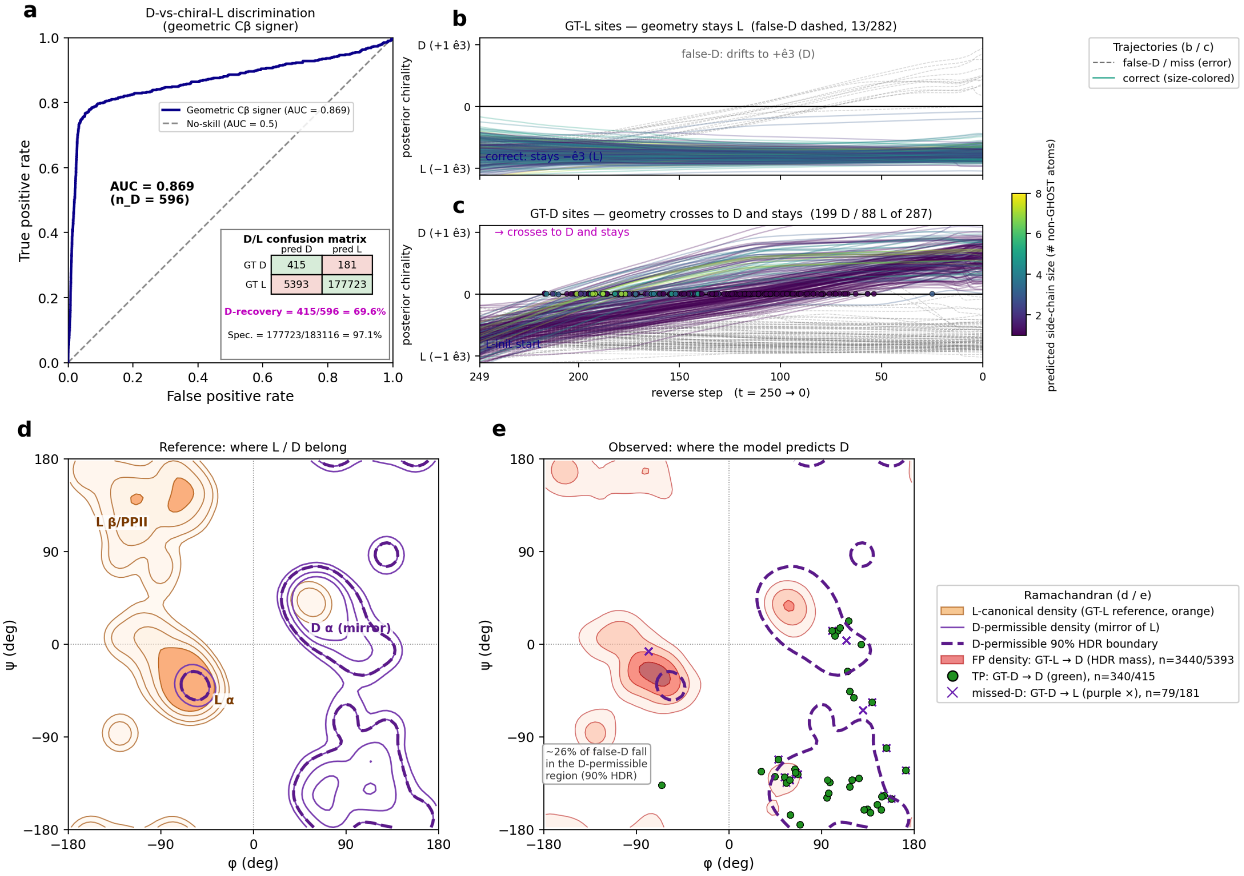


Figure S3: **Stereochemistry is resolved from geometry under a blind L-initialized start.** Every residue position is initialized in the L-chiral direction at $t = T$ with no ground-truth chirality supplied; handedness is read from the predicted cloud as the sign of the C$\beta$ out-of-plane projection ($+\hat{\mathbf{e}}_3$ = D face), and ground-truth handedness is assigned per residue from geometry (the signed N–C$\alpha$–C / C$\beta$ triple product), so mislabeled deposits resolve automatically (e.g. the geometrically-L DPN in 1ABI is scored L; chain-terminal canonical backbones that are geometrically D are counted as D) with no per-residue special-casing. Panel (a) is computed over the full 1k held-out test set plus the leg-D holdout structures (10 designs per peptide); panels (d,e) require a Ramachandran $(\phi, \psi)$ coordinate and so use the subset of those positions with both backbone dihedrals defined, dropping chain-terminal residues. (per-class $n$ in the legend). **(a)** ROC for D-vs-L-chiral discrimination from the geometric read ($n_{\mathrm{D}} = 596$, $n_{\mathrm{L}} = 183{,}116$; ROC-AUC $= 0.869$; achiral glycine and chirality-undetermined positions excluded). *Inset:* the D/L confusion matrix (achiral row/column omitted) — the model recovers D at 415/596 = 69.6% of signed D positions at a 2.9% L-chiral false-D rate (Spec. 97.1%). **(b,c)** Per-reverse-step posterior chirality (signed C$\beta$ out-of-plane projection, $+\hat{\mathbf{e}}_3$ = D) along the $t$:250 $\to$ 0 trajectory, on an *illustrative curated subset* (a spectrum of D-type residues; 282 ground-truth L and 287 ground-truth D site-trajectories from the same model, with per-step dumps), colored by predicted side-chain atom count; errors dashed grey. **(b)** Ground-truth L sites: 269/282 correctly stay on the L face; the 13 false-D (grey) are rare and arise late from single-atom read-out degeneracy, not the chirality head. **(c)** Ground-truth D sites: the side-chain crosses the mirror plane and locks onto the D face, earlier for bulkier residues; the supervised stereochemistry head steers this crossing but is itself a high-noise driver that hands off before $t = 0$, the committed geometry being the resolved handedness. **(d,e)** Ramachandran $(\phi, \psi)$ view, shared axes. **(d)** *Reference regions:* the empirical L-canonical density (orange), computed from our peptide-interface data, and the D-permissible density (purple), its exact mirror $D(\phi, \psi) = L(-\phi, -\psi)$ — a D residue is permissible wherever the mirror-image L conformation is populated; the purple-dashed contour is the 90% highest-density region (HDR: the smallest $\phi/\psi$ area containing 90% of the mirrored density). Basins are labeled L-$\alpha$, L-$\beta$/PPII and D-$\alpha$. The small purple pocket *inside* the orange L-$\alpha$ basin is the mirror of the minor $\alpha_L$ (left-handed helix) basin near $(+60, +45)$, which reflects to $(-60, -45)$ and so falls within L-$\alpha$ — the sole place the L and D-permissible zones overlap. **(e)** *Model D-calls:* where the model actually predicts D, over the same D-permissible 90% HDR boundary — false-D (predicted D, GT-L) as a red density (shading = HDR mass), correctly-recovered D (predicted D, GT-D) as green points, and missed-D (GT-D, predicted L) as purple $\times$. The false-D concentrate in the strongly-L $\alpha/\beta$ basins (genuine chirality errors); only $\approx$26% of the ($\phi/\psi$-defined) false-D fall in the D-permissible region (890/3440) — precisely the $\alpha_L$-mirror overlap of (d) — while the correctly-called D sit in the positive-$\phi$ D-permissible zone.

| CCD | Chemistry | Set | $n$ | sites | t1 | t3 | t10 | pctGT |
|---|---|---|---|---|---|---|---|---|
| HYP | 4-hydroxy-Pro | seen | 390 | 39 | 0.44 | 0.76 | 0.79 | 97.1 |
| SEP | phospho-Ser | seen | 340 | 34 | 0.14 | 0.30 | 0.44 | 92.9 |
| TPO | phospho-Thr | seen | 110 | 11 | 0.19 | 0.30 | 0.34 | 87.2 |
| ALY | acetyl-Lys | seen | 100 | 10 | 0.20 | 0.24 | 0.31 | 80.4 |
| PTR | phospho-Tyr | seen | 100 | 10 | 0.60 | 0.60 | 0.60 | 73.9 |
| DAL | D-Ala | seen | 70 | 7 | 0.34 | 0.36 | 0.71 | 97.2 |
| 2GX | aromatic | seen | 60 | 6 | 0.20 | 0.23 | 0.37 | 68.9 |
| M3L | trimethyl-Lys | seen | 60 | 6 | 0.28 | 0.40 | 0.73 | 97.8 |
| TYS | sulfo-Tyr | seen | 50 | 5 | 0.04 | 1.00 | 1.00 | 99.3 |
| MAA | N-methyl-Ala | seen | 40 | 4 | 0.00 | 0.00 | 0.00 | 79.9 |
| ORN | ornithine | seen | 40 | 4 | 0.17 | 0.20 | 0.20 | 93.7 |
| 1MH | aromatic | seen | 30 | 3 | 0.00 | 0.00 | 0.13 | 85.1 |
| ASA | Asp-analog | seen | 30 | 3 | 0.00 | 0.00 | 0.30 | 92.5 |
| B3L | $\beta$-homo-Leu | seen | 30 | 3 | 0.20 | 0.40 | 0.90 | 97.2 |
| HLX | homo-Leu | seen | 30 | 3 | 0.00 | 0.00 | 0.00 | 84.7 |
| PRS | aliphatic | seen | 30 | 3 | 0.00 | 0.00 | 0.00 | 29.1 |
| 41H | aromatic | seen | 20 | 2 | 0.00 | 0.00 | 0.00 | 47.3 |
| ALC | aliphatic | seen | 20 | 2 | 0.00 | 0.00 | 0.40 | 84.9 |
| CHG | cyclohexyl-Gly | seen | 20 | 2 | 0.00 | 0.00 | 0.00 | 88.5 |
| DGL | D-Glu | seen | 20 | 2 | 0.00 | 0.00 | 0.00 | 88.4 |
| DPN | D-Phe | seen | 20 | 2 | 0.05 | 0.75 | 0.85 | 98.6 |
| HIX | aromatic | seen | 20 | 2 | 0.00 | 0.00 | 0.00 | 78.6 |
| HT7 | D-aromatic | seen | 20 | 2 | 0.10 | 0.10 | 0.95 | 97.8 |
| LE1 | aliphatic | seen | 20 | 2 | 0.00 | 0.00 | 0.00 | 70.8 |
| MEA | N-methyl-Phe | seen | 20 | 2 | 0.00 | 0.00 | 0.00 | 64.8 |
| MLE | N-methyl-Leu | seen | 20 | 2 | 0.00 | 0.30 | 0.50 | 93.5 |
| MLY | dimethyl-Lys | seen | 20 | 2 | 0.00 | 0.20 | 0.50 | 88.4 |
| NLE | norleucine | seen | 20 | 2 | 0.00 | 0.50 | 0.50 | 88.4 |
| NVA | norvaline | seen | 20 | 2 | 0.00 | 0.00 | 0.00 | 66.0 |
| PCA | pyroglutamate | seen | 20 | 2 | 0.00 | 0.00 | 0.00 | 77.2 |
| SAR | N-methyl-Gly | seen | 20 | 2 | 0.00 | 0.00 | 0.85 | 96.2 |
| SC2 | aliphatic | seen | 20 | 2 | 0.00 | 0.00 | 0.00 | 85.2 |
| 0BN | aromatic | seen | 10 | 1 | 0.00 | 0.00 | 0.00 | 20.3 |
| AIB | $\alpha$-methyl-Ala | seen | 10 | 1 | 0.00 | 0.00 | 0.00 | 73.7 |
| CIR | citrulline | seen | 10 | 1 | 0.20 | 0.70 | 0.80 | 98.7 |
| CSO | oxidized-Cys | seen | 10 | 1 | 0.00 | 0.00 | 0.00 | 11.5 |
| DPR | D-Pro | seen | 10 | 1 | 0.00 | 0.20 | 0.80 | 97.3 |
| DTY | D-Tyr | seen | 10 | 1 | 0.00 | 0.00 | 0.00 | 87.6 |
| FP9 | fluoro-Pro | seen | 10 | 1 | 0.00 | 0.00 | 0.20 | 83.4 |
| HMR | $\beta$-homo-Arg | seen | 10 | 1 | 0.00 | 0.00 | 0.60 | 96.9 |
| IYR | iodo-Tyr | seen | 10 | 1 | 0.00 | 0.00 | 0.00 | 91.4 |
| MVA | N-methyl-Val | seen | 10 | 1 | 0.00 | 0.00 | 0.40 | 93.4 |
| NAL | naphthyl-Ala | seen | 10 | 1 | 0.00 | 0.30 | 0.50 | 94.6 |
| NMM | methyl-Arg | seen | 10 | 1 | 0.00 | 0.00 | 0.20 | 62.0 |
| ORQ | Orn-analog | seen | 10 | 1 | 0.00 | 0.00 | 0.00 | 55.1 |
| YNM | aromatic | seen | 10 | 1 | 0.00 | 0.00 | 0.00 | 66.5 |
| MK8 | $\alpha$-Me-norleucine | held | 170 | 17 | 0.00 | 0.04 | 0.05 | 72.6 |
| 2MR | sym. dimethyl-Arg | held | 110 | 11 | 0.03 | 0.03 | 0.12 | 57.2 |
| MP8 | 4-methyl-Pro | held | 100 | 10 | 0.00 | 0.00 | 0.00 | 58.0 |
| OMY | chloro-Tyr | held | 80 | 8 | 0.00 | 0.00 | 0.00 | 44.9 |
| KCR | $N^6$-crotonyl-Lys | held | 40 | 4 | 0.00 | 0.20 | 0.45 | 72.2 |
| CGU | $\gamma$-carboxy-Glu | held | 10 | 1 | 0.00 | 0.00 | 0.00 | 51.1 |

Table S2: **Per-type non-canonical recovery over the 300-residue vocabulary under full-joint design**, for every non-canonical type with $n \geq 10$ (at least one site) in the 1k held-out test set — all 46 seen types — plus the six zero-shot holdouts. Columns are top-1/top-3/top-10 recovery and the ground-truth percentile ("pctGT": where the ground-truth residue falls in the readout's ranking of all 300 candidates, as a percentile, so 50 is chance and 100 means it was ranked first), with $n$ the number of sampled positions (10× the distinct sites). *Seen* types are scored in the 1k held-out test set; *held* types are the six zero-shot holdouts, carried by reference geometry alone and scored at real peptide:target interfaces (§S11), those being the only places a held-out type occurs in an experimentally determined complex. *TYS/PTR caveat:* sulfo-Tyr (TYS) and phospho-Tyr (PTR) are vocab5-degenerate (both S and P map to the aggregate X element), so the rank-1 split between them is not meaningful — the readout assigns rank-1 to PTR with TYS at rank 2–3 (TYS t3/t10 = 1.00); both fully localize the modification. *Note:* chance baselines over the 300 candidates are t1 = 1/300 ≈ 0.003, t3 = 0.01, t10 ≈ 0.033, percentile = 50.

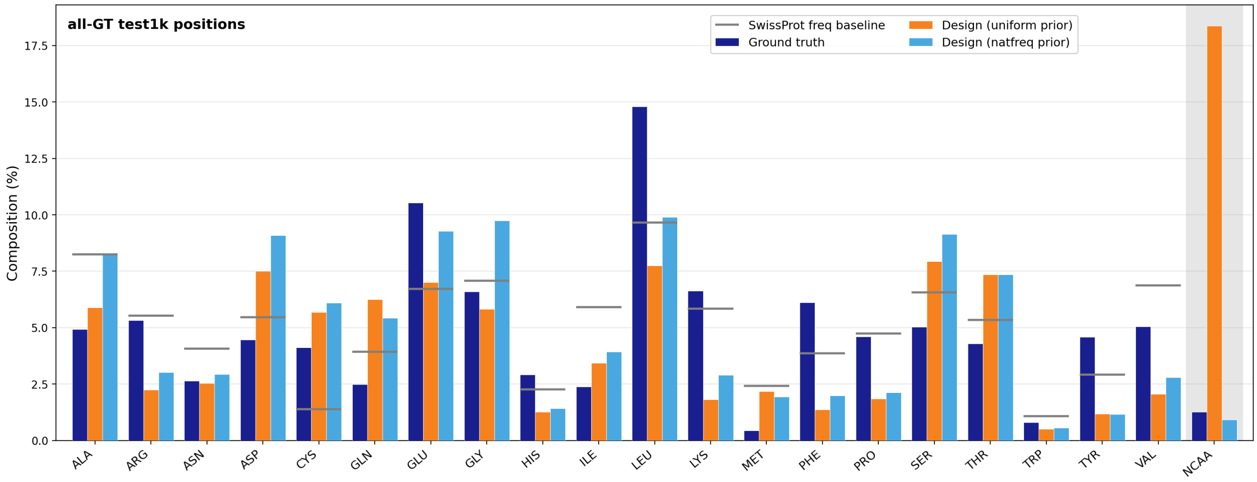


Figure S4: **Residue-type composition of the designs against ground truth** (1k held-out test set, all 199,350 positions from 10 designs each). Fraction of ground-truth positions the decode-time discretizer assigns to each residue type — the twenty canonical amino acids plus one aggregate non-canonical bin — overlaid on the ground-truth composition, shown both as the readout produces it (uniform prior, i.e. unmodified) and after multiplying in a *natural-frequency prior*: SwissProt residue frequencies over the twenty canonical classes, with a shared flat floor for the non-canonicals, applied at decode time to the same fixed readout. The readout spans the full canonical alphabet and tracks the ground-truth composition. Read without a prior it over-assigns the non-canonical bin ($\approx 18\%$ of positions, against the native rate of 1.3%); the natural-frequency prior brings this to $\approx 0.9\%$ — slightly *below* the native rate, so the readout is conservative on non-canonical calls — while shifting the canonical composition modestly toward the highest-frequency residues.

| | | | Selected backbones | | |
|---|---|---|---|---|---|
| PDB | UniProt | Protein | 8 | 16 | 24 |
| 1FGL | P62937 | Peptidyl-prolyl cis-trans isomerase A (PPIA / cyclophilin A) | 7 | 10 | 10 |
| 3JQ5 | P60045 | Acidic phospholipase A2 3 | 10 | 10 | 10 |
| 7YKH | Q9WVQ1 | MAGI2 (PDZ0–GK fragment) | 10 | 10 | 10 |
| 6YOO | Q9H0R8 | GABARAPL1 | 10 | 10 | 10 |
| 3N00 | P20393 | Nuclear receptor subfamily 1 group D member 1 (NR1D1 / Rev-erb$\alpha$) | 10 | 10 | 10 |
| 6O33 | Q13526 | PIN1 | 10 | 10 | 10 |
| 3O2M | P45983 | MAPK8 (JNK1) | 10 | 10 | 10 |
| 4RRV | O15530 | PDPK1 (PDK1) | 10 | 10 | 10 |
| 4Y7R | P61964 | WDR5 | 10 | 10 | 10 |
| 6CDG | Q8IVV7 | GID4 | 10 | 10 | 10 |
| 7OUN | Q9NZQ7 | CD274 (PD-L1) | 10 | 6 | 10 |
| 9CDT | Q07820 | MCL1 | 10 | 10 | 10 |

Table S3: **PeptideArena Target proteins and the number of selected de novo peptide backbones.** For 8-mer 1FGL and 16-mer 7OUN, the pool of binders generated by boltzgen did not produce enough distinct backbones.

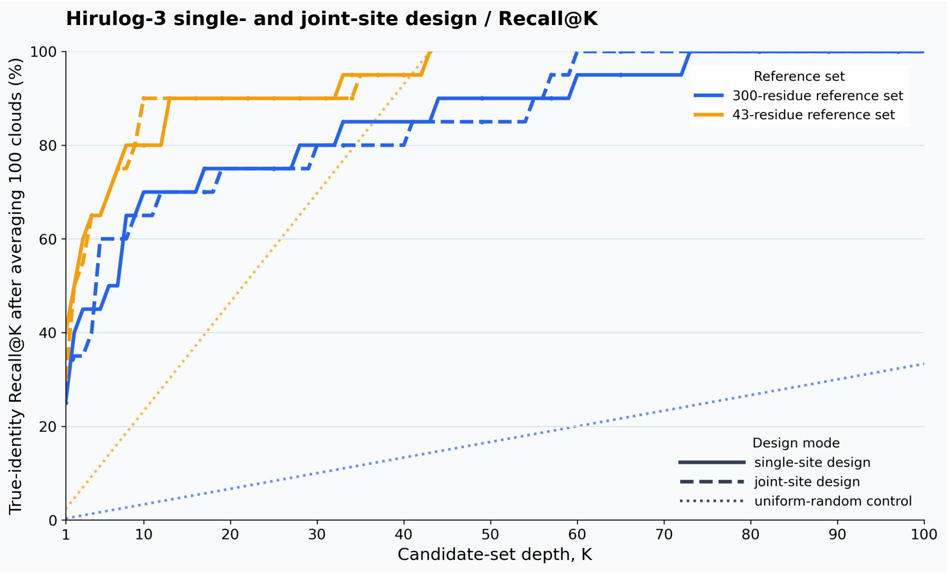


Figure S5: **Recall@K for Hirulog-3 residue recovery after averaging readout probabilities across 100 atom clouds per site.** Solid and dashed curves show single-site and joint-site design, respectively, evaluated with AtomWeaver-open (300-residue) or AtomWeaver-DBeta (43-residue) reference libraries; dotted curves indicate the corresponding uniform-random baselines.

| Peptide | Target | Sites | Mutants | Canonical | Non-canonical |
|---|---|---|---|---|---|
| PUMA BH3 | MCL1 | 27 | 1,080 | 513 | 567 |
| CP2 | KDM4A | 12 | 468 | 216 | 252 |
| Total | | 39 | 1,548 | 729 | 819 |

Table S4: **Benchmark coverage after mapping the scan onto the deposited structures.** PUMA is scanned as a 34-mer; positions outside 4–30 are absent from the 2ROC density and are dropped. Twenty-one mutants carry an independent SPR measurement used to audit the heat-map transcription (12 PUMA, 9 CP2).

DMS, and AtomWeaver-canon apply alternative learned identity readouts to the same stored clouds rather than independently sampling the generator. The DMS readout covers the 20 canonical and 21 non-canonical DMS identities. Before log averaging across draws, probabilities are floored at $10^{-12}$ and renormalized solely for numerical stability.

**Statistic, aggregation, and uncertainty.** At every rankable site (at least five candidates), we compute $-\rho_{\mathrm{Spearman}}$ between the model score and measured $\Delta\Delta G$ over the same candidates; larger positive values denote stronger agreement. Site values are averaged separately within CP2 and PUMA, and their unweighted average is reported as the equal-system mean. At both sampling temperatures, every joint-design baseline row is rankable at all 12 CP2 and 27 PUMA positions.

For the primary statistic, Supplementary Table S5 reports stratified, nonparametric site-bootstrap 90% percentile intervals. In each of 100,000 replicates, the 12 CP2 positions and 27 PUMA positions are resampled independently with replacement; the two resampled system means are then averaged with equal weight. These intervals quantify variation across the fixed benchmark positions conditional on the DMS labels and stored model draws. They are neither independent experimental-replication intervals nor estimates of draw-level Monte Carlo uncertainty. Boldface identifies the numerically highest inverse-folding value in a table column, not a significance test. NCFlow is shown as a reference-only comparator, not an eligible winner, using eight conformer draws for each candidate in the DMS candidate alphabet. The stratified bootstrap uses fixed random seed 20260921; NCFlow's upstream conformer run did not record an explicit seed.

### S18 Hirulog-3 extended data

### S19 DMS-derived expectation for WT recovery

This auxiliary analysis is distinct from the PUMA/CP2 DMS ranking benchmark in §4.5. Whereas that benchmark compares a model's within-site ranking with measured substitution effects, this section uses additional DMS datasets to estimate how often a wild-type residue is among the best measured choices under a physical objective. We treat that frequency as an empirical ceiling on wild-type recovery for a model optimizing the same objective, not as a second head-to-head inverse-folding comparison.

Two types of DMS experiment are included: assays whose readout is monomeric protein stability, and assays whose readout is binding affinity in a protein complex.

For each position, the candidate set consisted of the wild-type residue and all measured single amino acid substitutions that passed the dataset filters. Scores were oriented so that higher values were better, with wild

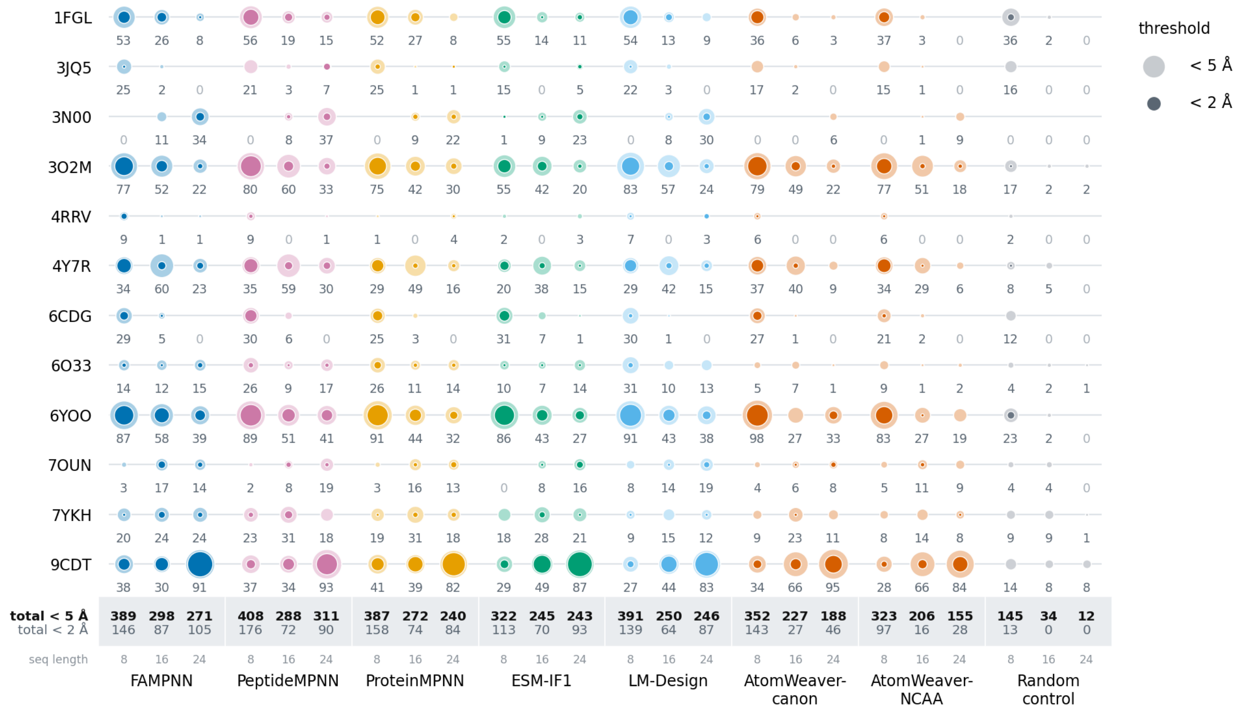


Figure S6: **PeptideArena Designs refolding under 5 Å (outer disc) and 2 Å (solid core) scRMSD, by target and peptide length.** Each cell holds one disc per peptide length, 8 / 16 / 24 from left to right. The annotation corresponds to the number of designs <5 Å scRMSD (outer disc size). Random control is a randomly sampled sequence based on natural amino acid frequency in the training set. It should be noted that for 3N00, 99.8% of the folds had a receptor RMSD > 2 Å and is therefore invalid.

type as the reference for binding assays. Candidate residues tied for the top score defined a top set of size `N`. The wild-type residue contributed `1/N` if it was in this top set and 0 otherwise. This tie-splitting rule avoids arbitrarily assigning full credit to one residue when multiple residues share the top score. The reported recovery is the mean score across scored positions or, for multi-mutant monomer analyses, across scored position-by-background contexts.

DMS positions were mapped to structures by exact numbering, constant offset, or sequence alignment, depending on the dataset. For monomers, side-chain relative solvent accessibility was computed from side-chain heavy atoms plus $C_\alpha$ and normalized by the same atoms in an isolated residue conformation. Positions were classified as buried (relSASA $< 0.05$), partially exposed ($0.05 \leq$ relSASA $< 0.25$), or exposed (relSASA $\geq 0.25$). For complexes, interface exposure was computed as the loss of side-chain solvent-accessible surface area between the isolated mutated chain and the bound complex; residues for which the partner buried at least 5 Å$^2$ of side-chain surface area were classified as interface positions, and all other residues kept the free-chain buried/partial/exposed classification. In the double-mutant monomer analysis, a scanned position and the fixed background mutation were classified as contacting when their $C_\beta$ distance was at most 7 Å, with glycine $C_\beta$ positions inferred from the backbone.

**Monomer folding-stability recovery.** For monomer folding stability, native recovery was modest overall. Across 21,323 scored positions in 382 natural domains, WT recovery was 0.273 (Table S7).

Recovery depended strongly on burial. In the native background, buried positions had recovery of 0.524, whereas exposed positions had recovery of 0.158. Thus folding stability substantially constrains core identity, but many exposed residues can be replaced without reducing the measured physical objective below wild type.

The double-mutant subset was a harder subset of the data: the matched single-mutant baseline was lower than the full single-mutant baseline (0.238 versus 0.273). Within that matched subset, scoring recovery in a one-mutation background increased native recovery to 0.325. This comparison should be read with the usual double-mutant caveat that contexts are not independent and are conditioned on the background mutation remaining folded. Within each burial stratum, local contact between the background mutation and the scanned position reduced native dominance, consistent with local epistasis.

**Interface binding recovery.** Interface binding recovery was not explained by interface geometry alone (Table S8). Protein-protein interfaces had a higher pooled recovery than protein-peptide interfaces (0.421 versus 0.125), but the clearest contrast came from the two sides of the same SARS-CoV-2 RBD/ACE2 interface. Mutating the viral RBD side, which was selected for ACE2 binding, gave high native bias (recovery 0.619). Mutating the host ACE2 side against SARS-CoV-2 spike gave much lower native bias (recovery 0.273). Because the structural interface is approximately held fixed, this

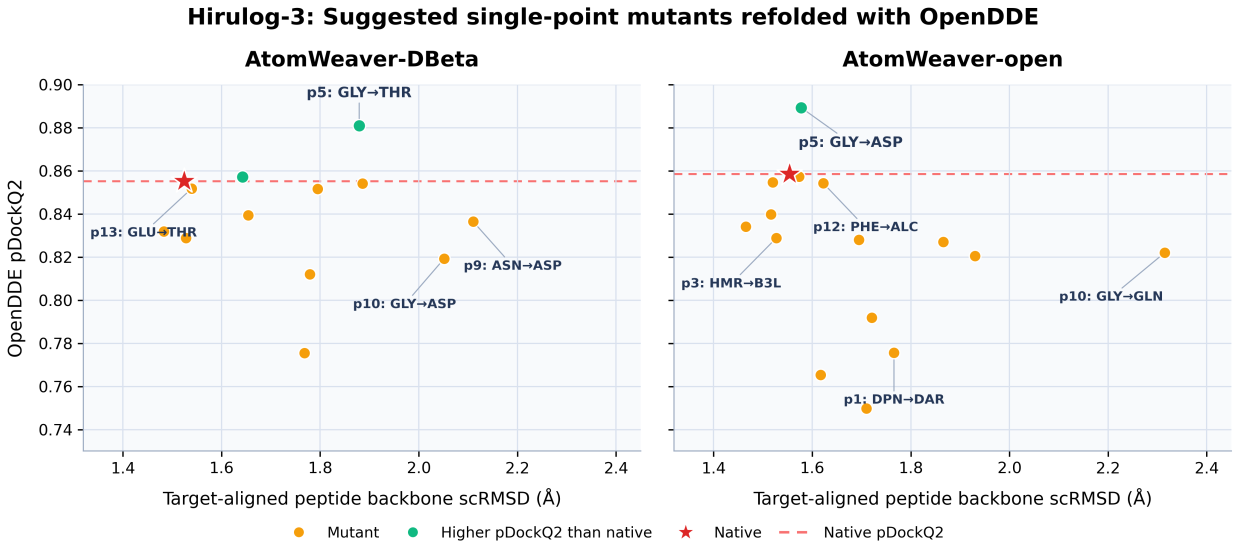


(a) OpenDDE screen of top AtomWeaver single-point suggestions. Each labeled point is refolded in complex with thrombin; dashed lines denote the corresponding corrected-native OpenDDE prediction.

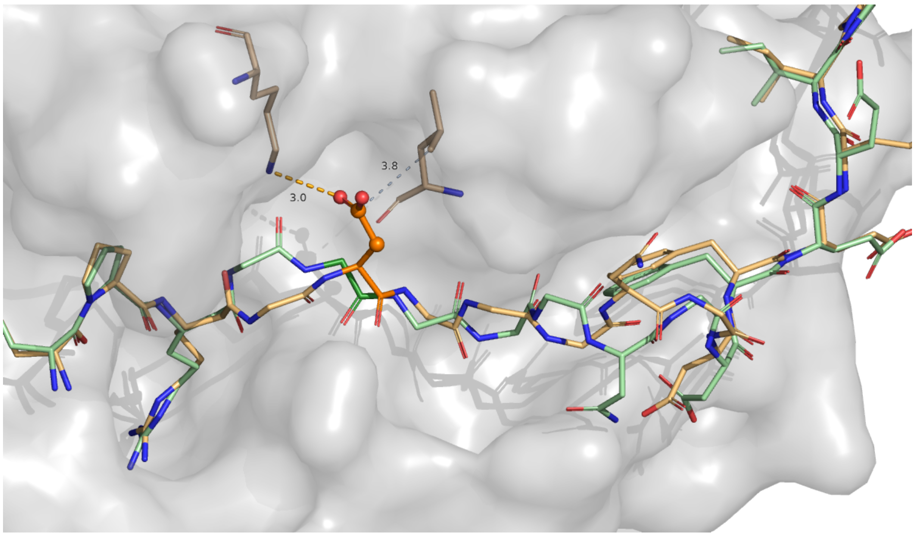


(b) Predicted thrombin-bound pose of the AtomWeaver-open Gly5→Asp candidate, superposed on the deposited Hirulog-3–thrombin complex.

Figure S7: **Prospective prioritization of AtomWeaver-suggested Hirulog-3 single-point variants.** AtomWeaver-DBeta and AtomWeaver-open use the 43- and 300-residue reference libraries, respectively. Both identify Gly5 substitutions with a compact, native-like predicted peptide pose; the AtomWeaver-open Gly5→Asp candidate shows reproducibly higher predicted complex confidence across paired OpenDDE refolds. This is a model-prioritized, structurally plausible testable analogue, not evidence of improved affinity or proteolytic stability.

| Method | Single-site design | | Joint design | |
|---|---|---|---|---|
| | CAA-only | CAA+NCAA | CAA-only | CAA+NCAA |
| AtomWeaver-open | **0.274** [0.195, 0.351] | 0.167 [0.108, 0.224] | **0.264** [0.181, 0.344] | 0.165 [0.101, 0.227] |
| AtomWeaver-DMS | 0.244 [0.167, 0.320] | **0.186** [0.135, 0.238] | 0.251 [0.166, 0.334] | **0.178** [0.123, 0.233] |
| AtomWeaver-canon | 0.229 [0.154, 0.304] | – | 0.241 [0.159, 0.319] | – |
| ProteinMPNN ($T$=0.2)‡ | – | – | 0.233 [0.122, 0.338] | – |
| ProteinMPNN ($T$=1.0) | 0.250 [0.146, 0.351] | – | 0.234 [0.124, 0.339] | – |
| PeptideMPNN ($T$=0.2)‡ | – | – | 0.244 [0.155, 0.325] | – |
| PeptideMPNN ($T$=1.0) | 0.248 [0.169, 0.320] | – | 0.249 [0.163, 0.328] | – |
| ESM-IF1 ($T$=0.2)‡ | – | – | 0.212 [0.110, 0.308] | – |
| ESM-IF1 ($T$=1.0) | 0.244 [0.143, 0.341] | – | 0.220 [0.121, 0.316] | – |
| FAMPNN ($T$=0.2)‡ | – | – | 0.218 [0.113, 0.317] | – |
| FAMPNN ($T$=1.0) | 0.243 [0.148, 0.335] | – | 0.218 [0.113, 0.318] | – |
| LM-Design ($T$=0.2)‡ | – | – | 0.205 [0.088, 0.320] | – |
| LM-Design ($T$=1.0) | 0.179 [0.062, 0.294] | – | 0.208 [0.090, 0.325] | – |
| NCFlow side-chain† | 0.071 [-0.034, 0.176] | 0.254 [0.159, 0.349] | 0.071 [-0.034, 0.176] | 0.254 [0.159, 0.349] |
| NCFlow peptide† | 0.225 [0.156, 0.295] | 0.357 [0.279, 0.430] | 0.225 [0.156, 0.295] | 0.357 [0.279, 0.430] |

Equal-system mean [90% CI]; 100,000 stratified site-bootstrap replicates.
‡$T$=0.2: joint design only. † Reference-only affinity scorer.

Table S5: **DMS Spearman agreement with 90% site-bootstrap confidence intervals.** Larger values indicate better agreement. Each of 100,000 replicates resamples the 12 CP2 and 27 PUMA sites independently, then averages the two system means with equal weight. Intervals quantify variation across benchmark sites conditional on the fixed DMS labels and stored model draws; they are not independent experimental-replication or draw-level intervals. Bold identifies the numerically highest inverse-folding point estimate in a column, not a statistically resolved difference. NCFlow is included as a reference-only affinity-scoring comparator and is not eligible for bolding.

comparison argues that evolutionary optimization for the measured partner is a major determinant of native recovery.

The remaining binding datasets support the same interpretation. GB1, a bacterial IgG-binding domain, showed relatively high interface native recovery. In contrast, PDZ3/CRIPT and GRB2-SH3/GAB2 showed low native recovery, consistent with peptide-recognition interfaces being tuned for specificity and cellular interaction context rather than maximal affinity to the assayed peptide alone.

Overall, the DMS-derived expectation for native recovery from single-position optimization of measured folding stability or binding affinity was well below unity and often far below sequence recoveries reported for machine-learning inverse-folding models. These results suggest that high model WT recovery is not explained by physical-objective optimization alone. It likely also reflects learned evolutionary sequence priors: buried residues and partner-optimized interfaces show stronger wild-type bias, whereas exposed positions, unoptimized binding partners, and specificity-tuned peptide interfaces often admit non-native amino acids that are as good as or better than wild type under the measured objective.

| objective | DMS sources | interpretation and filters |
|---|---|---|
| Monomer folding stability | Tsuboyama/Rocklin megascale stability Dataset1 [57, 58], natural monomeric domains, paired with AlphaFold [59] model structures | `DeltaG` was treated as folding stability, with higher values more stable. We required a measured wild-type stability, required wild type to be folded (`DeltaG` $\geq 0$), deduplicated domain versions to distinct PDB cores, removed variants with `DeltaG_95CI` $> 3.0$ kcal/mol, clamped stability to `[-3, 5]` kcal/mol for scoring, and scored positions with at least 15 measured substitutions. |
| Interface binding | Starr 2020 SARS-CoV-2 RBD/ACE2 [60], Chan 2020 ACE2/spike [61], Olson 2014 GB1/IgG-Fc [62], McLaughlin 2012 PSD95-PDZ3/CRIPT [63], and Faure 2022 doubledeepPCA/MoCHI outputs [64] for GRB2-SH3/GAB2, GB1/IgG-Fc, and PSD95-PDZ3/CRIPT | Binding scores were normalized so higher values indicate tighter binding, with wild type as the zero reference. Only one partner was mutated in each dataset. For Faure 2022, we used the published MoCHI-inferred binding coefficients as fold-deconvolved direct binding effects. Raw GB1 and PDZ scans were retained as checks, but their Faure-derived counterparts were used for pooled interface summaries to avoid double-counting. |

Table S6: **DMS objectives and dataset filters.**

| track | recovery | recovery buried/partial/exposed | n | domains |
|---|---|---|---|---|
| Single mutants, all domains | 0.273 | 0.524 / 0.351 / 0.158 | 21,323 | 382 |
| Single mutants, matched to double-mutant positions | 0.238 | 0.453 / 0.265 / 0.133 | 2,526 | 41 |
| Double mutants, one-mutation background | 0.325 | 0.550 / 0.412 / 0.184 | 5,012 | 41 |

Table S7: **Native recovery for monomer folding-stability tracks.**

| dataset | interface class | WT optimization for assayed partner | interface positions | recovery |
|---|---|---|---|---|
| SARS-CoV-2 RBD/ACE2 (Starr 2020) | protein-protein | optimized | 21 | 0.619 |
| ACE2/SARS-CoV-2 spike (Chan 2020) | protein-protein | not optimized | 22 | 0.273 |
| GB1/IgG-Fc, raw (Olson 2014) | protein-protein | optimized | 14 | 0.429 |
| PSD95-PDZ3/CRIPT, raw (McLaughlin 2012) | protein-peptide | not optimized / specificity-tuned | 11 | 0.273 |
| GRB2-SH3/GAB2, ddPCA (Faure 2022) | protein-peptide | not optimized / specificity-tuned | 13 | 0.077 |
| GB1/IgG-Fc, ddPCA (Faure 2022) | protein-protein | optimized | 14 | 0.357 |
| PSD95-PDZ3/CRIPT, ddPCA (Faure 2022) | protein-peptide | not optimized / specificity-tuned | 11 | 0.182 |
| Pooled protein-protein interfaces | protein-protein | mixed | 57 | 0.421 |
| Pooled protein-peptide interfaces | protein-peptide | not optimized / specificity-tuned | 24 | 0.125 |

Table S8: **WT recovery for interface binding datasets.** Pooled rows use the non-duplicated datasets: RBD/ACE2, ACE2/spike, GRB2-SH3/GAB2 ddPCA, GB1/IgG-Fc ddPCA, and PSD95-PDZ3/CRIPT ddPCA. Raw GB1 and raw PDZ are shown as independent checks but are not included in the pooled rows.

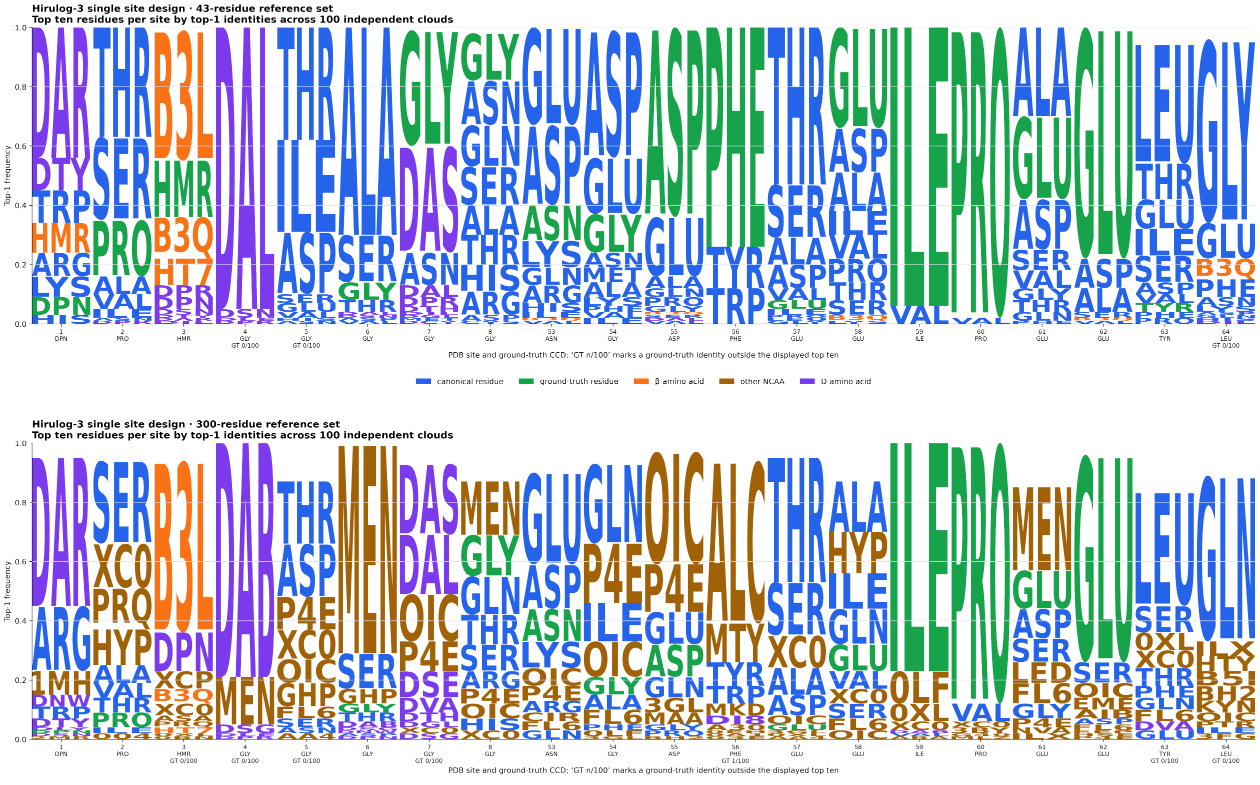


Figure S8: **Per-site top-1 identity frequencies across 100 Hirulog-3 single-site design draws. Top:** AtomWeaver-DBeta (43-residue D/$\beta$-aware reference set). **Bottom:** AtomWeaver-open (full 300-residue reference set). At each site, letters show the ten residues most frequently selected as top-1; green marks the deposited identity, blue canonical residues, purple D-amino acids, orange $\beta$-amino acids, and brown other non-canonical residues.

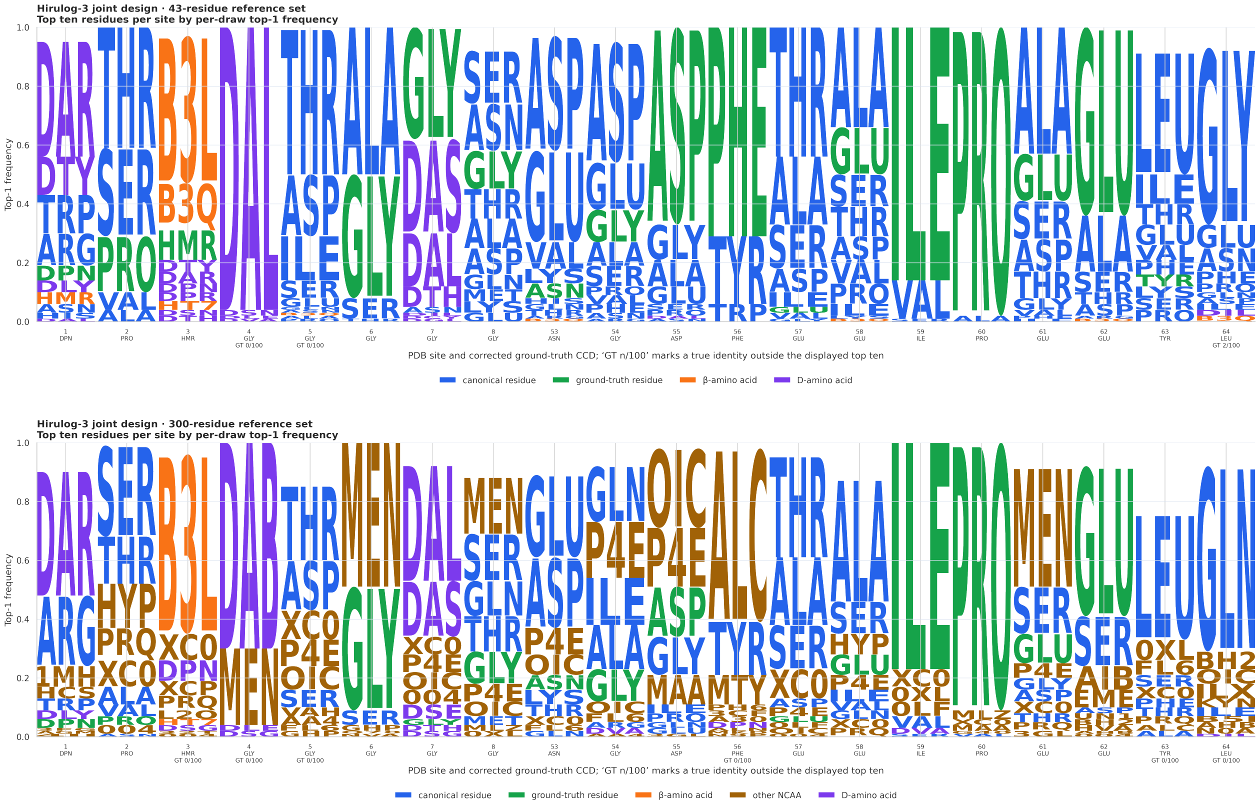


Figure S9: **Per-site top-1 identity frequencies across 100 Hirulog-3** ***joint-site*** **design draws. Top:** AtomWeaver-DBeta (43-residue D/$\beta$-aware reference set). **Bottom:** AtomWeaver-open (full 300-residue reference set). At each site, letters show the ten residues most frequently selected as top-1; green marks the deposited ground-truth identity, blue canonical residues, purple D-amino acids, orange $\beta$-amino acids, and brown other non-canonical residues.

### S20 The 300-residue reference vocabulary

Table S9 lists every residue in the decode-time reference library: the 20 canonical amino acids and the 280 non-canonical types. This is the full set a predicted cloud is matched against at decode time, and extending it is what admits a new residue without retraining (§S5.3).

| CCD | Chemistry | Cl. | Mod. | Set |
|---|---|---|---|---|
| ALA | alanine | $\alpha$ | | can. |
| ARG | arginine | $\alpha$ | | can. |
| ASN | asparagine | $\alpha$ | | can. |
| ASP | aspartate | $\alpha$ | | can. |
| CYS | cysteine | $\alpha$ | | can. |
| GLN | glutamine | $\alpha$ | | can. |
| GLU | glutamate | $\alpha$ | | can. |
| GLY | glycine | $\alpha$ | | can. |
| HIS | histidine | $\alpha$ | | can. |
| ILE | isoleucine | $\alpha$ | | can. |
| LEU | leucine | $\alpha$ | | can. |
| LYS | lysine | $\alpha$ | | can. |
| MET | methionine | $\alpha$ | | can. |
| PHE | phenylalanine | $\alpha$ | | can. |
| PRO | proline | $\alpha$ | | can. |
| SER | serine | $\alpha$ | | can. |
| THR | threonine | $\alpha$ | | can. |
| TRP | tryptophan | $\alpha$ | | can. |
| TYR | tyrosine | $\alpha$ | | can. |
| VAL | valine | $\alpha$ | | can. |
| 004 | Phe-analog, aromatic, 1 ring, 11 at. | $\alpha$ | | seen |
| 02K | analog (10 heavy atoms) | $\alpha$ | | seen |
| 02V | Phe-analog, aromatic, 1 ring, 14 at., hydroxyl | $\alpha$ | *N*Me | seen |
| 03E | analog (11 heavy atoms) | $\alpha$ | | seen |
| 0A1 | Tyr-analog, aromatic, 1 ring, 14 at. | $\alpha$ | | seen |
| 0AF | Trp-analog, bicyclic aromatic, 16 at., hydroxyl | $\alpha$ | | seen |
| 0BN | aromatic | $\alpha$ | | seen |
| 0EH | Lys-analog (13 heavy atoms) | $\alpha$ | D | seen |
| 0JY | tert-butyl-L-alanine | $\alpha$ | | seen |
| 0LF | Pro-analog, aromatic, 1 ring, 19 at. | $\alpha$ | | seen |
| 0XL | Leu-analog (9 heavy atoms) | $\alpha$ | | seen |
| 11Q | analog (15 heavy atoms) | $\alpha$ | *N*sub | seen |
| 1MH | aromatic | $\alpha$ | | seen |
| 1OP | Tyr-analog, aromatic, 1 ring, 19 at., hydroxyl | $\alpha$ | | seen |
| 1PA | aromatic, 1 ring, 16 at., carboxylate | $\alpha$ | | seen |
| 2GX | aromatic | $\alpha$ | | seen |
| 2JH | Ala-analog (10 heavy atoms) | $\alpha$ | | seen |
| 2KK | Lys-analog, aromatic, 1 ring, 19 at. | $\alpha$ | halo | seen |
| 2KY | Glu-analog (10 heavy atoms) | $\alpha$ | | seen |
| 2L5 | Phe-analog, aromatic, 1 ring, 13 at. | $\alpha$ | halo | seen |
| 2MR | sym. dimethyl-Arg | $\alpha$ | | **held** |
| 2MT | Pro-analog, thiazolidine, 10 at. | $\alpha$ | | seen |
| 2RX | Ser-analog (11 heavy atoms) | $\alpha$ | phospho | seen |
| 32T | Phe-analog, bicyclic aromatic, 14 at. | $\alpha$ | | seen |
| 3BY | Pro-analog (9 heavy atoms) | $\alpha$ | *N*Me | seen |
| 3CF | Phe-analog, aromatic, 1 ring, 14 at., nitrile | $\alpha$ | | seen |
| 3EG | Ser-analog (10 heavy atoms) | $\alpha$ | halo | seen |
| 3FG | Tyr-analog, aromatic, 1 ring, 13 at., hydroxyl | $\alpha$ | | seen |
| 3GL | analog (11 heavy atoms, carboxylate, hydroxyl) | $\alpha$ | | seen |
| 3YM | aromatic, 1 ring, 15 at., hydroxyl | $\alpha$ | | seen |
| 41H | aromatic | $\alpha$ | | seen |
| 45W | D-Pro, 11 at. | $\alpha$ | D | seen |
| 4BF | Phe-analog, aromatic, 1 ring, 13 at. | $\alpha$ | halo | seen |
| 4CF | Phe-analog, aromatic, 1 ring, 14 at., nitrile | $\alpha$ | | seen |

*continued overleaf*

*continued*

| CCD | Chemistry | Cl. | Mod. | Set |
|---|---|---|---|---|
| 4CY | Met-analog (10 heavy atoms, nitrile) | $\alpha$ | | seen |
| 4FO | Val-analog (8 heavy atoms, amine) | $\alpha$ | D | seen |
| 4FW | Trp-analog, bicyclic aromatic, 16 at. | $\alpha$ | halo | seen |
| 4II | Phe-analog, aromatic, 1 ring, 15 at., azide | $\alpha$ | | seen |
| 4J2 | Phe-analog, bicyclic aromatic, 16 at. | $\alpha$ | D | seen |
| 4LZ | Tyr-analog, aromatic, 1 ring, 18 at., amine | $\alpha$ | | seen |
| 4OG | Phe-analog, bicyclic aromatic, 15 at. | $\alpha$ | | seen |
| 4PH | Phe-analog, aromatic, 1 ring, 13 at. | $\alpha$ | | seen |
| 4PQ | Trp-analog, bicyclic aromatic, 16 at., hydroxyl | $\alpha$ | | seen |
| 4WQ | Lys-analog (13 heavy atoms) | $\alpha$ | | seen |
| 5GM | Leu-analog (11 heavy atoms) | $\alpha$ | | seen |
| 5PG | Tyr-analog, aromatic, 1 ring, 13 at., hydroxyl | $\alpha$ | *N*Me | seen |
| 5T3 | Lys-analog (15 heavy atoms, amine) | $\alpha$ | | seen |
| 66D | Leu-analog (11 heavy atoms) | $\alpha$ | | seen |
| 66E | Val-analog (10 heavy atoms) | $\alpha$ | *N*sub | seen |
| 6CW | Trp-analog, bicyclic aromatic, 16 at. | $\alpha$ | halo | seen |
| 6FL | Leu-analog (15 heavy atoms) | $\alpha$ | halo | seen |
| 6G5 | Trp-analog, bicyclic aromatic, 16 at. | $\alpha$ | halo | seen |
| 6ZS | Val-analog (8 heavy atoms) | $\alpha$ | | seen |
| 73C | Ser-analog (11 heavy atoms) | $\alpha$ | | seen |
| 73O | Tyr-analog, aromatic, 1 ring, 14 at., hydroxyl | $\alpha$ | | seen |
| 7O5 | Glu-analog (11 heavy atoms) | $\alpha$ | acyl | seen |
| 7T2 | Phe-analog, aromatic, 1 ring, 14 at. | $\alpha$ | *N*Me, halo | seen |
| 7VU | Phe-analog, aromatic, 1 ring, 15 at. | $\alpha$ | *N*sub | seen |
| 81R | Val-analog (10 heavy atoms) | $\alpha$ | | seen |
| 81S | Val-analog (10 heavy atoms) | $\alpha$ | | seen |
| 9DT | Phe-analog, aromatic, 1 ring, 14 at. | $\alpha$ | halo | seen |
| 9R1 | analog (13 heavy atoms, carboxylate) | $\alpha$ | | seen |
| 9R4 | analog (16 heavy atoms, amine, carboxylate) | $\alpha$ | acyl | seen |
| 9R7 | analog (14 heavy atoms, carboxylate) | $\alpha$ | *N*sub, acyl | seen |
| A30 | aromatic, 1 ring, 15 at. | $\alpha$ | | seen |
| AA4 | analog (9 heavy atoms, hydroxyl) | $\alpha$ | | seen |
| ABA | Ala-analog (7 heavy atoms) | $\alpha$ | | seen |
| AC5 | analog (9 heavy atoms) | $\alpha$ | | seen |
| AGM | Arg-analog (13 heavy atoms, guanidinium) | $\alpha$ | | seen |
| AHB | Asn-analog (10 heavy atoms, hydroxyl) | $\alpha$ | | seen |
| AHP | Leu-analog (10 heavy atoms) | $\alpha$ | | seen |
| AIB | $\alpha$-methyl-Ala | $\alpha$ | | seen |
| ALC | aliphatic | $\alpha$ | | seen |
| ALN | Phe-analog, bicyclic aromatic, 16 at. | $\alpha$ | | seen |
| ALY | acetyl-Lys | $\alpha$ | acyl | seen |
| API | analog (13 heavy atoms, amine, carboxylate) | $\alpha$ | | seen |
| ASA | Asp-analog | $\alpha$ | | seen |
| B3L | $\beta$-homo-Leu | $\beta$ | | seen |
| B3Q | $\beta$-homo-Gln, 11 at. | $\beta$ | | seen |
| B5I | Leu-analog (11 heavy atoms) | $\alpha$ | | seen |
| BH2 | analog (10 heavy atoms, carboxylate, hydroxyl) | $\alpha$ | | seen |
| BIF | Phe-analog, bicyclic aromatic, 18 at. | $\alpha$ | | seen |
| BMT | Thr-analog (14 heavy atoms, hydroxyl) | $\alpha$ | *N*Me | seen |
| BW5 | Ser-analog (16 heavy atoms) | $\alpha$ | | seen |
| CCS | analog (11 heavy atoms, carboxylate) | $\alpha$ | | seen |
| CGU | $\gamma$-carboxy-Glu | $\alpha$ | | **held** |
| CHG | cyclohexyl-Gly | $\alpha$ | | seen |
| CIR | citrulline | $\alpha$ | | seen |

*continued overleaf*

*continued*

| CCD | Chemistry | Cl. | Mod. | Set |
|---|---|---|---|---|
| CME | Cys-analog (11 heavy atoms, hydroxyl) | $\alpha$ | | seen |
| CSA | Cys-analog (11 heavy atoms) | $\alpha$ | | seen |
| CSO | oxidized-Cys | $\alpha$ | | seen |
| CY1 | Cys-analog (12 heavy atoms) | $\alpha$ | acyl | seen |
| D0Q | Trp-analog, bicyclic aromatic, 16 at. | $\alpha$ | | seen |
| D11 | Thr-analog (12 heavy atoms) | $\alpha$ | phospho, D | seen |
| D4P | Tyr-analog, aromatic, 1 ring, 12 at., hydroxyl | $\alpha$ | | seen |
| DA2 | Arg-analog (14 heavy atoms, guanidinium) | $\alpha$ | | seen |
| DAB | Val-analog (8 heavy atoms, amine) | $\alpha$ | | seen |
| DAL | D-Ala | $\alpha$ | D | seen |
| DAR | Arg-analog (12 heavy atoms, guanidinium) | $\alpha$ | D | seen |
| DAS | analog (9 heavy atoms, carboxylate) | $\alpha$ | D | seen |
| DBZ | Ala-analog, aromatic, 1 ring, 15 at. | $\alpha$ | acyl | seen |
| DC0 | Phe-analog, aromatic, 1 ring, 19 at., amine | $\alpha$ | *N*sub | seen |
| DCY | D-Cys, 7 at. | $\alpha$ | D | seen |
| DGL | D-Glu | $\alpha$ | D | seen |
| DGN | D-Gln, 10 at. | $\alpha$ | D | seen |
| DHI | His-analog, aromatic, 1 ring, 11 at. | $\alpha$ | D | seen |
| DI7 | Tyr-analog, aromatic, 1 ring, 15 at., hydroxyl | $\alpha$ | | seen |
| DI8 | Pro-analog, aromatic, 1 ring, 13 at. | $\alpha$ | | seen |
| DIL | D-Ile, 9 at. | $\alpha$ | D | seen |
| DLE | D-Leu, 9 at. | $\alpha$ | D | seen |
| DLY | D-Lys, 10 at., amine | $\alpha$ | D | seen |
| DMK | analog (11 heavy atoms, carboxylate) | $\alpha$ | | seen |
| DNE | D-norleucine, 9 at. | $\alpha$ | D | seen |
| DNW | Ala-analog, aromatic, 1 ring, 19 at., amine | $\alpha$ | | seen |
| DPN | D-Phe | $\alpha$ | D | seen |
| DPP | analog (7 heavy atoms, amine) | $\alpha$ | | seen |
| DPR | D-Pro | $\alpha$ | D | seen |
| DSE | D-Ser, 8 at., hydroxyl | $\alpha$ | D, *N*Me | seen |
| DSG | D-Asn, 9 at. | $\alpha$ | D | seen |
| DSN | D-Ser, 7 at., hydroxyl | $\alpha$ | D | seen |
| DTH | D-Thr, 8 at., hydroxyl | $\alpha$ | D | seen |
| DTR | D-Trp-analog, bicyclic aromatic, 15 at. | $\alpha$ | D | seen |
| DTY | D-Tyr | $\alpha$ | D | seen |
| DVA | D-Val, 8 at. | $\alpha$ | D | seen |
| E95 | Trp-analog, bicyclic aromatic, 16 at. | $\alpha$ | | seen |
| E9M | Trp-analog, bicyclic aromatic, 16 at. | $\alpha$ | *N*Me | seen |
| E9V | His-analog, aromatic, 1 ring, 12 at. | $\alpha$ | *N*Me | seen |
| EEP | Ser-analog (11 heavy atoms, hydroxyl) | $\alpha$ | | seen |
| ELY | Lys-analog (14 heavy atoms, amine) | $\alpha$ | | seen |
| EME | analog (11 heavy atoms, carboxylate) | $\alpha$ | *N*Me | seen |
| EU0 | Val-analog (11 heavy atoms, guanidinium) | $\alpha$ | *N*sub | seen |
| FAK | Lys-analog (16 heavy atoms) | $\alpha$ | halo | seen |
| FCL | Phe-analog, aromatic, 1 ring, 13 at. | $\alpha$ | halo | seen |
| FL6 | Asp-analog (10 heavy atoms) | $\alpha$ | | seen |
| FLA | Ala-analog (9 heavy atoms) | $\alpha$ | halo | seen |
| FP9 | fluoro-Pro | $\alpha$ | halo | seen |
| FTR | Trp-analog, bicyclic aromatic, 16 at. | $\alpha$ | halo | seen |
| FVA | Val-analog (10 heavy atoms) | $\alpha$ | *N*sub, acyl | seen |
| FX9 | Phe-analog, aromatic, 1 ring, 16 at. | $\alpha$ | halo | seen |
| FXC | Phe-analog, aromatic, 1 ring, 16 at. | $\alpha$ | halo, D | seen |
| GHP | Tyr-analog, aromatic, 1 ring, 12 at., hydroxyl | $\alpha$ | D | seen |
| GNC | Gln-analog (11 heavy atoms) | $\alpha$ | *N*Me | seen |
| H14 | Phe-analog, aromatic, 1 ring, 13 at., hydroxyl | $\alpha$ | | seen |

*continued overleaf*

*continued*

| CCD | Chemistry | Cl. | Mod. | Set |
|---|---|---|---|---|
| HAQ | Pro-analog, aromatic, 1 ring, 18 at., amine | $\alpha$ | acyl | seen |
| HCS | Cys-analog (8 heavy atoms) | $\alpha$ | | seen |
| HIC | His-analog, aromatic, 1 ring, 12 at. | $\alpha$ | | seen |
| HIX | aromatic | $\alpha$ | | seen |
| HJY | Phe-analog, aromatic, 1 ring, 14 at. | $\alpha$ | *N*Me, halo, D | seen |
| HLX | homo-Leu | $\alpha$ | | seen |
| HMR | $\beta$-homo-Arg | $\beta$ | | seen |
| HPE | Phe-analog, aromatic, 1 ring, 13 at. | $\alpha$ | | seen |
| HRG | Arg-analog (13 heavy atoms, guanidinium) | $\alpha$ | | seen |
| HRP | Trp-analog, bicyclic aromatic, 16 at., hydroxyl | $\alpha$ | | seen |
| HSE | Ser-analog (8 heavy atoms, hydroxyl) | $\alpha$ | | seen |
| HT7 | $\beta$-homo-Trp, aromatic, indole | $\beta$ | | seen |
| HTR | Trp-analog, bicyclic aromatic, 16 at., hydroxyl | $\alpha$ | | seen |
| HTY | Tyr-analog, aromatic, 1 ring, 14 at., hydroxyl | $\alpha$ | D | seen |
| HVA | Val-analog (9 heavy atoms, hydroxyl) | $\alpha$ | | seen |
| HYP | 4-hydroxy-Pro | $\alpha$ | | seen |
| I2M | Val-analog (10 heavy atoms) | $\alpha$ | | seen |
| IAE | Phe-analog, aromatic, 1 ring, 17 at. | $\alpha$ | halo | seen |
| IGL | Phe-analog, aromatic, 1 ring, 14 at. | $\alpha$ | | seen |
| ILX | Ile-analog (11 heavy atoms, hydroxyl) | $\alpha$ | | seen |
| IML | Ile-analog (10 heavy atoms) | $\alpha$ | *N*Me | seen |
| IYR | iodo-Tyr | $\alpha$ | halo | seen |
| KCJ | Phe-analog, aromatic, 1 ring, 11 at. | $\alpha$ | | seen |
| KCR | $N^6$-crotonyl-Lys | $\alpha$ | acyl | **held** |
| KHB | Lys-analog (16 heavy atoms, hydroxyl) | $\alpha$ | acyl | seen |
| KYN | Phe-analog, aromatic, 1 ring, 15 at., amine | $\alpha$ | | seen |
| L2O | $\beta$-homo-Leu, 11 at., hydroxyl | $\beta$ | | seen |
| LAV | Val-analog (15 heavy atoms, amine) | $\alpha$ | *N*sub | seen |
| LBZ | Lys-analog, aromatic, 1 ring, 18 at. | $\alpha$ | acyl | seen |
| LE1 | aliphatic | $\alpha$ | | seen |
| LED | Leu-analog (10 heavy atoms) | $\alpha$ | | seen |
| LMF | Lys-analog (14 heavy atoms, amine) | $\alpha$ | | seen |
| LWI | Phe-analog, aromatic, 1 ring, 14 at., amine | $\alpha$ | | seen |
| LYZ | Lys-analog (11 heavy atoms, amine, hydroxyl) | $\alpha$ | | seen |
| M2L | Met-analog (12 heavy atoms, amine) | $\alpha$ | | seen |
| M3L | trimethyl-Lys | $\alpha$ | | seen |
| MAA | N-methyl-Ala | $\alpha$ | *N*Me | seen |
| MEA | N-methyl-Phe | $\alpha$ | *N*Me | seen |
| MED | D-Met, 9 at. | $\alpha$ | D | seen |
| MEN | Asn-analog (10 heavy atoms) | $\alpha$ | acyl | seen |
| MF3 | Met-analog (12 heavy atoms) | $\alpha$ | halo | seen |
| MHS | His-analog, aromatic, 1 ring, 12 at. | $\alpha$ | | seen |
| MK8 | $\alpha$-Me-norleucine | $\alpha$ | | **held** |
| MKD | Ile-analog (12 heavy atoms) | $\alpha$ | | seen |
| ML3 | Met-analog (13 heavy atoms, amine) | $\alpha$ | | seen |
| MLE | N-methyl-Leu | $\alpha$ | *N*Me | seen |
| MLU | D-Leu, 10 at. | $\alpha$ | D, *N*Me | seen |
| MLY | dimethyl-Lys | $\alpha$ | | seen |
| MLZ | Lys-analog (11 heavy atoms, amine) | $\alpha$ | | seen |
| MP8 | 4-methyl-Pro | $\alpha$ | | **held** |
| MPQ | Gly-analog, aromatic, 1 ring, 12 at. | $\alpha$ | *N*Me | seen |
| MTY | aromatic, 1 ring, 13 at., hydroxyl | $\alpha$ | | seen |
| MV9 | D-Val, 9 at. | $\alpha$ | D, *N*Me | seen |
| MVA | N-methyl-Val | $\alpha$ | *N*Me | seen |
| MYN | Arg-analog (12 heavy atoms, guanidinium) | $\alpha$ | | seen |

*continued overleaf*

*continued*

| CCD | Chemistry | Cl. | Mod. | Set |
|---|---|---|---|---|
| N0A | Phe-analog, aromatic, 1 ring, 13 at. | $\alpha$ | halo | seen |
| N2C | Cys-analog (9 heavy atoms) | $\alpha$ | $N$Me | seen |
| N7P | Pro-analog (11 heavy atoms) | $\alpha$ | $N$sub, acyl | seen |
| N9P | Phe-analog, aromatic, 1 ring, 12 at. | $\alpha$ | | seen |
| NAL | naphthyl-Ala | $\alpha$ | | seen |
| NCY | Cys-analog (8 heavy atoms) | $\alpha$ | $N$Me | seen |
| NIY | Tyr-analog, aromatic, 1 ring, 16 at., amine, hydroxyl | $\alpha$ | | seen |
| NLE | norleucine | $\alpha$ | | seen |
| NMC | analog (9 heavy atoms) | $\alpha$ | $N$sub | seen |
| NMM | methyl-Arg | $\alpha$ | | seen |
| NRG | Arg-analog (15 heavy atoms, guanidinium) | $\alpha$ | | seen |
| NVA | norvaline | $\alpha$ | | seen |
| NZC | Thr-analog (9 heavy atoms, hydroxyl) | $\alpha$ | $N$Me | seen |
| OBF | aliphatic (9 heavy atoms) | $\alpha$ | halo | seen |
| OCS | Cys-analog (10 heavy atoms) | $\alpha$ | sulfo | seen |
| OCY | Cys-analog (10 heavy atoms, hydroxyl) | $\alpha$ | | seen |
| OGC | His-analog, aromatic, 1 ring, 13 at., hydroxyl | $\alpha$ | | seen |
| OIC | Pro-analog (12 heavy atoms) | $\alpha$ | | seen |
| OLT | Thr-analog (9 heavy atoms) | $\alpha$ | | seen |
| OMX | Tyr-analog, aromatic, 1 ring, 14 at., hydroxyl | $\alpha$ | | seen |
| OMY | chloro-Tyr | $\alpha$ | halo | **held** |
| ORD | Lys-analog (9 heavy atoms, amine) | $\alpha$ | D | seen |
| ORN | ornithine | $\alpha$ | | seen |
| ORQ | Lys-analog | $\alpha$ | acyl | seen |
| P4E | Phe-analog, aromatic, 1 ring, 14 at. | $\alpha$ | | seen |
| PCA | pyroglutamate | $\alpha$ | acyl | seen |
| PF5 | Phe-analog, aromatic, 1 ring, 17 at. | $\alpha$ | halo | seen |
| PFF | Phe-analog, aromatic, 1 ring, 13 at. | $\alpha$ | halo | seen |
| PH6 | Pro-analog (14 heavy atoms) | $\alpha$ | | seen |
| PH8 | Val-analog, aromatic, 1 ring, 14 at. | $\alpha$ | | seen |
| PHI | Phe-analog, aromatic, 1 ring, 13 at. | $\alpha$ | halo | seen |
| PM3 | Phe-analog, aromatic, 1 ring, 17 at. | $\alpha$ | phospho | seen |
| PPN | Phe-analog, aromatic, 1 ring, 15 at., amine | $\alpha$ | | seen |
| PRK | Lys-analog (14 heavy atoms) | $\alpha$ | acyl | seen |
| PRQ | Phe-analog, aromatic, 1 ring, 15 at., amine | $\beta$ | | seen |
| PRS | aliphatic | $\alpha$ | | seen |
| PRV | Phe-analog, aromatic, 1 ring, 14 at., amine | $\alpha$ | | seen |
| PTR | phospho-Tyr | $\alpha$ | phospho | seen |
| Q78 | Phe-analog, aromatic, 1 ring, 11 at. | $\alpha$ | | seen |
| SAC | Ser-analog (10 heavy atoms, hydroxyl) | $\alpha$ | acyl | seen |
| SAR | N-methyl-Gly | $\alpha$ | $N$Me | seen |
| SC2 | aliphatic | $\alpha$ | acyl | seen |
| SEM | Ser-analog, aromatic, 1 ring, 14 at. | $\alpha$ | | seen |
| SEP | phospho-Ser | $\alpha$ | phospho | seen |
| SLL | Ala-analog (17 heavy atoms, carboxylate) | $\alpha$ | acyl | seen |
| SMC | Cys-analog (8 heavy atoms) | $\alpha$ | | seen |
| SMF | Phe-analog, aromatic, 1 ring, 17 at. | $\alpha$ | sulfo | seen |
| SNC | Cys-analog (9 heavy atoms) | $\alpha$ | | seen |
| TBG | Val-analog (9 heavy atoms) | $\alpha$ | | seen |
| TH5 | Thr-analog (11 heavy atoms) | $\alpha$ | | seen |
| TIG | Trp-analog, bicyclic aromatic, 18 at., amine | $\alpha$ | $N$sub | seen |
| TIH | Phe-analog, aromatic, 1 ring, 11 at. | $\alpha$ | | seen |
| TPO | phospho-Thr | $\alpha$ | phospho | seen |
| TRF | Trp-analog, bicyclic aromatic, 17 at. | $\alpha$ | | seen |
| TRN | Phe-analog, bicyclic aromatic, 15 at. | $\alpha$ | | seen |

*continued overleaf*

*continued*

| CCD | Chemistry | Cl. | Mod. | Set |
|---|---|---|---|---|
| TRX | Trp-analog, bicyclic aromatic, 16 at., hydroxyl | $\alpha$ | | seen |
| TVA | Val-analog, aromatic, 1 ring, 19 at., amine, hydroxyl | $\alpha$ | $N$sub | seen |
| TYI | Tyr-analog, aromatic, 1 ring, 15 at., hydroxyl | $\alpha$ | halo | seen |
| TYS | sulfo-Tyr | $\alpha$ | sulfo | seen |
| U2M | Lys-analog (11 heavy atoms) | $\alpha$ | | seen |
| UKD | Ala-analog, aromatic, 1 ring, 15 at. | $\alpha$ | phospho | seen |
| UKY | Ala-analog, aromatic, 1 ring, 15 at. | $\alpha$ | phospho | seen |
| UN1 | analog (11 heavy atoms, carboxylate) | $\alpha$ | | seen |
| UX8 | Trp-analog, bicyclic aromatic, 16 at., hydroxyl | $\alpha$ | | seen |
| V7T | D-Lys, 13 at., guanidinium | $\alpha$ | D, $N$sub | seen |
| VAH | Thr-analog (9 heavy atoms, hydroxyl) | $\alpha$ | | seen |
| VR0 | Arg-analog (16 heavy atoms, guanidinium) | $\alpha$ | | seen |
| WLU | Leu-analog (11 heavy atoms, hydroxyl) | $\alpha$ | $N$Me | seen |
| WPA | Phe-analog, aromatic, 1 ring, 14 at. | $\alpha$ | | seen |
| WVL | Leu-analog (11 heavy atoms) | $\alpha$ | | seen |
| XA6 | Phe-analog, aromatic, 1 ring, 15 at. | $\alpha$ | | seen |
| XC0 | Leu-analog (11 heavy atoms) | $\alpha$ | | seen |
| XCP | $\beta$-homo-analog, 9 at. | $\beta$ | | seen |
| XYC | Ala-analog (11 heavy atoms) | $\alpha$ | | seen |
| YNM | aromatic | $\alpha$ | $N$Me | seen |
| Z36† | O-methyl-L-homoserine | $\alpha$ | | seen |
| Z37† | N-methyl-L-2-aminobutyric acid | $\alpha$ | $N$Me | seen |
| Z3E | Thr-analog, aromatic, 1 ring, 15 at. | $\alpha$ | | seen |
| ZCL | Phe-analog, aromatic, 1 ring, 14 at. | $\alpha$ | halo | seen |
| ZIQ | Trp-analog, bicyclic aromatic, 16 at. | $\alpha$ | | seen |
| ZNY | Pro-analog (11 heavy atoms) | $\alpha$ | | seen |
| ZU0 | Thr-analog (12 heavy atoms) | $\alpha$ | | seen |

Table S9: **The 300-residue reference vocabulary.** Every residue the decode-time readout can return: the 20 canonical amino acids and 280 non-canonical types. *Cl.* is the backbone class, measured from each reference structure as the amino-to-carboxyl path length rather than read from a catalogued label; all 300 are $\alpha$ or $\beta$ and none is a $\gamma$-amino acid. Note that this describes the *backbone* only: a residue may carry substituents at its own $\beta$, $\gamma$ or $\delta$ positions while still having an $\alpha$ backbone, so $\gamma$-carboxyglutamate (CGU), for instance, is listed here correctly as $\alpha$. *Mod.* records modifications detected from structure — D configuration, backbone-nitrogen substitution ($N$Me for a methyl, $N$sub for a bulkier group), and post-translational chemistry identified by element and moiety (phospho, sulfo, halo, acyl). A blank means none of these was detected, not that the residue is unmodified in every sense. *Set* distinguishes the canonical twenty, the 274 non-canonical types seen in training, and the 6 zero-shot holdouts withheld from the generative model and carried by reference geometry alone. This set is a default rather than a limit: because identity is decided at decode time by matching against these references, a user can extend the vocabulary by supplying a reference structure for a residue not listed here, with no retraining of the generative model. We are also working toward a larger default vocabulary with high-quality rotamers. †Z36 and Z37 are ProteinQure identifiers rather than CCD codes: neither residue has an equivalent entry in the Chemical Component Dictionary.
